# Animal tracking with particle algorithms for conservation

**DOI:** 10.1101/2025.02.13.638042

**Authors:** Edward Lavender, Andreas Scheidegger, Carlo Albert, Stanisław W. Biber, Jakob Brodersen, Dmitry Aleynik, Georgina Cole, Jane Dodd, Peter J. Wright, Janine Illian, Mark James, Sophie Smout, James Thorburn, Helen Moor

## Abstract

The movements of aquatic animals affect their exposure to threats and the efficacy of conservation measures, such as Marine Protected Areas (MPAs). However, many species’ movements remain poorly understood and difficult to reconstruct from available datasets, hampering conservation efforts. This is especially the case for species that rarely surface, for which data are often limited to observations from acoustic telemetry (detections) and ancillary sensors, such as archival tags. Here, we pioneer the use of state-of-the-art particle algorithms to model animal movement, integrate datasets and assess MPA design, using a case study of the Critically Endangered flapper skate (*Dipturus intermedius*) in Scotland. Our algorithms led to 5-fold improvements in maps of space use and 30-fold improvements in residency estimates (lower mean error) compared to prevailing heuristic methods. By formally integrating tracking datasets, we were uniquely able to examine movements beyond receivers into fished zones, MPA-scale residency and specific habitats beyond protected areas that may warrant protection. This work showcases a probabilistically sound modelling framework that is sufficiently fast, flexible and accessible to meet the demands of modern animal-tracking datasets in acoustic telemetry systems. This represents a marked advance for analyses of animal movements and MPA efficacy worldwide.

## 1. Introduction

Ocean biodiversity is increasingly threatened by anthropogenic activities, such as overfishing^1^. Since 1970, the Marine Living Planet Index has fallen by 56 %^2^. Many marine taxa have declined, especially in coastal ecosystems^3^. In some taxa, such as elasmobranchs (sharks, skates and rays), rates of decline are now critical^4,5^. There is a pressing need for research designed to inform conservation measures that can support these species and bend the curve of aquatic biodiversity decline^6^.

Marine Protected Areas (MPAs) are an important conservation solution^7^. An effective MPA is a refuge that locally reduces the pressures to which individuals are exposed (especially fishing) and supports population recovery^8^. Designing MPAs for mobile species requires an understanding of animal movement, which shapes individual exposure to threats, hotspots of habitat use and residency in selected areas^9^. This requirement has motivated huge interest in animal electronic tagging and tracking^10^.

Tagging and tracking technologies for aquatic species have proliferated in recent years^11^. For marine mammals and seabirds, satellite transmitters are widely used^12,13^. These tags periodically collect/transmit location data from which movement trajectories can be reconstructed using well-established statistical approaches^14^. However, for species that rarely surface, satellite tracking is limited and alternative technologies are required^11^. Passive acoustic telemetry systems are extensively deployed^15^. These use receiver arrays to detect individual-specific acoustic transmissions of tagged animals when they move within range. Since array coverage is often limited, detections are usually sparse and may be considerably enhanced by ancillary datasets, such as archival (e.g. depth) observations^16^. However, integrating sparse detections with ancillary datasets to reconstruct movement patterns (within and beyond receiver arrays) remains a substantial challenge that has significantly hampered the use of these data to inform MPA design.

Heuristic methods currently dominate efforts to analyse movements in passive acoustic telemetry systems^17–19^. These methods use detections and apply summary statistics, tuning parameters and other heuristics to map space use around receivers^17^. For example, the centre-of-activity (COA) algorithm computes weighted averages of the receiver locations where detections were recorded over sequential time intervals (of duration Δ*T*)^20,21^. Similarly, the Refined Shortest Path (RSP) algorithm interpolates ‘relocations’ along the shortest paths between the receivers that recorded sequential detections, assuming distance-dependent interpolation errors (in line with a tuning parameter termed e>r>.ad>)^22^. Post-hoc smoothing is used to generate maps of space use or utilisation distributions (UDs). Residency indices (such as ‘detection days’ or the proportion of days with detections) have also been developed to quantify residency around receivers^23^. These methods have been subject to limited formal evaluation, but their limitations in sparse receiver arrays (where individual movements are uncertain) are acknowledged^17–19^.

Recent developments in state-space modelling create major opportunities to move beyond heuristics and strengthen animal-tracking analyses for conservation^19,24^. In an animal-tracking context, a state-space model is a hierarchical framework that models an underlying movement process and the observation processes that connect movements to observations^25^. Until recently, state-space modelling routines for passive acoustic telemetry data were bespoke, computationally expensive and limited to detection data^24,26,27^. However, it is now possible to fit state-space models that integrate detections and diverse ancillary datasets (from sensor measurements to mark-recapture events) using particle filtering–smoothing algorithms^19,28^. As examples, this paper considers algorithms that incorporate acoustic observations, depth observations, or both sets of observations simultaneously, i.e., the acoustic-container (AC), depth-contour (DC) and acoustic-container depth-contour (ACDC) algorithms^19^. These algorithms represent an individual’s possible location probabilistically with a set of weighted samples, termed ‘particles’. A simulation study showed that particle algorithms consistently outperform heuristic methods, generating refined maps of space use and residency estimates for entire regions of interest, such as MPAs^19^. However, particle algorithms have yet to be exploited to inform MPA design in any real-world system.

Here, we pioneer the application of particle algorithms for conservation with a case study of the Critically Endangered flapper skate (*Dipturus intermedius*) (Figure 1). This is a large-bodied, largely benthic species that occupies habitats from 0–1200 m deep in north-western Europe^29^. Once decimated by overfishing^30^, flapper skate remain vulnerable as bycatch^31^. In 2014, the Loch Sunart to the Sound of Jura MPA was designated for flapper skate in Scotland and acoustic/archival tagging were later undertaken for monitoring^32–34^. Preliminary analyses demonstrated that skate exhibit localised movements within the MPA^32,35,36^. However, key conservation questions pertaining to MPA efficacy—including the use of seasonally fished zones beyond receivers, the extent to which skate remain in the MPA in detection gaps and the suitability of MPA boundaries—remained poorly addressed, given incomplete receiver coverage. Movements beyond the MPA, and connectivity to the only known egg nursery (in the Red Rocks and Longay MPA), also remained poorly understood^37^. Such knowledge gaps are pervasive among studies reliant upon sparse receiver arrays^38,39^ and highlight the need for statistical tools that can enhance the conservation of mobile aquatic species.

**Figure 1.**
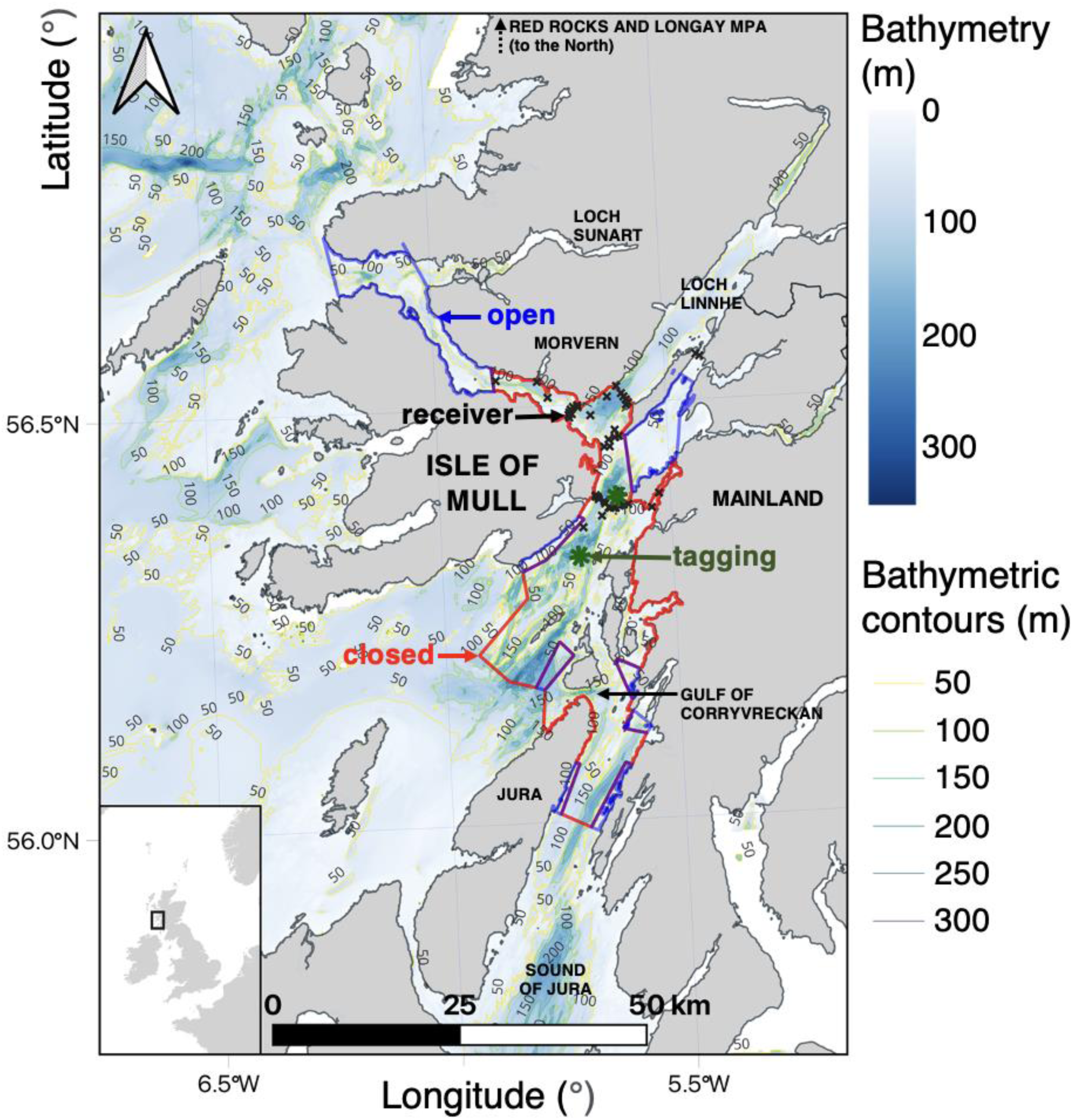
A case-study system. The inset shows the location of the study system in the United Kingdom. The main panel shows study area. The coloured polygons mark the boundary of the Loch Sunart to the Sound of Jura MPA. This includes zones that are open (blue) and closed (red) to fisheries. Tagging locations (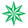), receivers (**x**) and bathymetric contours are marked. For spatial data sources, see Supporting Information §1 and Table S1.

Combining simulations and real-world analyses, this study showcases how particle algorithms can reveal marked improvements in maps of space use, residency in regions beyond receivers, MPA suitability and specific habitats beyond MPA boundaries where additional protection may be warranted. By comparing maps based on acoustic and/or archival data, we also quantify the contributions of different datasets, providing valuable information for monitoring programmes. This work provides a basis to strengthen the use of movement data for mobile species conservation across the globe.

## 2. Results

### 2.1. Overview

We conducted simulation-based and real-world analyses of animal movements in relation to an MPA using two heuristic algorithms (COAs and RSPs) and three particle filtering–smoothing algorithms (AC, DC and ACDC) (Figures 1 and S1). In simulation analyses (labelled A1 and A2), we evaluated algorithm performance (A1) and sensitivity (A2). In real-world analyses (A3 and A4), we modelled electronic tagging and tracking data from flapper skate (Figure S2). Table S2 provides the overview and a labelling hierarchy for analyses (A1, A1.1, etc.).

### 2.2. Simulation analysis

#### 2.2.1. Performance analyses

In simulation analyses of algorithm performance (A1), particle algorithms reproduced the utilisation distributions (UDs) and residency patterns exhibited by 100 simulated paths with impressive accuracy and precision, while the performance of prevailing heuristic methods varied. Heuristic algorithms were calibrated for the analysis by quantifying the mean absolute error (ME) between ‘true’ and reconstructed UDs across a range of candidate parameter values (A1.1.1). For the COA algorithm an optimal setting of Δ*T* = 2 days was identified (Figure S3). For RSPs, ME weakly declined with increasing distance-error (e>r>.ad>) values but there was no winning choice of e>r>.ad> on average. We selected e>r>.ad> = 500 m as an ‘optimal’ compromise between ME and implementation success (Figure S3). Particle algorithms were parameterised following the data-generating movement and observation processes used to simulate paths, based on prior research, domain knowledge and a literature review (see Methods). All algorithms were successfully implemented (A1.2) for all simulated paths (Figure S4).

In a visual analysis of UDs for a subset (1–3) of simulated paths (A1.3.1), we found that heuristic UDs were moderately accurate but surpassed by those from particle algorithms (Figures S5–6 versus S7). Both heuristic algorithms effectively reconstructed the extent of movements for Path 3 (for which movements were localised around receivers), but underestimated those for Paths 1 and 2 (which included movements to different areas) (Figures S5–6). Hotspot placement was partially correct (Figures S5–6). Particle algorithms represented simulated patterns more accurately (Figure S7).

Across all 100 simulated paths, a clear ranking of algorithms emerged from analysis (A1.3.2) of the ME in UD estimation (Figure 2A). All algorithms outperformed the null model on average. Performance of the two heuristic algorithms was similar and varied substantially across repeated realisations of the same data-generating-processes (occasionally overlapping with the null model). Particle algorithms consistently outperformed the heuristic algorithms— with median MEs, and standard deviations in ME, 2.7–5.0 and 1.7–3.6 times lower, respectively. For particle algorithms, the ranking was DC, AC and ACDC (best). Compared to the AC and DC algorithms, median ME for the ACDC algorithm was 1.6–1.8 times lower.

**Figure 2.**
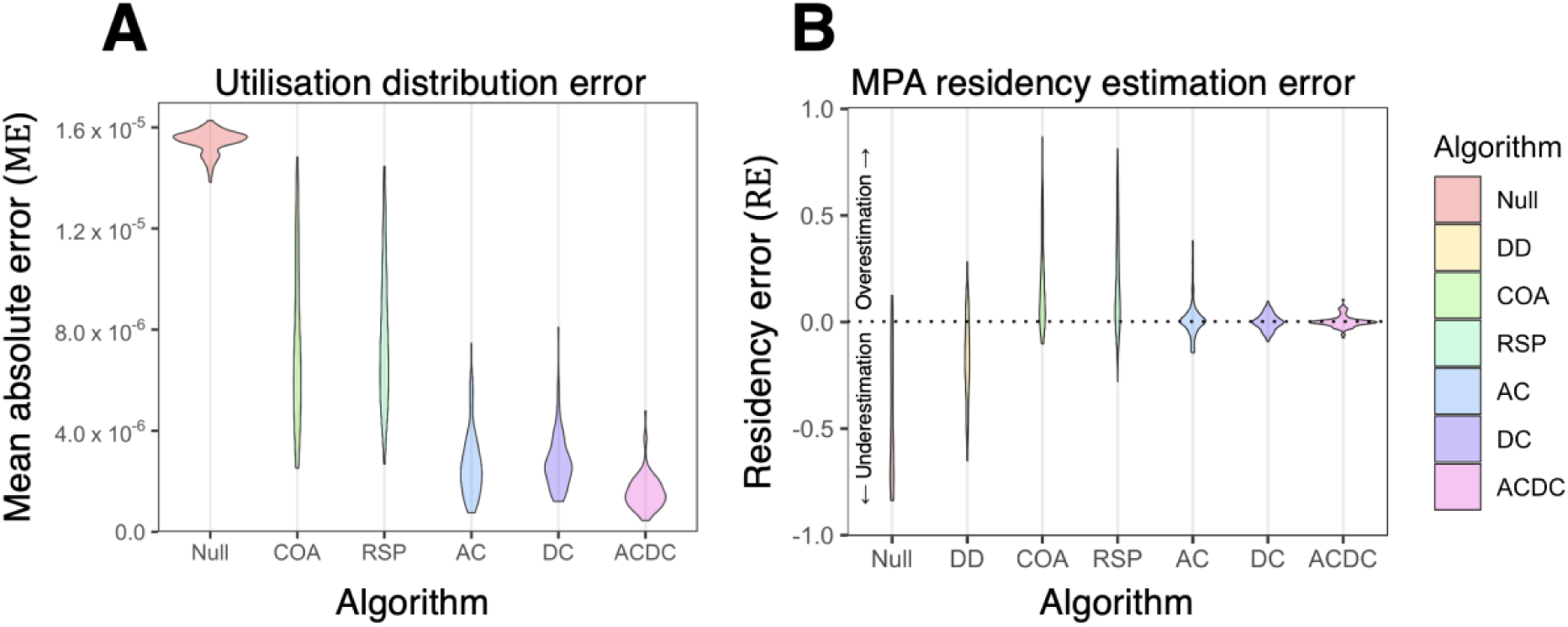
Algorithm performance for estimating (A) utilisation distributions and (B) residency. Violins show the distribution of error across 100 simulated paths. See text for abbreviations. For the full figure, see Figure S8.

Algorithm ranking for residency estimation (A1.3.3) was similar (Figures 2B and S8). The null model consistently underestimated residency. The proportion of days with detections (DD) benchmark also underestimated residency, while heuristic algorithms overestimated residency, except in open areas where receivers were absent. For heuristic algorithms, median residency error for the MPA was 12 % (standard error ≈ 22 %). Particle algorithms performed better, estimating residency in the MPA with a median error below 1 % and a precision (standard error) of 7.6 (AC), 3.7 (DC) and 2.8 (ACDC) %. These statistics represent >30-fold improvements in residency error and 2.8–8.3-fold improvements in precision compared to heuristic algorithms.

#### 2.2.2. Sensitivity analysis

The sensitivity analyses (A2) revealed algorithm sensitivity to mis-specification. We reimplemented algorithms with more restrictive and flexible parameterisations and examined variation in performance. Heuristic algorithms were successfully re-implemented for all simulated paths (A2.1). For particle algorithms, parameter mis-specification was associated with convergence failures (Figure S4). The restrictive depth observation model, which underestimated the amount of error in the depth observations, was particularly problematic. However, multiple independent runs of the particle filter facilitated convergence in some simulations.

Patterns of space use and residency estimates (A2.2) were relatively robust to algorithm parameterisation. The visual analysis (A2.2.1) revealed consistency in UDs between algorithm parameterisations (Figures S5–6 and S9–10). For heuristic algorithms, more flexible parameterisations produced more diffuse maps, but differences were small (Figures S5–6). For the particle algorithms that converged, maps were relatively robust to the degree of parameter mis-specification we explored (Figures S9–10). The main exception to this was mis-specification of the depth observation model (in the DC and ACDC algorithms), which affected the distribution of hotspots and patterns of space use (Figure S9). Particle-based maps were highly reproducible (Figure S10). Analyses of UD ME (A2.2.2) and residency (A2.2.3) produced similar results (Figures S11–12).

### 2.3. Real-world analysis

#### 2.3.1. Observations

Real tracking data from modelled skate exhibited diverse patterns (Figure S2). The number of detections per individual/month varied from 43–8887 (median = 1152). Some individuals were detected regularly while others went undetected for longer periods (up to 25 days). Most individuals were only detected around southerly receivers. The two exceptions were individual 25 (a mature female), which moved between southern and northern receivers over a 14-month period, and individual 28 (an immature female), which was primarily detected at southerly receivers but spent some time around northerly receivers in early 2017. Depth time series were variable and included extended periods (>1 week) with limited change alongside movements from a maximum depth (≈150 m) into shallower water and extensive transitions between shallow (<50 m) and deep (>200 m) areas (Figure S2).

#### 2.3.2. Main analyses

In our main analyses (A3), algorithms were successfully implemented (A3.1) for most individuals/months (Figure S13). The success rate was high for the heuristic algorithms (98– 100 %). In the particle algorithms, the convergence rate was high for the AC algorithm (100 %), but lower in the DC (79 %) and ACDC (85 %) algorithms.

Estimated UDs (A3.2.1) largely concentrated within the MPA but differed among algorithms (Figures 3–4 and S14–18). Of the two heuristic algorithms, the COA algorithm generally produced highly restricted UDs that concentrated around southerly receivers, although maps for two individuals (25 and 28) spanned a larger area (Figure S14). UDs from the RSP algorithm similarly centred in this region, but were more spread out (Figure S15). As for the COA algorithm, RSP UDs were driven by receiver locations. Maps for individuals detected around southerly receivers (e.g., 35) were unaffected by the temporal pattern of detections (Figures S2 and S15).

**Figure 3.**
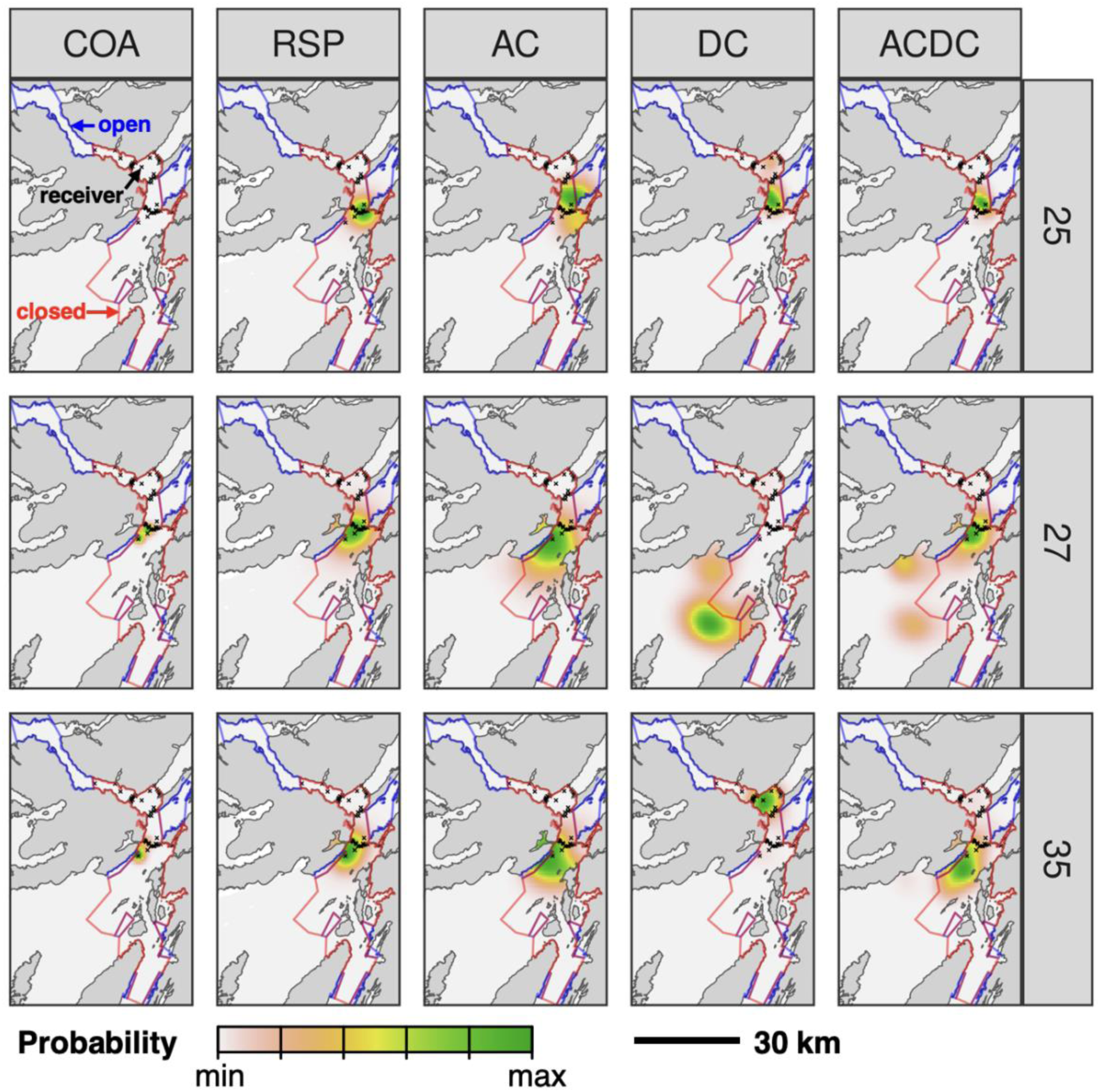
Example utilisation distributions (UDs) from tagged skate. Each panel shows a UD for April 2016 for a given individual (row) and algorithm (column). Receivers (**x**) and zones open/closed to fisheries are marked. In the top-left panel, the UD is so concentrated that it is hidden by receivers.

**Figure 4.**
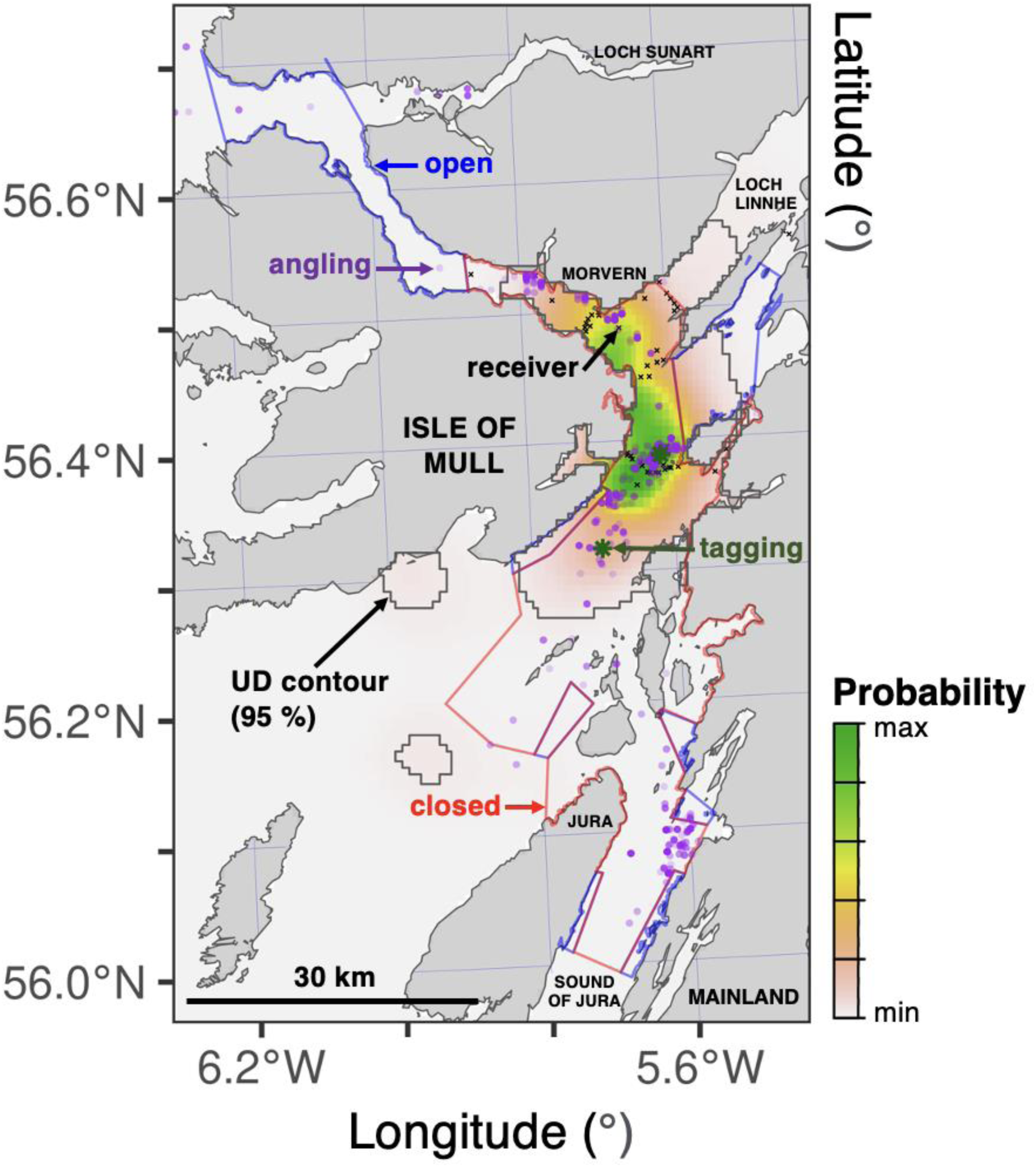
An overall utilisation distribution for modelled skate, reconstructed by the ACDC algorithm. Colours represent the probability that a modelled individual was located in a given cell at a randomly chosen time (within the analysed time series). The region containing 95 % of the probability mass is delineated by the black contour. Zones open/closed to fisheries, tagging locations (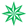), receivers (**x**) and locations where skate have been captured by anglers (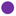) are marked.

Particle algorithms indicated more nuanced movement patterns (Figures 3 and S16–18). In the AC algorithm, most UDs spanned the southern receivers (where detections occurred) but were more diffuse and not exclusively centred on receivers (Figure S16). These patterns reflect array design, detection gaps and individuals’ capacity to move away from receivers. Some UDs suggest movement into receiver gaps in the array’s centre, fished zones and/or further south (beyond MPA boundaries). UDs from the DC algorithm exhibited both similarities and distinct features in the absence of acoustic constraints, the latter including hotspots in shallow to deep water south of the MPA (Figure S17). ACDC-derived UDs integrate features of AC and DC maps. Compared to the AC algorithm, maps were more concentrated within the MPA but also exhibited a redistribution of probability mass beyond MPA boundaries associated with the depth information (Figure S18). For both males and females detected around southerly receivers, the maps suggest notable movements to habitat patches beyond the MPA (especially off the Isle of Mull and north-western Jura). There are also indications of movements further north (including in the deep water off Morvern and beyond the MPA up Loch Linnhe). (For place names, see Figures 1/4.)

Within the MPA, our maps broadly align with skate presence records from angling (A3.2.2), which concentrates in this region (Figure 4). Potential movements beyond the MPA are poorly represented in angling records.

In line with UDs, residency estimates (A3.2.3) were high in the MPA but differed among algorithms (Figure 5). Detection day (DD) proportions varied from 0.07–0.94 (median = 0.48). As in simulations (Figures 2 and S8), heuristic residency estimates were consistently low in open zones (median = 0.05) and high in closed areas (median = 0.88) and the MPA as a whole (median = 0.93). These estimates were relatively insensitive to differences in detection patterns among individuals (Figure S2). Particle algorithm estimates were somewhat higher in open zones and lower in closed zones and the entire MPA. In the AC algorithm, residency ranged between 0.01–0.45 (median = 0.10) in open zones, 0.32–0.95 (median = 0.78) in closed zones and 0.68–0.97 (median = 0.90) in the entire MPA. In the absence of acoustic constraints, DC algorithm estimates were more variable, though median residency estimates were similar (0.04, 0.80 and 0.88). Median estimates from the ACDC algorithm were also broadly similar, ranging from 0.05 in open zones to 0.88 in closed zones and 0.92 in the entire MPA. Given limited data, seasonal trends in residency are unclear, although it is notable that ACDC residency estimates are broadly elevated in open areas over winter (when commercial fishing is permitted).

**Figure 5.**
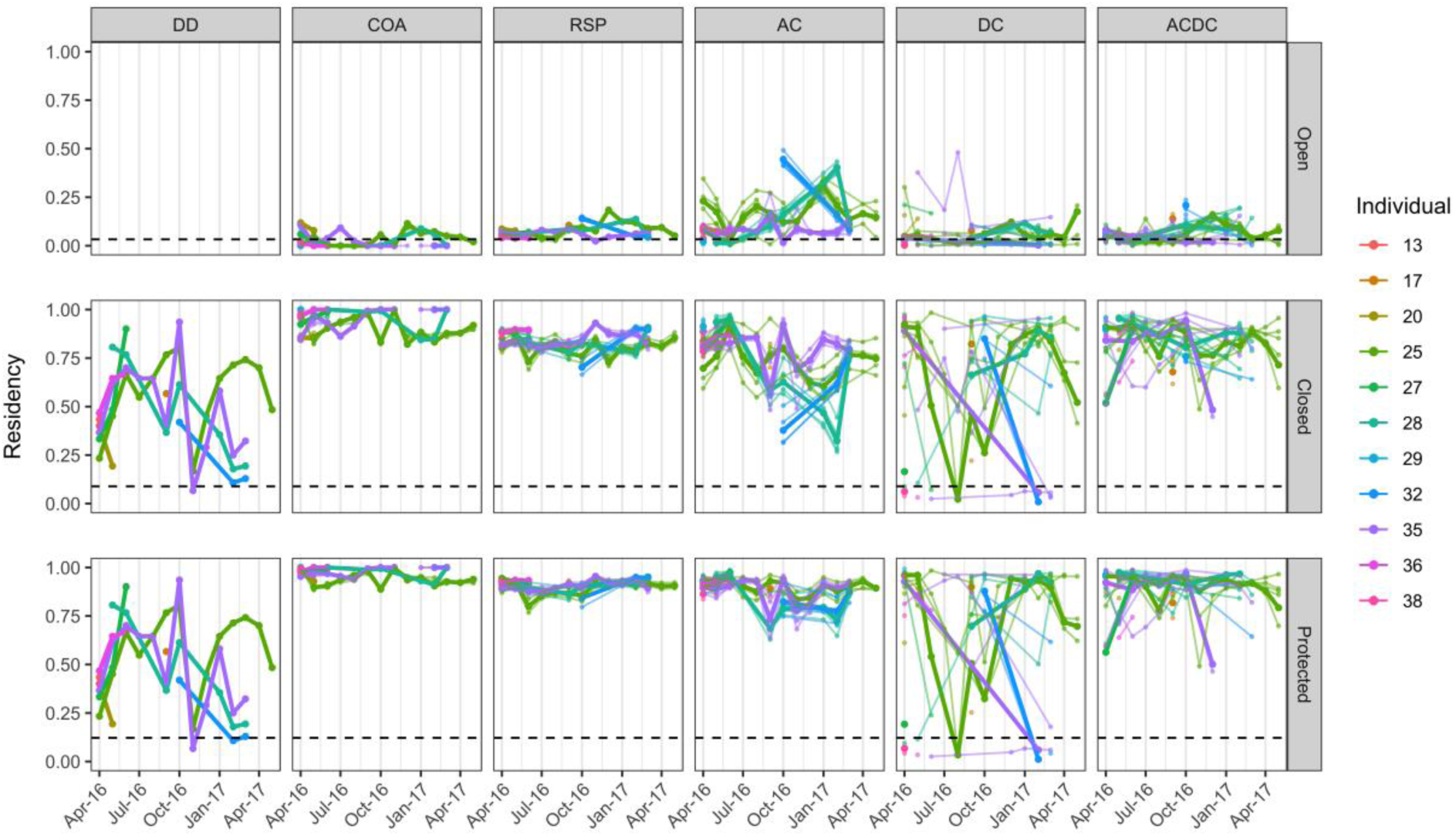
Residency of tagged skate in the MPA. Panels show individual residency time series for a given management area (row) and algorithm (column). The terms ‘Open’, ‘Closed’ and ‘Protected’ refer to zones within the MPA that are open or closed to fisheries and the entire MPA. Residency is the expected proportion of time spent in each area (in a given month). Thick points/lines are best-guess residency estimates; thinner points/lines are estimates from other algorithm parameterisations (sensitivity analysis). Estimates are only shown for months with sufficient data (hence gaps in some time series). Within panels, the feature of interest is the overall spread of estimates (rather than individual trajectories). Horizontal lines are null model estimates.

#### 2.3.3. Sensitivity analyses

In real-world sensitivity analyses (A4), particle algorithm implementation success (A4.1) was affected by algorithm parameterisation (Figure S13). Convergence rates were highest in AC algorithm implementations (98–100 %) and lowest in DC algorithm implementations (67–83 %). Lower convergence was associated with both restricted and flexible parameterisations.

In the UD analysis (A4.2.1), maps produced by different model parameterisations were broadly similar but differed in detail (Figure S19). As in simulations (Figures S3 and 5–6), heuristic algorithm UDs were largely unaffected by tuning parameters (Figure S19). In the AC algorithm, restrictive parameterisations generally concentrated maps and flexible parameterisations spread them out (Figure S19). DC algorithm sensitivity was more complex and included contractions, expansions and shifts in the distribution of hotspots. In the ACDC algorithm, restrictive and flexible parameterisations generally concentrated/expanded patterns of space use, weakening/strengthening the relative importance of habitat patches beyond the MPA, respectively (Figure S19). Where convergence was achieved, the restrictive depth observation model parameterisation in particular strengthened patterns of space use within the MPA and dampened the apparent importance of habitats further south.

Residency estimates (A4.2.2) showed a similar degree of sensitivity to algorithm parameterisation (Figure 5). Restrictive parameterisations were generally associated with elevated residency estimates while flexible parameterisations were associated with reduced estimates, especially in the DC algorithm. However, median residency estimates remained remarkably stable. In the ACDC algorithm, median estimates, accounting for all algorithm parameterisations, were 0.05 (in open zones), 0.87 (in closed zones) and 0.93 (in the entire MPA).

## 3. Discussion

This study sets a new state-of-the-art in animal tracking for conservation with underwater biotelemetry, building on decades of valuable work in this field. We reveal particle algorithms as a powerful methodology that can strengthen analyses of space use and residency, while addressing the limitations of long-established heuristic methods. Our analyses show that particle algorithms achieve substantial improvements in accuracy and precision, while heuristic algorithms can be difficult to tune, insensitive to the temporal pattern of detections and exhibit marked variation in performance. For the Critically Endangered flapper skate, particle algorithms suggested concentrated patterns of space use within an MPA, including time spent beyond receivers in zones that are open to fishing, as well as movements beyond MPA boundaries—significantly extending work on this and related species, with important implications for species conservation^32,38,39^. While heuristic methods have been valuable components of the animal-tracking toolbox^17,21,22^, particle algorithms provide a probabilistically sound statistical methodology that is sufficiently fast, flexible and accessible to meet the challenges of many modern animal-tracking datasets. We encourage their application to analyses of species’ movements and MPA efficacy across aquatic ecosystems^9,40^.

We developed a simulation analysis to calibrate heuristic algorithms and evaluate algorithm performance and sensitivity. For heuristic algorithms, previous studies have typically relied on default values^22^ or subjective judgement^20^ to set tuning parameters, such that variation in algorithm performance and sensitivity is poorly documented^19^. In our study system, for the COA algorithm, we found that a setting of Δ*T* = 2 days was marginally better than other settings, while for the RSP algorithm there was no optimal choice for the distance-error (e>r>.ad>) parameter on average. This insensitivity appears reassuring, but it calls into question the relevance of tuning parameters and their ability to envelop the movement and detection processes that generate observations, upon which the utility of heuristic methods rests^17,19^.

Our simulations revealed a clear ranking of algorithms for estimating patterns of space use and residency. Performance of the two heuristic algorithms was similar but highly variable: simulated patterns were captured well in some instances and poorly in others. In residency analyses, the widely used ‘detection days’ metric underestimated residency in the MPA, in line with the limited movement capacity of simulated individuals (which could linger within the MPA but beyond receivers). Meanwhile, heuristic algorithms (which restrict positional estimates within receiver arrays, even if individuals move further afield) overestimated residency by a median 12 % (standard error ≈ 22 %). These results are specific to our study system but pertinent for many real-world arrays, where behaviour of prevailing analytical methods remains understudied^17,19^.

Compared to heuristic methods, particle algorithms produced more accurate insights into space use and residency. There were clear benefits of data integration, with depth observations refining movements in the AC algorithm and acoustic observations helping to localise movements to relevant regions in the DC algorithm (which otherwise sometimes misplaced individual hotspots). Overall, residency was accurately estimated within a standard error of 7.6 (AC), 3.7 (DC) and 2.8 (ACDC) %. This quantification of ‘array precision’ should become an effective means to evaluate and improve monitoring programmes in future (Lavender et al., in prep). In this study, simulations suggest that data integration enabled up to three-fold improvements in estimates of space use and residency (lower mean error, higher precision). We anticipate considerable opportunities to extend this kind of analysis in other systems to evaluate (a) existing arrays, (b) alternative array designs and (c) the number of receivers required to estimate metrics of interest with a specified degree of accuracy (Lavender et al., in prep).

The above simulation results were broadly robust to algorithm parameterisation. Heuristic algorithms were highly insensitive to algorithm parameterisation, but even particle algorithms were generally robust to mis-specification of individual parameters (such as the maximum movement speed), given the presence of other constraints (such as detection range)^19^. The DC algorithm showed the greatest sensitivity, as expected given the absence of acoustic constraints and the heterogenous distribution of depth habitats in our study system. In other study systems, we recommend that studies leveraging multiple datasets conduct similar analyses to examine algorithm sensitivity^41^.

Our real-world analyses of flapper skate showcase significant enhancements in the use of acoustic telemetry for mobile species conservation. Within the MPA we study, we found that heuristic methods produced limited maps of space use that were driven by receiver positions (and insensitive to the temporal pattern of detections), while particle analyses indicated specific habitat patches in-between receivers that appear to be important, filling gaps in angling records and detection analyses^32^. The drivers of movement within these regions are unclear, but depth, temperature, light, sediment and/or prey preferences may play a role, as shown for other elasmobranchs^42^. A wide range of studies have linked detection patterns to habitat selection^43,44^ and this is an important next step in our research. By probabilistically representing individual movements, particle algorithms have clear potential to strengthen habitat-selection analyses and support the identification of sites, within and beyond MPAs, that may benefit from conservation measures^45,46^.

Within the MPA, our algorithms also suggested movements beyond receivers into fished zones. Given the sensitivity of elasmobranchs to trawling^5,29^, this result is potentially concerning and suggests further analyses of skate–fisheries interactions in our study area are warranted. Further afield, we recognise that many MPAs contain fished zones where receiver deployments are limited by fishing activity^7^. The capacity to investigate movements into these zones is an important development for quantifying the trade-offs between species conservation and extractive activities. This should inform marine spatial planning in many regions^47^.

By integrating tagging and tracking datasets with prior knowledge of species movement, we were also able to estimate individual residency at spatial scales relevant to management, notwithstanding limited receiver coverage. For analysed skate, our algorithms indicate a remarkably high degree of residency in the MPA. Earlier analysis of residency indices indicated that skate can spend weeks at a time around receivers^32^ but robust estimates of residency for flapper skate (and other mobile species) in regions with incomplete receiver coverage have remained elusive. In this study, we found the commonly used ‘detection days’ metric^17^ variable and poorly indicative of residency. Meanwhile, heuristic algorithms indicated consistently high MPA-scale residency across individuals, irrespective of detection gaps. Properly accounting for movements during detection gaps is a significant step forward in studies of the efficacy of MPAs for mobile species. For flapper skate, our best (ACDC) median residency estimate is 92 %, suggesting that the MPA is sufficiently large to protect analysed skate for much of the year. This result is robust to algorithm parameterisation. This work strengthens the evidence that MPAs can be effective conservation measures for mobile species and supports the use of permanent, rather than seasonal, fisheries closures for flapper skate and other species that display similar movement patterns^9,38,39^. However, fisheries closures should be accompanied by wider fisheries management measures to protect individuals that exhibit weaker site affinity^31^.

We also documented evidence for movements into specific habitat patches beyond MPA boundaries that may warrant protection. For flapper skate, these include an area of moderately deep (>100 m) water off Jura and a shallow (50 m) sandy–muddy habitat off the Isle of Mull that is bounded by a deeper (100 m) trench (punctuated by subsurface rocks). It is tantalising to notice characteristics of an egg nursery in this location^37^. Currently, the only known flapper skate nursery is in the Red Rocks and Longay MPA (further north). While sample size is limited, we found no strong evidence for movements towards Red Rocks, so the search is on to determine where flapper skate using the MPA lay eggs^37^. The identification of essential fish habitats is crucial for mobile species conservation and a key objective of many telemetry studies, including our own work^40^. The unique capacity of our algorithms to facilitate this research, via the identification of specific habitats beyond receivers that are used by tagged animals, is a useful step forward. This should support the establishment of connected MPA networks for mobile species^40^.

Improving the movement and observation models we use to model underwater biotelemetry data is an important task for future research. Our skate movement model would be informed by additional (e.g., accelerometer) data on individual behaviour and swimming speeds. Multi-sensor tags hold considerable promise for this in future and should be leveraged by other studies^11^. Our acoustic observation model would also be improved with additional (sentinel) tag deployments and expanded detection range testing. Such data are widely collected and should be incorporated in analyses^48^. In our study system, improvements to the depth observation model are a priority: in sensitivity analyses, different model parameterisations affected the localisation of hotspots within the MPA and the relative importance of habitat patches further afield. The key knowledge gap is the extent to which flapper skate exhibit benthic versus pelagic behaviour. In rugged bathymetric landscapes, fish behaviour affects the locational information provided by depth observations and the ease with which particle filters converge. In this study, we attribute convergence failures to potential inadequacies in the depth observation model and the difficulty of finding valid routes through a bathymetric maze. In complex environments, particle algorithms may struggle and discretisation of the state-space for filtering^49^ or tempered Hamiltonian Monte Carlo^50^ may be required. For flapper skate and other aquatic species, continued tagging and monitoring, coupled with further analytical development in these directions, will support improved analyses in years to come.

This work has significant implications for the spatial management of mobile species. By integrating animal movement modelling and electronic tagging and tracking data, we reveal how particle algorithms can represent movements beyond receivers, refine maps of space use and improve residency estimates. These developments enhance the value of biotelemetry data for mobile species conservation, informing MPA placement, size and management^9^. Our case-study analyses of flapper skate provide a concrete example: confirming the value of fisheries restrictions in an existing MPA; indicating patterns of habitat use; quantifying movements into fished zones; and highlighting specific habitats beyond the MPA that may warrant protection. These insights into space use, residency in regions of interest and unprotected habitats beyond receivers represent great steps forwards for aquatic conservation^44^. That being said, model precision remains dependent on data quality and algorithm-informed study design may be beneficial moving forward. This work should support the conservation of flapper skate and other threatened species across the globe^10,40^.

Particle algorithms have potential applications beyond spatial management. Acoustic telemetry and biologging are expanding rapidly^15,16^. With the growth of international telemetry networks, the spatial scale over which animals can be tracked is also widening. Analyses of telemetry data require statistical models that accurately reconstruct movements and represent uncertainty. Our integrative modelling framework is thus well-placed to strengthen research across the biotelemetry sphere, from ecological analyses of individual movements^44^ through to the conservation objectives addressed in this paper^40^. We foresee opportunities to refine habitat selection analyses^46^, investigate co-occurrence patterns^51^ and support animal-borne oceanography^52^, fisheries management^53^, climate change mitigation^54^, habitat restoration^55^ and impact assessments^56^. The methods are relevant for research projects across disparate systems^15,16^ and have the potential to support progress towards strategic objectives, such as Sustainable Development Goal 14^40^. We hope that this study encourages advances towards this goal in the second half of the United Nations Decade of Ocean Science for Sustainable Development.

## 4. Methods

### 4.1. Study system

We selected a 14,000 km^2^ case-study system in Scotland (Figure 1). The bathymetry encompasses shallow-water platforms, deep basins and channels up to 350 m in depth. Within the site, the Loch Sunart to the Sound of Jura MPA spans 741 km^2^ and a depth range of 0–290 m. Current management prohibits commercial fishing except in eight, seasonally (October– March) fished zones. For further description, including bathymetric data sources, see Supporting Information §1 and Table S1.

From 2016–17, a passive acoustic telemetry system comprising 58 Vemco receivers was established in part of the MPA^32^. Skate were captured in the same area and tagged with acoustic transmitters and archival (depth) tags^35^. For full details, including array and tag properties, see Supporting Information §2.

### 4.2. Workflow

We conducted simulation-based and real-world analyses of animal movements (see Table S2 for an overview and §4.3–4 for details). In each analysis, we analysed movements using two heuristic algorithms (COAs and RSPs) and our particle algorithms. The workflow comprised three stages: (a) coordinate estimation, (b) mapping and (c) residency estimation, which are explained in general terms in §4.2.1–3 before the details of our analyses. Algorithms were implemented in R>, v.4.3.1^57^, using the p>a>t>t>e>r>^28^ and R>S>P>^22^ packages. All code is available online^58^.

#### 4.2.1. Coordinate estimation

##### COAs

The first algorithm we used to estimate coordinates for mapping was the COA algorithm. This estimates coordinates as weighted averages of receiver locations where detections were recorded over sequential time intervals (duration: Δ*T*). We selected three Δ*T* values (an ‘optimal’ value, a restrictive value and a more flexible value) to examine algorithm performance and sensitivity (see §4.3).

##### RSPs

The RSP algorithm interpolates coordinates along the shortest paths between the receivers. We used the default settings for tuning parameters except e>r>.ad>, which tunes coordinate weights for mapping. As above, we selected three values by simulation (see §4.3). For implementation details, see Supporting Information §3.

##### Particle algorithms

We also implemented the acoustic-container (AC), depth-contour (DC) and acoustic-container depth-contour (ACDC) particle filtering–smoothing algorithms^19^. To implement these algorithms, we formulated a Bayesian state-space model (SSM) for the location (strictly ‘state’) ***s*** an individual through time (*t* ∈ 1, 2, …, *T*}), given the (acoustic and archival) observations (***y***_1:*T*_), that is, *f*(*s*_1:*T*_| ***y***_1:*T*_). Our SSM represents an underlying behavioural switching movement process that includes a ‘low activity’ state (encompassing resting) and a more active state. Movements are connected to acoustic observations (detections, non-detections) by a Bernoulli observation model in which detection probability declines logistically with distance from receivers. The depth observation process is described by a truncated Gaussian distribution centred on the seabed (capturing skate’s benthic lifestyle) with a variance accounting for observational and bathymetric uncertainty. For the model formulation, see Supporting Information §4.1 (mathematics), Table S3 (notation summary) and Figure S1 (visualisation). We considered a ‘best-guess’ parameterisation as well as a restrictive and flexible parameterisation for the movement and observation sub-models. SSM parameterisations were based on prior research, domain knowledge and a literature review. See Supporting Information §4.2–3 (for details) and Table S4 (for a summary of parameter values).

Particle filtering and smoothing algorithms perform model inference for the SSM, targeting the marginal distribution *f*(*s*_*t*_ | ***y***_1:*T*_). The distribution of individual locations is approximated by a Monte Carlo simulation of *N* weighted particles, which represent candidate positions of the individual. In the particle filter, a movement model simulates particle movement from one time step to the next; observation models weight particles in line with their compatibility with the data; and a resampling step duplicates or eliminates particles accordingly. The AC, DC and ACDC algorithms differ only in the data incorporated during this process (acoustic, depth or both datasets). The number of particles is a trade-off between speed and convergence: sufficient particles are required to ensure that at least some particles are compatible with the data at every time step. After subsequent particle smoothing, the result is a set of particles that approximate *f*(*s*_*t*_ | ***y***_1:*T*_). For implementation details, see Supporting Information §4.4.

#### 4.2.2. Mapping

For COAs and particle algorithms, we generated maps of space use, i.e., utilisation distributions (UDs), by kernel smoothing estimated coordinates^28^. For the RSP algorithm, a dynamic Brownian-bridge movement model is used to smooth coordinates^22^. For implementation details, see Supporting Information §5.

#### 4.2.3. Residency

Alongside UDs, we estimated the proportion of time spent in selected areas (i.e., residency). We considered residency in our study system in three areas: zones (a) open and (b) closed to fisheries and (c) the entire MPA. Residency was estimated as the proportion of (i) UD volume (for heuristic algorithms) or (ii) resampled particles (for particle algorithms) in each area. We also considered (iii) the proportion of days with detections in each area (a standard metric) for comparison.

### 4.3. Simulation-based analysis

Our first analysis was a simulation-based analysis (see Table S2 for the overview). In this analysis, we simulated individual movements and corresponding observations in our study system and reconstructed patterns of space use using COAs, RSPs and particle algorithms. We used simulation results to calibrate heuristic algorithms for real-world analyses, evaluate algorithm performance and sensitivity, and validate the use of particle algorithms for later analyses (see §4.4). Full details are available in Supporting Information §6.

In outline, we simulated movements from ‘tagging locations’ within the study area for 100 hypothetical flapper skate over a one-month period, according to our best-guess skate movement model (see Supporting Information §6.1). For each movement path, we simulated acoustic and depth observations, following best-guess models for the aforementioned acoustic and depth observation processes. For each path, we generated a ‘true’ UD via kernel smoothing. Residency was calculated as the proportion of path steps in each area. Using the simulated datasets, we assessed the performance and sensitivity of algorithms applied to simulated observations in terms of how well they recovered path UDs and residency (see Supporting Information §6.2–3). In the following description, we label performance and sensitivity analyses as ‘A1’ and ‘A2’, respectively. Analytical steps are labelled in the same way (as A1.1, A1.2, etc.), following Table S2.

In performance analyses, we evaluated the skill with which algorithms reconstructed simulated patterns of space use and residency (A1). This analysis included a null model (uniform UD, excluding land) alongside heuristic and particle algorithms parameterised with optimal or best-guess parameters (A1.1). For heuristic algorithms, a simulation approach was used to select optimal parameter values: for each path, we computed the mean absolute error (ME) between the UD for each simulated path and reconstructed UDs for a range of candidate parameter values and identified the parameter value with the lowest ME on average (A1.1.1). Particle algorithms were parameterised according to the data-generating processes (A1.1.2). Heuristic algorithms were implemented for each acoustic time series and particle algorithms (AC, DC and ACDC) were implemented for each (a) acoustic, (b) depth and (c) combined dataset, respectively (A1.2). We then evaluated algorithm performance (A1.3) by visually comparing simulated and reconstructed UDs for three selected paths (A1.3.1), the distribution of ME between all simulated and reconstructed UDs (A1.3.2) and the distribution of residency error (estimated residency minus true residency) between simulated and reconstructed patterns, by algorithm (A1.3.3). Residency error was quantified for the null model and the detection days metric (as benchmarks), as well as the heuristic and particle algorithms.

In sensitivity analyses, we used a subset of paths (1–3) to explore algorithm sensitivity (A2). In this analysis, we re-implemented the algorithms with restrictive and flexible parameterisations (A2.1). For the particle algorithms (which are stochastic), we ran each algorithm implementation three times to examine reproducibility. We then analysed algorithm sensitivity and reproducibility (A2.2) by visualising patterns of space use (A2.2.1), ME (A2.2.3) and residency error (A2.2.3) for each algorithm parameterisation/implementation.

For our study area, the simulation analyses (a) confirmed that particle algorithms outperform heuristics, (b) revealed algorithm sensitivity and (c) demonstrated that stochastic particle runs produce consistent results. By visually comparing maps between AC, DC and ACDC algorithms, we also gauged the relative importance of acoustic and/or depth datasets in refining maps of space use.

### 4.4. Real-world analyses

In real-world analyses (A3–4), we analysed movement patterns of tagged skate (Figure S2, Table S5). Data were sourced from a study of 42 individuals tagged with acoustic and archival tags (33 of which were detected)^32^. We selected 13 individuals (30 %) with sufficient data for analysis. (For data processing, see Supporting Information §7.1.) For each individual, we analysed movements over each month using (a) the acoustic data, (b) the depth data and (c) the combined data. We analysed 48 individual/month time series in total (1–14 months per individual).

Following the simulation workflow (§4.3), for each individual, for each month, we analysed data using heuristic algorithms (acoustic datasets only) and the particle algorithms (acoustic, depth and combined datasets). In our main analyses (A3), each algorithm was implemented once using optimal (heuristic) or best-guess (particle algorithm) parameters (A3.1). We estimated UDs for each dataset (A3.2.1). We assessed similarities and differences among individuals and algorithms visually, given limited data. UDs were aggregated (A3.2.2) to map the overall pattern of space use and assess overlap with long-term (1975–2024) skate presence angling records^59^. Residency was estimated as in simulations (A3.2.3). In sensitivity analyses (A4), we re-implemented algorithms with restricted and flexible parameterisations (A4.1) and analysed changes in UDs (A4.2.1) and residency (A4.2.2). For the overview, see Table S2. For full details, see Supporting Information §7.2.

## Data availability statement

Data are available from NatureScot and Marine Scotland Science. Code is archived on Zenodo at https://doi.org/10.5281/zenodo.1480516158.

## Author contributions

This study builds on the Movement Ecology of Flapper Skate (MEFS) project, established by JT and developed through EL’s PhD Studentship at the University of St Andrews (2018–22), supervised by PJW, MJ and JI plus JT and SS (Principal Investigators) and supported by SB, JD, GC and DA. EL conceived and designed the present study during a postdoctoral fellowship at Eawag (2023–24) with inputs from all authors, especially AS, CA and HM (Principal Investigator). Tracking data were made available through the MEFS project (via JT). DA provided expertise and WeStCOMS model outputs for the study area that informed the analysis. JD provided Skatespotter data. EL analysed the data and led the writing of the manuscript. All authors shared expertise, provided inputs and approved publication.

### Acknowledgments

We thank Ronnie Campbell, Roger Eaton, Francis Neat and the Flapper Skate Working Group for supporting the MEFS project and our research. Thanks to Jelger Elings and Luis Habersetzer for additional support.

