## Supporting Information for "Animal tracking with particle algorithms for conservation"

Supporting information, figures and tables

#### Supporting materials

**Contents** **Pages**


| | |
|---|---|
| **Supporting information** | **1–29** |
| **1. Study site** | **1–3** |
| 1.1. Domain | 1 |
| 1.2. Bathymetry | 1–3 |
| 1.2.1. Data sources | 1 |
| 1.2.2. Dataset assembly | 2 |
| 1.2.3. Dataset uncertainty | 2–3 |
| 1.3. Flow regime | 3 |
| **2. Passive acoustic telemetry** | **4–5** |
| 2.1. Array design | 4 |
| 2.2. Acoustic and archival tags | 4 |
| 2.3. Detection probability | 4–5 |
| **3. Heuristic algorithms** | **5–6** |
| **4. Particle algorithms** | **6–19** |
| 4.1. State-space model | 6–10 |
| 4.1.1. Posterior | 6–7 |
| 4.1.2. Prior | 7–9 |
| 4.1.3. Likelihood | 9–10 |
| 4.2. Model parameterisation | 10–17 |
| 4.2.1. Movement model | 10–14 |
| 4.2.1.1. Literature review | 10–12 |
| 4.2.1.2. Parameter settings | 12–14 |
| 4.2.2. Acoustic observation model | 14–16 |
| 4.2.3. Depth observation model | 16–17 |
| 4.3. Model uncertainty | 17–18 |

**Contents** **Pages**

4.4. Model inference 18–19

**5. Mapping** **19–20**

5.1. Kernel smoothing 19–20

5.2. Brownian bridges **20**

**6. Simulation analyses** **20–25**

6.1. Data simulation 20–22

6.2. Performance analyses (A1) 22–24

6.3. Sensitivity analyses (A2) 24–25

**7. Real-world analyses** **26–29**

7.1. Datasets 26

7.2. Analyses (A3–4) 26–29

7.2.1. Particle algorithm initialisation 27–28

7.2.2. Particle algorithm implementation 28–29

**Contents** **Pages**

**Supporting figures** **30–53**

| | | |
|---|---|---|
| Figure S1. | A state-space model for tracking flapper skate. | 30–31 |
| Figure S2. | Observed time series. | 32–33 |
| Figure S3. | Simulation analysis: heuristic algorithm performance profiles for (A) COAs and (B) RSPs. | 34 |
| Figure S4. | Simulation analysis: particle algorithm convergence properties for (A) performance and (B) sensitivity analyses. | 35 |
| Figure S5. | Simulation analysis: COA algorithm performance and sensitivity. | 36 |
| Figure S6. | Simulation analysis: RSP algorithm performance and sensitivity, following Figure S5. | 37 |
| Figure S7. | Simulation analysis: particle algorithm performance, following Figure S5. | 38 |
| Figure S8. | Simulation analysis: model performance for estimating residency. | 39 |
| Figure S9. | Simulation analysis: particle algorithm sensitivity. | 40–41 |
| Figure S10. | Simulation analysis: particle algorithm repeatability, following Figure S9. | 42–43 |
| Figure S11. | Simulation analysis: algorithm sensitivity for mapping space use. | 44 |
| Figure S12. | Simulation analysis: algorithm sensitivity for estimating residency. | 45–46 |
| Figure S13. | Real-world analysis: particle algorithm convergence properties. | 47 |
| Figure S14. | Real-world analysis: COA algorithm maps from the main analysis. | 48 |
| Figure S15. | Real-world analysis: RSP algorithm maps from the main analysis, following Figure S14. | 49 |
| Figure S16. | Real-world analysis: AC algorithm maps from the main analysis, following Figure S14. | 50 |
| Figure S17. | Real-world analysis: DC algorithm maps from the main analysis, following Figure S14. | 51 |
| Figure S18. | Real-world analysis: ACDC algorithm maps from the main analysis, following Figure S14. | 52 |
| Figure S19. | Real-world analysis: sensitivity in maps of space use. | 53 |

**Contents** **Pages**

**Supporting tables** 54–62

Table S1. Bathymetric data sources. 54–55

Table S2. Summary of the workflow of simulation and real-world analyses. 56–57

Table S3. Summary of notation. 58–59

Table S4. Parameter values for algorithm implementations. 60–61

Table S5. Summary of modelled flapper skate. 62

**References** **63–66**

Supporting information, figures and tables

**Abbreviations**

| | |
|---|---|
| AC | : Acoustic-container |
| ACDC | : Acoustic-container depth-contour |
| COA | : Centre of activity |
| DC | : Depth-contour |
| MPA | : Marine protected area |
| RSP | : Refined shortest path |
| UD | : Utilisation distribution |

### Supporting information

#### 1. Study site

##### 1.1. Domain

Our study system forms a 14,000 km$^2$ area on the west coast of Scotland. The study system is centred in the Firth of Lorn and was chosen to envelope the Loch Sunart to the Sound of Jura Marine Protected Area (MPA) and surrounding regions, while remaining sufficiently small in size for analyses[1].

##### 1.2. Bathymetry

###### 1.2.1. Data sources

We required a bathymetric raster to reconstruct patterns of space use for tagged flapper skate within the study area[2]. A raster for the study system was built using data from three sources:

- **A Firth-of Lorn dataset** (5 x 5 m), compiled from an Order 1a hydrographic survey, obtained from Howe et al. (2014).
- **Supplementary datasets** for selected regions within the study area, obtained from Admiralty (https://seabed.admiralty.co.uk/).
- **A Scotland-wide dataset** (6 arc-second resolution), obtained from Digimap © Crown copyright and database rights [2019] Ordnance Survey (100025252).

For full details on bathymetric data sources, see Table S1. Coastline data (for UD estimation[3] and/or mapping) were obtained from Digimap (Westminster constituencies) and GADM (wider context) (Global Administrative Areas, 2022).

---

[1] In our analysis, the study area is represented as a (bathymetric) grid. Smaller grids help to manage computational demands, including memory requirements, computation time and disk space, for RSP estimation (see §3), initialisation of the particle algorithms (see §7.2) and UD estimation (see §5).

[2] We reconstructed patterns of space use using the COA, RSP and particle (AC, DC and ACDC) algorithms. In the RSP algorithm, the bathymetric raster is used to distinguish water/land for shortest-path generation (Niella et al., 2020). In the particle algorithms, the raster is required to define the region within which movements can occur and to incorporate depth observations (see §4).

[3] Coastline data were used in UD estimation to account for habitat edges (see §5).

##### 1.2.2. Dataset assembly

**Study site datasets.** We assembled two datasets for the study system: a high-resolution dataset (named 'bathy-high') and a lower resolution dataset (named 'bathy-low'). Datasets were assembled as follows:

- **Merger.** The Firth of Lorn and supplementary datasets were merged onto a 5 x 5 m grid (556,740,000 cells) with a Universal Transverse Mercator (Zone 29N) Coordinate Reference System.
- **Resampling.** All supplementary datasets were resampled for the merger using bilinear interpolation.
- **Smoothing.** Deep spikes in the merged dataset, which reflect errors in bathymetric soundings (and are particularly prevalent along coastlines), were processed. Spikes were flagged in cells where the bathymetric depth was more than 50 m deeper than the median depth of the surrounding eight cells and cut back to the median depth. Visual assessment of this heuristic suggested it was effective.
- **Aggregation.** For selected analyses, bathymetric data were aggregated onto a 500-by-500-pixel grid using the arithmetic mean[4] (see §5–7). All receivers were in valid locations (i.e., in water) on the aggregated bathymetric dataset (bathy-low).

**Scottish dataset.** We processed the Scotland-wide dataset for RSP analyses[5] (see §5).

- **Cropping.** Data were cropped to the area spanned by our 'bathy' grids, expanded by 200 km in the four cardinal directions.
- **Resampling.** Data were resampled to match the resolution of bathy-low.
- **Merger.** Within the domain of the bathy-low dataset, we replaced the values on the Scottish dataset with those from bathy-low to ensure consistency between analyses.

##### 1.2.3. Dataset uncertainty

[4] This corresponds to a grid resolution $\approx 200 \times 278$ m.
[5] The RSP algorithm is one of the algorithms we used to reconstruct patterns of space use (Niella et al., 2020). This algorithm uses a dynamic Brownian bridge movement model which required a larger grid for successful UD estimation.

We required estimates of bathymetry uncertainty to reconstruct patterns of space use for tagged skate with the particle algorithms[6] (see §4.2.3). Uncertainty in bathymetric datasets depends on survey accuracy and subsequent processing:

- **Order 1a survey.** The Howe et al. (2014) survey was conducted to an industry standard, 95 % confidence vertical-uncertainty limit of $\sqrt{(0.500 + (0.013\ b(\boldsymbol{s}))^2}$, where $b(\boldsymbol{s})$ is the bathymetric depth in the location $(s_x, s_y)$ defined by $\boldsymbol{s}$ (International Hydrographic Organization, 2020). This ranges from 0.71 m to 3.85 m at 0 m and 290 m (the maximum depth in the domain of the dataset), respectively.
- **Order 2 surveys.** The accuracy of the supplementary datasets is uncertain. However, for Order 2 surveys, the bathymetric uncertainty at 350 m depth may be up to ± 8.08 m (International Hydrographic Organization, 2020).

Based on these statistics, we considered a bathymetric uncertainty of ± 10 m for the bathy-high dataset, given the 95 % confidence threshold, the merger of multiple datasets and spike processing. For the bathy-low dataset, there is an additional aggregation uncertainty that we represented (see §4.2.3).

#### 1.3. Flow regime

Information on the flow regime was required to parameterise our particle algorithms (see §4.2.1):

- **Tidal range.** Within the study area, tidal range can be up to ± 3 m (based on tidal charts).
- **Storm surges.** Tidal ranges may be compounded by ± 2 m storm surges (based on personal experience).
- **Flow regime.** The flow region is complex, driven by the interaction between the tides and bathymetry. In the West Scotland Coastal Ocean Modelling System (Aleynik et al., 2016; Davidson et al., 2021), in our study system, average bottom currents are 0.09 $ms^{-1}$ but can exceed 4 $ms^{-1}$. In the Gulf of Corryvreckan tidal channel (which is 3.5 km in length), current speeds reaching 4.75 $ms^{-1}$ have been recorded (Armstrong et al., 2021).

[6] The DC and ACDC algorithms incorporate depth observations, by measuring the probability of each depth observation in places differing in bathymetric depth, which requires a measure of bathymetric uncertainty.

#### 2. Passive acoustic telemetry

##### 2.1. Array design

From 2016–17, a passive acoustic telemetry system was established in the study site to monitor skate (Lavender et al., 2021b). In total, 58 Vemco VR2, VR2W and VR2AR 69 kHz receivers were deployed (Figure 1). Array configuration was driven by various research projects and logistics. Array coverage fluctuated in line with changes in receiver number and placement, with aggregated 50 % detection-probability regions spanning approximately 4–14 $km^2$ (Lavender et al., 2021b).

##### 2.2. Acoustic and archival tags

Flapper skate were captured by angling in and around the array and given acoustic and/or archival tags.

**Acoustic transmitters.** The acoustic tags (Vemco V13s and Thelma Biotel MP-13s) were programmed with random transmissions every 30–90 s (over an 18-month battery life). Tag power output was 147 dB and 153 dB re 1 uPa at 1 m for the V13 and MP-13 tags, respectively (Lavender et al., 2021b).

**Archival tags.** The archival tags (Star Oddi Milli-TDs) were programmed to record depth (± 4.77 m) every two minutes, until their removal from recaptured skate (Lavender et al., 2021a).

##### 2.3. Detection probability

We required estimates on detection probability to parameterise our particle algorithms (see §4.2.2). Drift tests of our array are described by Klöcker (2019) and summarised by Lavender et al. (2021b). The protocol was as follows:

- **Set up.** Tests were conducted (with limited resources) for a subset of receivers on 19th–20th October 2016.

- **Transmitter deployment.** For each test, two acoustic transmitters were deployed on a line from a boat (approximately 5 m below the surface and 2 m above the seafloor respectively).
- **Drift.** With the boat's engines and depth sounders turned off, the boat was allowed to drift away from each receiver. Locations and transmissions were recorded by a Vemco VR100 Digital Tracking Receiver.
- **Modelling.** Detection-probability relationships were explored by modelling acoustic observations (detections, non-detections), inferred from receiver records versus known transmissions, as a function of the distance between receivers and transmitters (and other variables) (Klöcker, 2019).

Drift test analyses demonstrated that detection probability declines from 0.97 near to receivers to 0.5 by 400–450 m (Klöcker, 2019). The maximum observed detection range was 708 m. However, in the nearby Loch Etive study system, a detection range of 767 m for V13 tags was recently measured (James Thorburn, personal communication). Considering limited testing, the maximum detection range in our system likely exceeds these thresholds (see §4.2.2).

#### 3. Heuristic algorithms

We reconstructed patterns of space use in simulation (§6) and real-world (§7) analyses using two heuristic algorithms as well as our particle algorithms:

- **COA algorithm.** The COA algorithm was implemented with the `coa()` function in the `patter` package (Lavender, 2024; Lavender et al., 2024b).
- **RSP algorithm.** The RSP algorithm was implemented with the `runRSP()` function in the `RSP` package[7] (Niella et al., 2020).

The algorithms depend on the following parameters:

- **$\Delta T$.** The COA algorithm depends on the time interval over which detections are averaged ($\Delta T$). In our analyses, we used an 'optimal' value, as well as a more restrictive and flexible setting, selected by simulation[8] (see §6.2).

[7] The `runRSP()` routine interpolates 'relocations' along the shortest paths between receivers for UD estimation. Shortest paths are computed using a transition layer that defines horizontal, in-water distances between receivers, which we built from our aggregated bathymetry dataset (bathy-low).
[8] Multiple values were selected to examine algorithm sensitivity.

- `er.ad`. The RSP algorithm depends on multiple tuning parameters. We selected three values for the `er.ad` parameter, which tunes the rate at which uncertainty around interpolated 'relocations' grows with distance from a receiver (see §6.2). The detection range ($\gamma$) was held constant at 3000 m, which is our best-guess for the study system (see §4.2.2). Default settings were used for the other parameters.

Coordinates estimated by COA and RSP algorithms were translated into UDs (see §5 for generic details). In selected analyses, UDs were used to estimate residency from the proportion of their volume in particular areas[9]. The analyses in this paper are outlined in Table S2. Implementation details are provided in sections §6–7.

#### 4. Particle algorithms

##### 4.1. State-space model

Alongside heuristic algorithms, we reconstructed patterns of space use using particle algorithms (Lavender et al., 2024a). Unlike heuristic methods, particle algorithms are based on a formal statistical (state-space) model of individual movements for which model inference is performed. In this section, we formalise a discrete-time model of skate movements in our study system in general terms, following Lavender et al. (2024a). See Figure S1 for a visualisation and Table S3 for a summary of notation. In subsequent sections, we explain model parameterisation (§4.2–3) and model inference using particle algorithms (see §4.4).

###### 4.1.1. Posterior

The model represents the joint distribution $f(\boldsymbol{s}_{1:T} \mid \boldsymbol{y}_{1:T})$ of a skate's state ($\boldsymbol{s}$) given the data ($\boldsymbol{y}$) through time ($t \in \{1, 2, \ldots, T\}$). The individual's state is denoted $\boldsymbol{s}_t = (s_{x,t}, s_{y,t}, \phi_t, \text{behaviour}_t)$ and comprises its two-dimensional location $(s_x, s_y)$ on (or near) the seabed, its heading ($\phi_t$) and behaviour[10]. The joint distribution is proportional to a prior $f(\boldsymbol{s}_{1:T})$ (the movement process) and a likelihood $f(\boldsymbol{y}_{1:T} \mid \boldsymbol{s}_{1:T})$ (the observation process):

[9] Residency is defined as the proportion of time spent in a particular region.
[10] We consider two behavioural states: a 'low activity' state (including resting) and an 'active' state (see §4.1.2). Behavioural states were defined *a priori*: we examined individual depth (vertical activity) time series and distinguished 'low activity' and 'active' behavioural states by a vertical activity threshold of 0.25 m, following

$$f(\boldsymbol{s}_{1:T} \mid \boldsymbol{y}_{1:T}) \propto f(\boldsymbol{s}_{1:T})\, f(\boldsymbol{y}_{1:T} \mid \boldsymbol{s}_{1:T}). \qquad \text{eqn 1}$$

We consider three versions of this model, with the same prior, but different observation processes for (a) acoustic data, (b) archival data and (c) the combined dataset. The models (and corresponding inference algorithms) are labelled AC, DC and ACDC respectively (Lavender et al., 2024a).

###### 4.1.2. Prior

We represent $f(\boldsymbol{s}_{1:T})$ as a discrete-time Markovian process:

$$f(\boldsymbol{s}_{1:T}) = f(\boldsymbol{s}_{t=1}) \prod_{t=2}^{T} f(\boldsymbol{s}_t \mid \boldsymbol{s}_{t-1}). \qquad \text{eqn 2}$$

**Initialisation.** The term $f(\boldsymbol{s}_{t=1})$ is the probability density function of the individual's initial state[11]. We treat the initial location as known (see §6.2), or uniform within some subset of the study area (see §7.2.1) compatible with the initial observations ($A$) and assume a uniformly distributed initial heading, i.e.,

$$f(\boldsymbol{s}_{t=1}) = \text{Uniform}\big(s_x, s_y; A\big) \cdot \text{Uniform}(\phi; -\pi, \pi). \qquad \text{eqn 3}$$

**Correlated random walk.** The term $f(\boldsymbol{s}_t \mid \boldsymbol{s}_{t-1})$ is the probability density of moving from $\boldsymbol{s}_{t-1} \rightarrow \boldsymbol{s}_t$. We model movement as a two-dimensional, behavioural switching correlated random walk, based on step lengths and turning angles i.e.,

$$\boldsymbol{s}_t \mid \boldsymbol{s}_{t-1} = \big(s_{x,t-1} + d_t \cos(\phi_{t-1} + \Delta\phi_t),\ s_{y,t-1} + d_t \sin(\phi_{t-1} + \Delta\phi_t),\ \phi_{t-1} + \Delta\phi_t,\ \text{behaviour}_t\big), \qquad \text{eqn 4}$$

where $d$ is the step length, $\phi$ is the heading and $\Delta\phi_t$ is the change in heading (turning angle)[12].

The movement density is represented by $f(\boldsymbol{s}_t \mid \boldsymbol{s}_{t-1})$. The density of the current step length and heading ($f_{d,\phi}$) is given by the densities of the step length ($f_d$) and turning angle ($f_{\Delta\phi}$):

---

Lavender et al. (2022b) (see §6.1 and §7.2.2). We recognise that we ought to infer behaviour alongside the other state variables jointly in a Bayesian framework; however, this is numerically expensive. Our approach here might lead to a slight underestimation of uncertainty, but is cheaper.

[11] The state $\boldsymbol{s}_t = (s_{x,t},\ s_{y,t},\ \phi_t,\ \text{behaviour}_t)$ includes behaviour. However, as behaviour states were assigned *a priori,* we drop this dimension from eqn 3 for clarity.

[12] During model inference, this sampling procedure is implemented iteratively until a valid location (in water) is found (see eqn 8 and §4.4).

$$f_{d,\phi}(d_t, \phi_t \mid \phi_{t-1}) = f_d(d_t)\, f_{\Delta\phi}(\phi_{t-1} - \phi_t). \qquad \text{eqn 5}$$

The step length and turning angle define the change in location in a polar coordinate system, which is mapped to the change in $x$ and $y$ coordinates $(s_x, s_y)$ by:

$$\boldsymbol{g}\left((\Delta s_x, \Delta s_y)\right) = (d_t, \phi_t) = \left(\sqrt{{\Delta s_x}^2 + {\Delta s_y}^2}\,, \arctan(\Delta s_y \,/\, \Delta s_x)\right), \qquad \text{eqn 6}$$

where $\Delta s_x = s_{x,t-1} - s_{x,t}$, $\Delta s_y = s_{y,t-1} - s_{y,t}$ and $\arctan(\Delta s_y \,/\, \Delta s_x)$ returns the angle (in radians) in the interval $[-\pi, \pi]$[13]. By change of variables, the density of the movement to $\boldsymbol{s}_t$ is thus:

$$f(\boldsymbol{s}_t \mid \boldsymbol{s}_{t-1}) = f_{d,\phi}\left(\boldsymbol{g}\left((\Delta s_x, \Delta s_y)\right) \,\middle|\, \phi_{t-1}\right) \left|\det\left(J\left((\Delta s_x, \Delta s_y)\right)\right)\right| \qquad \text{eqn 7}$$

where the absolute determinant of the Jacobian $J$ is derived from $\boldsymbol{g}$ as $1/\sqrt{{\Delta s_x}^2 + {\Delta s_y}^2}$.

Incorporating the criterion that the individual can only swim in water completes the movement model:

$$f(\boldsymbol{s}_t \mid \boldsymbol{s}_{t-1}, \text{in water}) = \frac{1}{z(\boldsymbol{s}_{t-1})} f(\boldsymbol{s}_t \mid \boldsymbol{s}_{t-1})\, I(\boldsymbol{s}_t \text{ is in water}), \qquad \text{eqn 8}$$

where $I$ is the indicator function and $z$ is a normalisation constant[14].

**Step lengths.** We model $f_d(d)$ using two Cauchy distributions for 'low activity' (e.g., resting) and 'active' behavioural states, truncated between zero and mobility:

$$f_d(d) = \begin{cases} \text{Truncated Cauchy}(d;\ k_1, \theta_1, 0, \text{mobility}) \text{ if behaviour}_{t-1} = 0\ (\text{'low activity'}) \\ \text{Truncated Cauchy}(d;\ k_2, \theta_2, 0, \text{mobility}) \text{ if behaviour}_{t-1} = 1\ (\text{'active'}). \end{cases} \qquad \text{eqn 9}$$

The location ($k$) and scale ($\theta$) parameters determine the shape of the distribution and the mobility is the maximum moveable distance between $t-1$ and $t$ (i.e., the truncation threshold). The behaviour states were defined *a priori*[15]. Smaller values for $k_1$ and $\theta_1$, compared to $k_2$ and $\theta_2$, produce a low-activity step-length distribution that is more restrictive than that for active behaviour.

---

[13] arctan, or `atan` in our `Julia` code, corresponds to the standard definition of an `atan2` function in computer programming languages. This is a variation of the arctangent function that returns the angle between $-\pi$ and $\pi$, as opposed to the usual atan function that returns the angle between 0 and $\pi$ (or $-\pi/2$ and $\pi/2$). For further details, see the Julia documentation (Bezanson et al., 2017).

[14] In practice, the normalisation constant is obtained during model inference by Monte Carlo simulation (see Lavender et al. (2024a) and §4.4).

[15] As noted above, behavioural states were defined from depth (vertical activity) time series. The behaviour at time $t-1$ was assigned based on vertical activity between $t-1$ and $t$. During model inference (§4.4), at each $t-1$, we simulated movement into new location(s), given the behaviour over that timeframe (at $t-1$).

**Turning angles.** Turning angles were modelled using a Gaussian distribution:

$$f_{\Delta\phi}(\Delta\phi) = \text{Gaussian}\left(0.0, \sigma_{\Delta\phi}^2\right). \qquad \text{eqn 10}$$

Smaller $\sigma_\phi$ settings in this model produce smaller turning angles (i.e., more correlated random walks).

###### 4.1.3. Likelihood

The term $f(\boldsymbol{y}_{1:T} \mid \boldsymbol{s}_{1:T}) = \prod_{t=1}^{T} f(\boldsymbol{y}_t \mid \boldsymbol{s}_t)$ in eqn 1 is the joint likelihood; that is the probability of the observations given the latent locations. Our observations include acoustic detections (1) and non-detections (0) at each operational receiver $k$ (denoted $y_{k,t}^{(A)} \in \{0, 1\}$) and depth observations (denoted $y_t^{(D)}$). In the three versions of our model for the acoustic, depth and combined data types, $f(\boldsymbol{y}_t \mid \boldsymbol{s}_t)$ equals $f\left(\boldsymbol{y}_t^{(A)} \middle| \boldsymbol{s}_t\right)$, $f\left(y_t^{(D)} \middle| \boldsymbol{s}_t\right)$ and $f\left(\boldsymbol{y}_t^{(A)} \middle| \boldsymbol{s}_t\right) f\left(y_t^{(D)} \middle| \boldsymbol{s}_t\right)$ respectively.

**Acoustic observations.** The likelihood of acoustic observations was modelled as a Bernoulli probability mass function:

$$f\left(\boldsymbol{y}_t^{(A)} \middle| \boldsymbol{s}_t\right) = \prod_k \text{Bernoulli}\left(y_{k,t}^{(A)}; p_{k,t}(\boldsymbol{s}_t)\right), \qquad \text{eqn 11}$$

where detection probability is denoted $p$. We modelled $p$ as a truncated logistic function of the Euclidean distance $h(\boldsymbol{s}_t, \boldsymbol{r}_k)$ between the transmitter and the receiver ($\boldsymbol{r}_k = (s_{x;k}, s_{y;k})$):

$$p_{k,t}(\boldsymbol{s}_t) = \begin{cases} \left(1 + e^{-(\alpha - \beta \times h(\boldsymbol{s}_t, \boldsymbol{r}_k))}\right)^{-1} \text{ if } h(\boldsymbol{s}_t, \boldsymbol{r}_k) < \gamma \\ 0 \text{ otherwise} \end{cases}, \qquad \text{eqn 12}$$

where α and $\beta$ are parameters and $\gamma$ is the detection range.

**Depth observations.** The likelihood of the depth observation is modelled using a Gaussian distribution with a mean ($\mu_{\text{depth}}$) on the seabed $\left(b(\boldsymbol{s}_t)\right)$ and a standard deviation $\sigma_{\text{depth}}$, truncated between the surface ($\text{lim}_{\text{shallow}} = 0$ m) and the maximum depth ($\text{lim}_{\text{deep}} = 350$ m); i.e.,

$$f\left(y_t^{(D)} \middle| \boldsymbol{s}_t\right) = \text{Truncated Gaussian}\left(y_t^{(D)}; \mu_{\text{depth}}, \sigma_{\text{depth}}^2, \text{lim}_{\text{shallow}}, \text{lim}_{\text{deep}}\right). \qquad \text{eqn 13}$$

This model effectively elevates the probability of $y_t^{(D)}$ in latent locations where $y_t^{(D)}$ is close to the seabed, in line with skate's benthic lifestyle. The $\sigma_{\text{depth}}$ parameter accounts for observational and bathymetric uncertainty and tunes the probability density of pelagic movements off the seabed.

##### 4.2. Model parameterisation

We formulated and parameterised our state-space model using available information for flapper skate and other elasmobranchs[16]. In the following sections, the formulation and parameterisation of the movement model (§4.2.1) and acoustic (§4.2.2) and depth (§4.2.3) observation models are explained. For a summary of all model parameterisations, see Table S4.

###### 4.2.1. Movement model

The movement model (prior) represents how far skate can swim in a given time step. This model was based on a review of the literature on locomotory capacity (especially swimming speeds) in other elasmobranchs and limited available data from flapper skate.

###### 4.2.1.1. Literature review

**Elasmobranchs.** In the literature, studies of elasmobranch locomotion are relatively scarce and focus on sharks (Lauder and Di Santo, 2015). Routine swimming speeds are particularly poorly documented, but in sharks are generally thought to be around one body length per second ($BLs^{-1}$) (Parsons, 1990; Parsons and Carlson, 1998; Lauder and Di Santo, 2015). In shortfin mako (*Isurus oxyrinchus*), which is one of the fastest shark species, an accelerometery study documented mean sustained cruising speeds of 0.4 $BLs^{-1}$ or 0.9 $ms^{-1}$, comparable to those of other regionally endothermic fish (0.7–1.5 $ms^{-1}$) (Waller et al., 2023). The maximum burst swimming speed was 2.5 $BLs^{-1}$ (5.0 $ms^{-1}$), though speeds exceeding 4.9 $BLs^{-1}$ (9.8 $ms^{-1}$) have been documented during 'escape' responses and speeds up to 10 BL $s^{-1}$ (20 $ms^{-1}$) are thought possible (Motta et al., 2012).

[16] We perform model inference for the latent locations and use domain knowledge to parameterise the movement and observation sub-models. While it is possible to infer locations and static parameters simultaneously, the information available in passive acoustic telemetry systems is limited (Lavender et al., 2024a). Data driven estimation of static parameters is also computationally expensive, so we do not pursue this option here.

**Batoids.** In batoids, locomotion is based on oscillatory (‘flapping’) or undulatory (‘rippling’) movements of the enlarged pectoral fins (Lauder and Di Santo, 2015). Pelagic rays swim principally by oscillatory locomotion—a type of lift-based propulsion that is efficient for high-speed cruising. For oceanic manta rays (*Manta birostris*), median speeds observed from satellite tracking of six medium-sized individuals (disc length [DL] ≈ disc width ≈ 3.5–4.5 m) were 0.09 DLs$^{-1}$ (0.033 ms$^{-1}$) (Graham et al., 2012). Higher speeds of 0.5–1.0 DLs$^{-1}$ (2.78–4.17 ms$^{-1}$ or 10–15 kph) have been documented during breeding (Yano et al., 1999). A more recent study reported speeds ranging between approximately 0.29 DLs$^{-1}$ (during resting) to 0.75 DLs$^{-1}$ during feeding (Fong et al., 2022). Other studies on cownose rays (*Rhinoptera bonasus*) have reported comparable statistics, with migration speeds between 2.06–2.57 ms$^{-1}$ (Fish et al., 2017). Turning velocity estimates are somewhat lower and in oscillatory and undulatory rays range between 0.2–1.1 ms$^{-1}$ and 0.3–0.8 ms$^{-1}$ respectively (Parson et al., 2011).

**Benthic batoids.** In benthic batoids (including the majority of stingrays and skates), undulatory locomotion is more common and swimming efficiency is generally considered to be lower (Fish et al., 2017). Flow tank experiments have been conducted for blue-spot stingray (*Taeniura Lymma*, DL ≈ 0.16 m) and juvenile freshwater stingray (*Potamotrygon orbignyi,* DL ≈ 0.13 m) at steady swimming speeds of 0.9–3.0 DLs$^{-1}$ (0.22–0.55 ms$^{-1}$) (Rosenberger and Westneat, 1999) and 2.5 DLs (0.33 ms$^{-1}$) (Blevins and Lauder, 2012), respectively.

**Skate.** For skate (Rajidae), data are limited. Skate are often sedentary and small species, such as little skate (*Leucoraja erinacea*, DL ≈ 7 cm) may ‘punt’ over the seafloor (Koester and Spirito, 2003). Observations from trawl fisheries show that skate (and sharks) are often unresponsive to trawling or swim at slower speeds (< 1 ms$^{-1}$) (Queirolo et al., 2012). For little skate (which is between an undulator and oscillator), optimal cruising speeds are 1 DLs$^{-1}$ (0.07 ms$^{-1}$), although speeds of 1.25 DLs$^{-1}$ can be sustained for a few minutes and speeds of up to 2 DLs$^{-1}$ are documented over shorter time intervals (Di Santo et al., 2017). The extent to which these results scale to other species is unclear, but it is generally thought that in larger species size-relative routine speeds are lower (though absolute speeds may be higher). Trawl footage of species such as barndoor skate (*Dipturus laevis*) show that individuals may swim for a few seconds in front of a trawl, implying movement speeds up to 1.5 ms$^{-1}$ (3 knots) or approximately 1 DLs$^{-1}$, before trying to escape below the gear (Bayse et al., 2016). However, swimming capacity depends on context-specific conditions, including substrate hydrodynamic forces and temperature (Lauder and Di Santo, 2015).

**Flapper skate.** We used the literature on elasmobranch locomotion to bound our models of flapper skate movement in Scotland. In our dataset, captured skate vary in disc width from 0.75–1.75 m (which we take as an approximate measure of DL). For much of the time, flapper skate are probably largely sedentary on the seabed, resting or waiting to ambush benthic prey (Lavender et al., 2021a). Putative cruising speeds up to 1 DLs (0.75–1.75 $ms^{-1}$) are broadly comparable to pelagic rays and suggests speeds of 90–210 m per 2 min (the resolution of depth measurements). Speeds up to 2 $DLs^{-1}$ (1.5–3.5 $ms^{-1}$ or 180–420 m per 2 min) seem plausible. Over a few seconds, maximum burst speeds may exceed these limits but are likely less than the 3 $DLs^{-1}$ required to reach the speeds (2.25–5.25 $ms^{-1}$) achieved by big pelagic rays and endothermic sharks. However, substantially faster ground speeds may occur in areas of strong current flow. While average current speeds in our study system are negligible, speeds up to 4.75 $ms^{-1}$, capable of carrying skate 570 m per 2 min, have been documented (see §1.3).

**Datasets.** Little data exists with which to corroborate suggested flapper skate swimming speeds. Previous analyses of movements between receivers suggested potential speeds of up to 273 m per 2 min (2.27 $ms^{-1}$), which for larger skate is 1.30 $DLs^{-1}$, but likely underestimate attainable speeds. Ascent rates of up to 100 m per two minutes, and vertical movements up to 150 m per 2 min, are known (Lavender et al., 2021a, 2022b). Unpublished analyses of higher resolution depth time series produce similar numbers (James Thorburn, personal communication). Based on these results, in a previous, small-scale, illustrative analysis, we formulated a behavioural switching movement model in which movement probabilities during periods of high activity declined rapidly beyond 250 m up to a maximum mobility of 500 m (2.38 $DLs^{-1}$ for a 1.75 m skate) (Lavender et al., 2023). However, for the present study, this is likely overly restrictive.

###### 4.2.1.2. Parameter settings

The aforementioned literature, our earlier analyses and exploratory trials of the convergence properties of different model formulations, led to the formulation of a correlated random-walk movement model for flapper skate[17] (eqns 4–10, Figure S1). This model was parameterised as follows.

**Step lengths: best-guess parameterisation.** In our best-guess parameterisation, step lengths were modelled with the following parameters during 'low-activity' or 'active' behaviour:

$$\begin{cases} (k_1, \theta_1, \text{mobility}) = (0.0, 5.0, 1095.0) \\ (k_2, \theta_2, \text{mobility}) = (5.0, 100.0, 1095.0). \end{cases} \quad \text{eqn 14}$$

This parameterisation exhibits the following characteristics:

- **Low-activity distribution $(\boldsymbol{k_1}, \boldsymbol{\theta_1}, \textbf{mobility})$.** During 'low-activity' (including presumed resting periods), the model makes steps of zero length most likely, steps up to 25 m relatively plausible and longer steps unlikely[18].
- **Active distribution $(\boldsymbol{k_2}, \boldsymbol{\theta_2}, \textbf{mobility})$.** In the 'active' step-length distribution, longer steps are also less likely, especially beyond 1 $DLs^{-1}$ (210 m per 2 min for a large skate).
- **Truncation by mobility.** Both 'resting' and 'active' step-lengths distributions were constrained by the mobility parameter of 1,095 m per 2 min (the maximum moveable distance in 2 min). This value is based on (a) an absolute maximum swimming speed of 525 m per 2 min (2.5 $DLs^{-1}$ for a 1.75 m skate or 4.37 $ms^{-1}$) in (b) an area of high current flow (4.75 $ms^{-1}$). This model formulation makes slower movements more likely, while permitting faster movements with lower probability[19].

To examine the sensitivity of our analysis to our parameter choices, we also formulated a more restrictive model and a more flexible model.

---

[17] This model recognises two constraints. First, prior knowledge on flapper skate movements is limited. We therefore opted for a simple model structure that would effective exploration of the seascape during model inference (see §4.4). In our Bayesian framework, we expected inadequacies in this model to be at least partially corrected by the data. Second, model inference (via particle filtering plus smoothing) requires an evaluation of movement densities, which places some constraints on model flexibility (see §4.4). For our simple model, density evaluations are relatively straightforward and computationally feasible.

[18] During model inference (with particle algorithms), this encourages particles to remain in one place during resting behaviour while permitting occasional, longer distance movements (associated with other forms of low activity as well as fast, horizontal movements in flat regions of the study area), if required.

[19] As part of the inference procedure, movements into new locations are simulated from the movement model. Under our model, most simulated movement distances will be relatively small, but some larger movements may also be simulated. These movements are less likely, but may be 'selected' if required by the data (see §4.4). From a particle algorithm perspective, we note that the location and scale parameters of the step-length distribution are more important than mobility, providing mobility is large enough (Lavender et al., 2024a).

**Step lengths: restricted parameterisation.** In the restrictive parameterisation, parameter settings were as follows:

$$\begin{cases}(k_1, \theta_1, \text{mobility}) = (0.0, 5.0, 985.5) \\ (k_2, \theta_2, \text{mobility}) = (2.5, 50.0, 985.5),\end{cases} \qquad \text{eqn 15}$$

which represents a 50 % shrinkage in the location and scale parameters during active behaviour and a 10 % shrinkage in mobility (Figure S1).

**Step lengths: flexible parameterisation.** In the flexible parameterisation, parameter settings were corresponding inflated:

$$\begin{cases}(k_1, \theta_1, \text{mobility}) = (0.0, 5.0, 1204.5) \\ (k_2, \theta_2, \text{mobility}) = (7.5, 150.0, 1204.5),\end{cases} \qquad \text{eqn 16}$$

with a 50 % increase in the location and scale parameters during active behaviour and a 10 % increase in mobility (Figure S1).

**Turning angles.** All movement model parameterisations used a Gaussian distribution to represent turning angle, with a standard deviation of $\sigma_{\Delta\phi} = 1.5$. This corresponds to a weak–moderate correlation ($\rho \approx 0.32$) in heading[20].

###### 4.2.2. Acoustic observation model

**Model formulation.** The acoustic observation model was based on a truncated, distance-decaying, logistic detection-probability function[21] (Figure S1). Detection probability was set to decline with distance from a receiver at a rate determined by two coefficients ($\alpha$, $\beta$), truncated by the maximum detection range ($\gamma$) (Lavender et al., 2023, 2024a). This model formulation is based on evidence that the distance between a receiver and transmitter is the main driver of detection probability and can be parameterised using standard range-testing protocols, such as drift tests[22] (Kessel et al., 2014). We recognise that other variables, such as

[20] This choice was based on experience in our study area that weakly correlated random walks have improved convergence properties during model inference. Due to computational restrictions, it was not possible to trial more restrictive/flexible turning angle distributions in our sensitivity analyses (see §6–7). However, we expect our results are robust to limited changes in correlation strength. Moderate to strong correlations ($\sigma_{\phi} \leq 1.2$) induced convergence failures in initial experiments.

[21] During model inference with particle algorithms, this model effectively up-weights particles that are close to the receiver(s) that recorded detection(s) at the moment of detection and down-weights particles close to receivers in the gaps between detections.

[22] In particle algorithms, this model is also computationally cheap to evaluate.

noise, affect detection probability, but their effects are less generalisable and difficult to parameterise with available datasets (Lavender et al., 2021b).

**Parameterisation information.** We parameterised our detection-probability model using drift-test analyses (see §2.3), plus literature and manufacturer specifications:

- **Drift tests.** Drift tests demonstrated that probability declines to 0.5 by 400–450 m and is low beyond 700 m (see §2.3).
- **Literature.** While comparable data from other study systems are limited, detection ranges of 500–850 m for V13 tags and VR2W receivers are not uncommon in the literature (Selby et al., 2016; Winter et al., 2021) and align with manufacture guidelines. In deep-water marine ecosystems, ranges up to at least 1,628 m have been reported (Long et al., 2023) and in other systems tests of similar technologies have reported ranges exceeding 9–10 km (Hayden et al., 2016; Klinard et al., 2019).

**Best-guess parameterisation.** In our best-guess parameterisation, we set:

$$(\alpha, \beta, \gamma) = (4.0, -0.0094, 3000.0). \qquad \text{eqn 17}$$

This parameterisation exhibits the following features:

- **Half-detection probability.** The model approximately recapitulates drift-test results, exhibiting half detection probability at 425 m and low detection probability beyond 700 m.
- **Detection range.** The detection range (3000 m) is approximately twice the maximum range reported in other deep water systems (Long et al., 2023) but considerably lower than maximum known ranges for similar technology (Hayden et al., 2016; Klinard et al., 2019). In our algorithms, the detection range sets a hard threshold beyond which detections are assumed impossible[23]. The value of 3000 m is compromise that permits detections, with low probability, at moderate, but not excessive, distances from receivers.

**Restrictive parameterisation.** In our restrictive parameterisation, we set:

$$(\alpha, \beta, \gamma) = (3.0, -0.01175, 2250.0), \qquad \text{eqn 18}$$

[23] Therefore, the range must be sufficiently large to permit successful model inference (particle algorithm convergence), but is otherwise less important than function shape (since it affects the tail of the distribution) (Lavender et al., 2024a).

which represents a 25 % steepening of the function (25 % decrease in $\alpha$, 25 % increase in $\beta$ and 25 % decrease in $\gamma$).

**Flexible parameterisation.** In our flexible parameterisation, we set:

$$(\alpha, \beta, \gamma) = (5.0, -0.00705, 3750.0), \quad \text{eqn 19}$$

which represents a 25 % shallowing of the function.

###### 4.2.3. Depth observation model

**Initial model formulation.** We formulated a Gaussian depth observation model (eqn 13, Figure S1). During model inference, this model updates particle weights in line with the correspondence between the observed depth $\left(y_t^{(D)}\right)$ and the bathymetric depth $(b(\boldsymbol{s}))$. The mean of the Gaussian distribution ($\mu_{\text{depth}}$) was set to the depth of the seabed, in line with the benthic lifestyle of skate (Wearmouth and Sims, 2009; Thorburn et al., 2021; Lavender, 2022). Initially, the standard deviation ($\sigma_{\text{depth}} = 20$) was set accounting for accounting for uncertainty in the bathymetric depth (± 10 m, see §1.2.3), tidal elevation (± 3 m, see §1.3), storm surges and waves (± 2 m, see §1.3) and manufacturer-quoted tag accuracy (± 4.77 m, see §2.2) but later revised (see below).

**Revised model formulation.** In initial simulation experiments, we found that model inference with the above model was extremely difficult: in a complex bathymetric landscape, the model opens up a labyrinth of possible routes that particles have to explore but ultimately renders few compatible with the data. This makes convergence in the particle filter extremely hard to achieve—a challenge that can be compounded by mismatches between the underlying data-generating process and the modelled process. This is a known difficulty in particle filtering contexts and a hard problem to solve. In this context, our solution was to aggregate the bathymetry data onto a 500-by-500-pixel grid (as described in §1.2.2) and propagate additional uncertainty in bathymetry depth in the depth-observation model[24]. This makes it much easier for particles to move into cells that are compatible with the depth observations (since cells are

[24] We mention other possible solutions in the Discussion (see Main Text).

larger and encompass a larger range of depths), facilitating convergence (at the cost of reduced precision) in the particle filter[25].

**Best-guess parameterisation.** In our best-guess parameterisation, we set:

$$(\mu_{\text{depth}}, \sigma_{\text{depth}}, \text{lim}_{\text{shallow}}, \text{lim}_{\text{deep}}) = (b(\boldsymbol{s}), 100.0, 0.0, 350.0). \quad \text{eqn 20}$$

This parameterisation exhibits the following features:

- **Mean.** The model upweights locations for which the bathymetric depth is close to the observed depth.
- **Standard deviation.** The standard deviation accounts for known sources of uncertainty (± 20 m) and the range (± 70 m) of bathymetric values in each cell[26], while permitting potential pelagic movements (Wheeler, 1969; Brown-Vuillemin et al., 2020), and effectively 'dampens' the influence of depth observations sufficiently to achieve convergence in most instances[27] (see Main Text Results).

**Restrictive parameterisation.** In the restrictive parameterisation, we trialled:

$$(\mu_{\text{depth}}, \sigma_{\text{depth}}, \text{lim}_{\text{shallow}}, \text{lim}_{\text{deep}}) = (b(\boldsymbol{s}), 50.0, 0.0, 350.0), \quad \text{eqn 21}$$

which represents a 50 % reduction in the standard deviation (i.e., reduced uncertainty, pelagic behaviour and/or dampening).

**Flexible parameterisation.** In the flexible parameterisation, we set:

$$(\mu_{\text{depth}}, \sigma_{\text{depth}}, \text{lim}_{\text{shallow}}, \text{lim}_{\text{deep}}) = (b(\boldsymbol{s}), 150.0, 0.0, 350.0), \quad \text{eqn 22}$$

which represents a 50 % increase in the standard deviation (i.e., increased uncertainty, pelagic behaviour and dampening).

##### 4.3. Model uncertainty

[25] Although the aggregation of the bathymetry data and the 'widening' of the depth observation model reduce precision, in our simulations even the DC algorithm (which only incorporates depth observations) outperformed heuristic methods (based on acoustic detections and tuning parameters) on average. Among the particle algorithms, the integration of depth observations also refined maps of space use (see §6 and Main Text Results). The aggregation of the bathymetry grid additionally improved computation time (in selected routines) and substantially reduced memory and disk-space requirements.

[26] The range in bathymetric depths was ≤ 70 m within 95 % of aggregated grid cells.

[27] While flapper skate are considered predominately benthic, some pelagic behaviour is expected based on diet analyses (Wheeler, 1969; Brown-Vuillemin et al., 2020). However, the degree to which flapper skate behave pelagically is unknown. Further information on the vertical movements of skate relative to the seabed would help improve the precision of the depth-observation model.

We recognise that our observation models are imperfect and uncertain[28]. We therefore examine algorithm sensitivity to variation in parameter values (within reasonable limits). By implementing the AC, AC and ACDC algorithms, we also reveal the extent to which independent datasets suggest similar patterns of space use and the extent to which patterns based only on acoustic or depth data are refined by the integration of both datasets.

##### 4.4. Model inference

The particle algorithms perform model inference for our state-space model (§4.1), given static parameters are specified (see §4.2). Particle algorithms were implemented using the `patter` package (Lavender, 2024; Lavender et al., 2024b):

- **Filter.** The particle filter was implemented with the `pf_filter()` function[29]. The computational complexity of this routine is $O(TN)$, where $T$ is the number of time steps and $N$ is the number of particles (Doucet and Johansen, 2009). The filter stores a subset of equally weighted particles at each time step[30].
- **Smoother.** Particle smoothing was performed using the two-filter smoother implemented by `pf_smoother_two_filter()`. This algorithm re-weights particles from a forward filter run and a backward filter run in line with the probability density of movements between pairs of locations (Doucet and Johansen, 2009). One hundred Monte Carlo simulations were used to compute the normalisation constant for the density of the movement model (see eqn 8).

In the particle filter, we used 100,000–150,000 particles[31]. In the particle smoother, we used 1,500–2,000 particles (see §6–7). The number of particles is a trade-off between the fidelity by

[28] That being said, even mis-specified state-space models can consistently outperform heuristic algorithms, where all data-generating processes must be enveloped by one or a small number of 'tuning' parameters (Lavender et al., 2024a).
[29] See the Main Text for an intuitive, algorithmic explanation of the particle filter. In this study, we implemented the movement model iteratively until a valid movement (on the bathy-low raster) was found or for a maximum number of 10,000 trials. Particles which failed to make a valid move in 10,000 trials were killed. Low-variance resampling was implemented every time step.
[30] The equal weighting is achieved by low-variance resampling.
[31] In the particle filter, large numbers of particles may be required to facilitate convergence (see Main Text). Fortunately, this is feasible because the computational complexity of the filter is only $O(TN)$. This differs from smoothing, where a smaller number of particles are used because (a) we only need sufficient particles to approximate (in this case) a two-dimensional distribution and (b) there is a much greater penalty for every particle $(O(TN^2))$. Computational time was an important constraint in our study because we analysed simulated and observed datasets for many individuals using multiple parameter settings. Note that with insufficient smoothing particles the two filters may not align at every time step (that is, there may be no valid moves between the subset

which the (marginal) distribution of an individual's location ($f(\boldsymbol{s}_t \mid \boldsymbol{y}_{1:T})$) is approximated and computational complexity, which is $\mathcal{O}(TN^2)$ for smoothing (Doucet and Johansen, 2009). UDs were estimated from particle samples as described in §5. In selected analyses, the expected time spent in selected areas (i.e., residency) was estimated from the proportion of (smoothed) particles in those areas[32].

#### 5. Mapping

##### 5.1. Kernel smoothing

For simulated paths (see §6.1) and coordinates estimated by the COA and particle algorithms (see §6–7), we generated maps of space use using kernel smoothing (Lavender et al., 2024b). For mapping, coordinates were summarised on our 500-by-500-pixel grid (bathy-low) in terms of probability-of-use ($P$), which is the probability that an individual is located in a particular cell at a randomly chosen time; i.e.,

$$P_I \propto \sum_{i,t} \delta_{I,I_{i,t}} w_{i,t}, \qquad \text{eqn 23}$$

where $I$ indexes grid cells, $i$ indexes coordinates, $\delta$ is the Kronecker delta and $w$ denotes weights[33] (Lavender et al., 2024b).

For the COA algorithm, $P_I$ amounts to the proportion of COAs in cell $I$, since only one COA is estimated per time interval and equal weights are assumed. For the particle algorithms, $P_I$ accounts for the number of copies of each particle within a cell (within and across time steps).

Smooth maps of space use were generated by kernel smoothing the summarised (weighted) coordinates, with edge correction[34]. For speed, kernel bandwidth was defined from the combined variance of summarised coordinates using the ad-hoc method (Worton, 1989).

---

of retained particles on the backward filter and those from the forward filter). In this instance, `patter` randomly retains 50 % of the particles from the forward filter and 50 % from the backward filter. In this study, we only analysed outputs for which proper smoothing was possible on >90 % of time steps.

[32] Recall that in our implementation of the filter/smoother, particles are resampled prior to storage. The residency estimate is technically a weighted proportion of particles (but, in this case, particle weights are incorporated via resampling).

[33] To avoid confusion, as above we re-iterate that stored particles are equally weighted.

[34] For edge correction, we used an observation window defined by the Westminster constituencies shapefile from Digimap (see §1.2.1). The shapefile was used (rather than our bathy-low pixel image) for computational efficiency. The window was simplified via the `simplify.owin()` function in the `spatstat.geom` package, using a tolerance (`dmin`) of 1,000 m (Baddeley and Turner, 2005). This heuristic maintained an accurate representation of the coastline while substantially speeding up UD estimation (from approximately 161 → 10 s).

Kernel smoothing was implemented via the `patter map_dens()` function, which wraps `density.ppp()` from `spatstat.explore` (Baddeley and Turner, 2005).

##### 5.2. Brownian bridges

For the RSP algorithm, a dynamic Brownian-bridge movement model is used to smooth coordinates (Niella et al., 2020). One map is estimated for each set of detections spaced less than one day apart. We implemented the method, as defined in the `RSP` package, using an enlarged bathymetric raster to mitigate estimation errors (see §1.2.1). For each dataset, the overall map was defined as the re-normalised aggregation of all maps for that dataset (resampled onto our 500-by-500-pixel [bathy-low] grid).

#### 6. Simulation analyses

##### 6.1. Data simulation

We performed simulation analyses to investigate algorithm behaviour (performance and sensitivity) and inform the implementation and interpretation of real-world analyses (see Table S2 for the overview and §7 for details).

The simulation analysis was based on 100 hypothetical movement paths (and corresponding observational datasets), simulated over a one-month period ($T = 21{,}600$ two-minute time steps)[35]. Paths were simulated as follows.

**Initialisation.** Each path was initialised randomly in a location $(s_{x,t=1}, s_{y,t=1})$ within the receiver array ($A$), with a random heading ($\phi_t$):

$$\begin{aligned}(s_{x,t=1}, s_{y,t=1}) &\sim \text{Uniform}(s_x, s_y; A)\\ \phi_{t=1} &\sim \text{Uniform}(-\pi, \pi).\end{aligned} \qquad \text{eqn 24}$$

**Behavioural states.** Subsequently, movements were simulated in discrete-time following our best-guess behavioural switching correlated random walk model (see §4.2.1). Low-activity and

[35] The two-minute time resolution was chosen to align with the time resolution of real-world analyses, which incorporated depth observations measured every two minutes (see §2.2 and §7).

active behavioural states were simulated based on observed properties of real depth time series. Following Lavender et al. (2022), we distinguished 'low activity' (0) and 'active' (1) behaviour in depth time series based on a vertical activity threshold of 0.25 m per 2 min. We observed that individuals spent approximately 50 % of time in each behavioural state and, in our simulations, sampled an initial behaviour from a discrete uniform distribution accordingly:

$$\text{behaviour}_{i=1} \sim \text{Uniform}(\{0,1\}). \qquad \text{eqn 25}$$

Subsequently, for each $\text{behaviour}_i$, we iteratively simulated blocks (indexed by $i$) of low-activity or active behaviour that were maintained for $k_i$ time steps. The duration for which behaviours were maintained was based on simple Gamma (shape, rate) models derived from the observed time series:

$$k_i \sim \begin{cases} \text{round}\big(\text{Gamma}(0.730, 0.163)\big) \text{ if behaviour}_i = 0 \\ \text{round}\big(\text{Gamma}(0.960,0.168)\big) \text{ if behaviour}_i = 1. \end{cases} \qquad \text{eqn 26}$$

Each $\text{behaviour}_i$ thus lasts for $k_i$ time steps (with different $\text{behaviour}_i$ generally lasting for different durations). After each block of low-activity or active behaviour, a behavioural transition occurred:

$$\text{behaviour}_{i+1} = 1 - \text{behaviour}_i \qquad \text{eqn 27}$$

Blocks of behaviours were simulated iteratively for the duration of the timeline, that is, while $\sum_{i=1} k_i \leq T$. These blocks were then unwrapped into a time series of behavioural states, such that each time step $t$ was associated with a single behavioural state. The time series of behavioural states were used in the next stage of the simulation.

**Movements.** Following initialisation, for each subsequent time step, movements were simulated by sampling from our best-guess flapper skate model (eqns 4–10). Step lengths were modelled using truncated Cauchy distributions (with location [$k_1$ and $k_2$], scale [$\theta_1$ and $\theta_2$] and truncation parameters [0.0, mobility])[36]:

$$f_d(d_t) = \begin{cases} \text{Truncated Cauchy}(d_t;\ 0.0, 5.0, 0.0, 1095.0) \quad \text{if behaviour}_{t-1} = 0 \\ \text{Truncated Cauchy}(d_t; 5.0, 100.0, 0.0, 1095.0) \text{ if behaviour}_{t-1} = 1. \end{cases} \qquad \text{eqn 28}$$

Turning angles were modelled using the Gaussian distribution:

$$f_{\Delta\phi}(\Delta\phi_t) = \text{Gaussian}(\Delta\phi_t;\ 0.0, 1.5). \qquad \text{eqn 29}$$

[36] As explained previously, these two models enable us to capture two behavioural states: a 'low activity' state with predominantly small steps and an 'active' state with larger steps on average.

Patterns of space use for the simulated path were generated by kernel smoothing (see §5).

**Observations.** For each movement path, we collected state vectors $\boldsymbol{s}_t = (s_{x,t}, s_{y,t}, \phi_t, \text{behaviour}_t)$ and simulated acoustic and depth observations, following the aforementioned data-generating processes (eqns 11–13 in §4.2) with our best-guess parameters (see eqns 17 and 20 in §4.2). That is, acoustic observations were sampled from a Bernoulli model:

$$\boldsymbol{y}_t^{(A)} \sim f\left(\boldsymbol{y}_t^{(A)} \mid \boldsymbol{s}_t\right)$$

$$f\left(\boldsymbol{y}_t^{(A)} \mid \boldsymbol{s}_t\right) = \prod_k \text{Bernoulli}\left(y_{k,t}^{(A)}; p_{k,t}(\boldsymbol{s}_t)\right) \quad \text{eqn 30}$$

$$p_{k,t}(\boldsymbol{s}_t) = \begin{cases} \left(1 + e^{-(4.0 - 0.0094 \times h(s_t, r_k))}\right)^{-1} & \text{if } h(\boldsymbol{s}_t, \boldsymbol{r}_k) < 3000 \text{ m} \\ 0 \text{ otherwise,} & \end{cases}$$

where $\boldsymbol{r}_k$ denotes the location for receiver $k \in \{1, 2, \ldots, 58\}$[37].

Depth observations were sampled from a Gaussian model with a mean $\mu_{\text{depth}} = b(\boldsymbol{s}_t)$ and a standard deviation $\sigma_{\text{depth}} = 100.0$, truncated between 0 and 350 m:

$$y_t^{(D)} \sim f\left(y_t^{(D)} \mid \boldsymbol{s}_t\right)$$

$$f\left(y_t^{(D)} \mid \boldsymbol{s}_t\right) = \text{Truncated Gaussian}\left(y_t^{(D)}; b(\boldsymbol{s}_t), 100.0, 0.0, 350.0\right). \quad \text{eqn 31}$$

#### 6.2. Performance analyses (A1)

In performance analyses, we evaluated the skill with which algorithms reconstructed simulated patterns of space use and residency. See Table S2 for the overview.

**A1.1. Algorithm parameterisation.** Algorithms were parameterised with optimal or best-guess parameters.

[37] In simulations, we used real receiver locations but imagined that all receivers were active in the one-month period over which data were simulated. We expected algorithms based on acoustic observations to perform better in simulations than in real-world analyses (§7), as in reality only a subset of receivers was deployed in any one month (and we accounted for this in real-world analyses).

**A1.1.1. Heuristic algorithms.** For heuristic algorithms, a simulation approach was used to select optimal parameter values for $\Delta T$ (COA) and `er.ad` (RSP)[38] (see Main Text and Figure S3).

**A1.1.2. Particle algorithms.** Particle algorithms were parameterised according to the data-generating processes (eqns 28–31).

**A1.2. Algorithm implementation.** Heuristic algorithms were implemented for each acoustic time series and particle algorithms (AC, DC and ACDC) were implemented for each (a) acoustic, (b) depth and (c) combined dataset, respectively[39]. We initialised particle algorithms at the simulated capture or recapture location (on the forward and backward filter runs, respectively). We used a 'capture container' to shrink the domain within which particles were resampled towards the simulated 'finishing' (capture or recapture) location[40]. Acoustic containers were similarly used for AC and ACDC algorithm runs to facilitate convergence[41] (Lavender et al., 2024a). In the particle filter, we used 100,000 particles. For smoothing, 1500 particles were used[42] (see §4.4). UDs were reconstructed as described in §5. For each algorithm, we checked the proportion of simulations for which UDs were successfully estimated.

**A1.3. Algorithm performance.** We evaluated algorithm performance by the accuracy with which patterns of space use and residency were estimated.

---

[38] As a reminder: for each path, we computed the mean error (ME) between the UD for each simulated path and reconstructed UDs, based on a range of parameter values, and identified the parameter value with the lowest ME on average (see Main Text Methods). For the COA algorithm, ME was minimised at $\Delta T = 2$ days (notwithstanding variation). For RSPs, ME weakly declined with increasing `er.ad` values, but there was no winning choice of `er.ad` on average. The setting `er.ad` $= 500$ m was taken as the optimum. This value is sufficiently large to reduce ME but not so large as to substantially increase computation time or fail (due to limited domain size). See Main Text Results and Figure S3.

[39] See §3 and §4.4 for generic heuristic and particle algorithm implementation details.

[40] The capture container was implemented as an additional 'observation model'. At each time step, we computed the probability of each particle given the distance of the particle from the 'finishing' location. Particles must be within radius = mobility + mobility × $\Delta T$ of the 'finishing' location (where mobility is the maximum moveable distance in one time step and $\Delta T$ is the number of time steps until the capture/recapture event). We used the extra mobility buffer to relax the requirement for particles to finish exactly in the correct location, instead permitting particles anywhere within mobility m of the correct location (that was also compatible with the simulated data).

[41] Acoustic containers work similarly to capture containers but kill particles that are incompatible with the location(s) of the next acoustic detection(s), given the detection range, rather than particles which are incompatible with a known 'finishing' location (Lavender et al., 2024a).

[42] We expected 1,500 particles to be sufficient to approximate the two-dimensional distribution $f(\boldsymbol{s}_t \mid \boldsymbol{y}_{1:T})$. In practice, we achieved relatively low effective sample sizes (ESS) with 1,500 particles. (For 'performance' analyses, the mean ESS across all simulations was 195 particles.) We also observed one situation in which the forward and backward filters were insufficiently aligned for adequate smoothing (see §4.4). For this reason, we boosted the number of particles in real-world analyses (see §7).

**A1.3.1.** ***Maps of space use: visual evaluation.*** We visually evaluated algorithm performance for the first three paths by comparing simulated and reconstructed UDs.

**A1.3.2.** ***Maps of space use: quantitative evaluation.*** To examine algorithm performance in terms of patterns of space use for all simulated paths, we computed and visualised the distribution of mean absolute error between simulated UDs, the UD for a null model[43] and UDs from the heuristic and particle algorithms[44].

**A1.3.3.** ***Residency***. We also examined algorithm performance in terms of the accuracy with which residency (within zones open or closed to fisheries and the MPA as a whole) was estimated. Residency for the simulated path was defined as the proportion of time steps spent in each zone. Residency for the null model amounts to the proportional area of each zone in the study (marine) domain. This was compared against a standard residency index (detection days) as well as residency estimated from the heuristic and particle algorithms. 'Detection days' were defined as the proportion of days with detections at receivers in each zone[45]. Residency for heuristic and particle algorithms was estimated as explained previously (see §3 and §4.4). We then evaluated algorithm performance in terms of the distribution of residency error (the difference between the estimated and true residency), across all paths, by algorithm/zone.

#### 6.3. Sensitivity analyses (A2)

In sensitivity analyses, we selected a subset of simulated datasets (1–3) for which to compare alternative algorithm parameterisations and thus evaluate algorithm sensitivity. See Table S2 for the overview.

**A2.1. Algorithm implementation.** We re-implemented all algorithms with 'restrictive' and 'flexible' settings. For COAs and RSPs, we restricted and inflated the $\Delta T \in \{2 \text{ days}, 1 \text{ day}, 3 \text{ days}\}$ and $\text{er.ad} \in \{500, 250, 750\}$ parameters, on the basis of the aforementioned simulation analysis of heuristic parameters (A1.1.1). For particle algorithms, we trialled restrictive and flexible parameterisations of the movement model

[43] The null model was a flat probability distribution spanning the marine habitats within the study area (defined by bathy-low).

[44] We used the mean error as a metric of model skill as a previous study showed this metric effectively distinguishes between the performance of our algorithms (Lavender et al., 2024a).

[45] All receivers were located in closed zones within the MPA, so we only computed one estimate of residency according to detection days that was identical in closed areas and the MPA as a whole. Detection days cannot be used to estimate residency in regions without receivers.

(see §4.2.1.2), acoustic observation model (see §4.2.2) and/or depth observation model (see §4.2.3), while holding other parameters constant (at data-generating settings). Each best-guess, restrictive and flexible parametrisation of the particle algorithms was implemented three times to examine algorithm reproducibility (given inference via particle algorithms is based on a stochastic sequential Monte Carlo procedure). Other settings, such as the number of particles, were fixed as in performance analyses (A1). Convergence for each algorithm parameterisation was tracked, as explained above (A1.2).

##### A2.2. Algorithm sensitivity and reproductivity.

**A2.2.1.** ***Maps of space use: visual examination.*** As in performance analyses (A1), we visualised UDs for the three selected paths for each algorithm parameterisation. For the particle algorithms, we also visualised UDs for a selected path (1) for the three independent runs of each algorithm parameterisation to examine reproducibility of maps of space use.

**A2.2.2.** ***Maps of space use: quantitative evaluation.*** We computed and visualised mean absolute error between UDs for each simulated path and the reconstructed UD from each algorithm parameterisation. As in performance analyses (A1), this analysis included UDs from the null model as well as UDs from the heuristic and particle algorithms.

**A2.2.3.** ***Residency***. We computed and visualised residency error for each path and algorithm parameterisation, as in performance analyses (A1).

#### 7. Real-world analyses

##### 7.1. Datasets

In our real-world analysis, we analysed patterns of space use from tagged flapper skate, using acoustic data (from 33 individuals) and archival data (from 21 individuals) made available by previous studies (Figure S2).

We considered three datasets:

- The acoustic data
- The archival data
- The combined data

Datasets were processed as follows:

- **Blocking.** Each dataset was split into individual/month datasets.
- **Individuals.** We focused on individuals/months with both sets of observations (to facilitate comparisons between AC, DC and ACDC algorithm outputs).
- **Inclusion criteria.** Datasets were retained only for individuals that were at liberty for the entire month (ensuring comparability among maps of space use) and for individuals that were detected in at least two different weeks on a total of seven days in a given month. This criterion excluded individuals with the sparsest time series (i.e., individuals that were detected following tagging and then appeared to leave the array).

##### 7.2. Analyses (A3–4)

For each individual/month dataset, we reconstructed patterns of space use and estimated residency using the COA, RSP and particle algorithms (see Table S2 for the overview). For each dataset, we implemented algorithms with optimal or best-guess parameters (A3) alongside restrictive and flexible algorithm parameterisations to quantify sensitivity (A4). The COA and RSP algorithms were implemented exactly as in simulation analyses (see §6). Particle algorithm implementations were adjusted to account for the features of the real-world data (see §7.2.1–2).

##### 7.2.1. Particle algorithm initialisation

Unlike simulation analyses, particle algorithms could not be initialised at known capture/recapture locations (since individuals were not captured/recaptured exactly at the start/end of each individual–month dataset).

Initially, we trialled initialising the filter in locations sampled uniformly from the part of the study area compatible with the data[46]. However, this approach performed poorly: more precise initial locations were required to achieve convergence with moderate numbers of particles[47].

To improve initialisation in the particle filter, for each dataset, we used a simulation approach:

- **Geolocation records.** We identified the nearest known geolocations of that individual (to the start and end of each time series), based on the full detection time series and the Skatespotter mark-recapture angling database[48] in which all known capture events (individual, date, location) are recorded[49]. This database generally only records capture dates, but we identified the exact time of capture events by visual examination of the raw depth time series[50] (Lavender et al., 2022b).
- **Movement simulation.** From each geolocation record, we simulated 30 realisations of our best-guess movement model over the duration between that record and the 'start' of the filter timeline ($t = 1$ or $t = T$ for the forward and backward filters, respectively).

[46] For example, we initialised the AC algorithm with particles sampled from within an 'acoustic container' around the receiver(s) that recorded the closest detection(s) in time to $t = 1$ (or $t = T$ on the backward filter run). This container defines the region within which the individual must have been located at $t = 1$. For instance, if an individual was first detected at $t = 5$, then at $t = 1$ we sampled locations in a container around that receiver of radius $= \gamma + 4 \times$ mobility. Automated sampling from the regions compatible with the data is the default initialisation option in `patter` (Lavender, 2024; Lavender et al., 2024b).

[47] Precise initial locations greatly facilitate convergence in the particle filter. In the example of the AC algorithm, the issue in our case is that mobility is large (relative to the size of the study area) and the container within which particles are sampled quickly becomes large as the time to the most recent detection(s) increases. Most particles are thus initialised in unlikely locations and are killed, leading to poor convergence properties.

[48] The Skatespotter database is hosted at https://skatespotter.sams.ac.uk. This database is based on voluntary submissions from charter skippers and recreational angling, which is common in our study area (Lavender et al., 2022a, 2022b). Skate caught by anglers can be identified from external tags, passive integrated transponder tags and/or photographs (Benjamins et al., 2018).

[49] This approach enables geolocation records beyond the one-month timeline directly accessible to the filter to refine filter initialisation.

[50] Following tagging, individuals rapidly descend (by >100 m) to the seabed; recapture events are visible as 'spikes' in the depth time series, as individuals are pulled to the surface and then released; and archival tag recovery events are similarly visible from rapid ascents (by > 100 m) as skate are pulled towards the surface. The exact coordinates of capture events are not always recorded exactly, but the vast majority of (recorded) angling in our study area occurs at known angling marks (Lavender et al., 2022b), which made it possible to assign capture locations with sufficient accuracy for this analysis.

The simulation of movement trajectories followed the workflow described previously (see §6.1).

- **Initialisation container.** At the start of the filter timeline, we assumed the individual was located in a 'container' around the geolocation record with a radius equal to (a) two thirds of the maximum detection range (2000 m) or (b) the maximum distance of simulated positions from the geolocation record (whichever was larger). For gaps exceeding 12 hours in duration, we added an additional 5000 m buffer (in acknowledgement of greater uncertainty in individual movements over longer periods).
- **Initialisation samples.** We further assumed that at the start of the filter timeline, individuals must be in a location on the high-resolution bathymetric grid (bathy-high) within 20 m of the observed depth, accounting for accounting for uncertainty in the bathymetric depth (± 10 m, see §1.2.3), tidal elevation (± 3 m, see §1.3), storm surges (± 2 m, see §1.3) and tag accuracy (± 4.77 m, see §2.2). We sampled initial locations uniformly, with replacement, from the set of possible locations on bathy-high within the initialisation container. This process was repeated for the start and end of each time series (for the forward and backward filter runs, respectively).

###### 7.2.2. Particle algorithm implementation

Particle algorithms were implemented for the acoustic (AC), depth (DC) and combined (ACDC) datasets.  Each algorithm run for a given individual/month was initialised with the same (best-guess) starting locations, since precise initialisation was important to achieve convergence. Behavioural states were assigned *a priori* to time steps, from the depth (vertical activity) time series, using the threshold 0.25 m (see also §6.1). Similar to simulation analyses (§6.2), all particle filters included an additional 'capture container' observation model structure to facilitate consistency between starting and finishing locations in the forward and backward filter runs. We used this structure to shrink the 'container' within which particles were located at each time step towards the centroid of the set of possible 'ending' locations[51]. (In the case of

[51] At the end of the forward filter, we assumed the individual must be within radius $=$ mobility $+$ $\max_j \left( \left\| \left(s_x^{(c)}, s_y^{(c)}\right) - \left(s_{x_{t=T}}^{(j)}, s_{y_{t=T}}^{(j)}\right) \right\| \right)$ of the centroid $\left(s_x^{(c)}, s_x^{(c)}\right)$ of all $j \in \{1, 2, \ldots, J\}$ possible ending locations $\left(s_{x_{t=T}}^{(j)}, s_{y_{t=T}}^{(j)}\right)$. Programmatically, at relevant time steps in the filter, we computed the distance of each particle to $\left(s_x^{(c)}, s_y^{(c)}\right)$ and killed particles that exceeded the radius for that time step. This helped to pull particles from the forward filter into the region spanned by particles used to initialise the backward filter (and vice versa). This is important for consistency between filter runs and for particle smoothing, which resamples particles in line with

the forward filter run, the ‘ending’ locations are used to initialise the backward filter; in the case of the backward run, the ‘ending’ locations are those used to initialise the forward filter.) Otherwise, particle algorithms proceeded as previously described (see §4.4). We used 150,000 particles for filtering and 2,000 particles for smoothing[52] (see §6.2).

the probability density of movements between locations on the two filter runs (which requires at least some particles from the two filter runs to be within mobility of each other).

[52] As in simulation analyses (see §6), these choices represent a compromise between performance and computation time. The number of particles for filtering appeared to be generally sufficient, as we observed high convergence of the filters (see Main Text Results). A larger number of particles for smoothing would have helped to improve alignment between the two filters for smoothing (which was achieved for 75 % of filter runs), but was not possible given the computational resources available for this study. For best-guess analyses, the mean effective sample size per time step across all algorithm runs was 311 particles (AC: 250, DC: 349, ACDC: 385).

**Supporting figures**

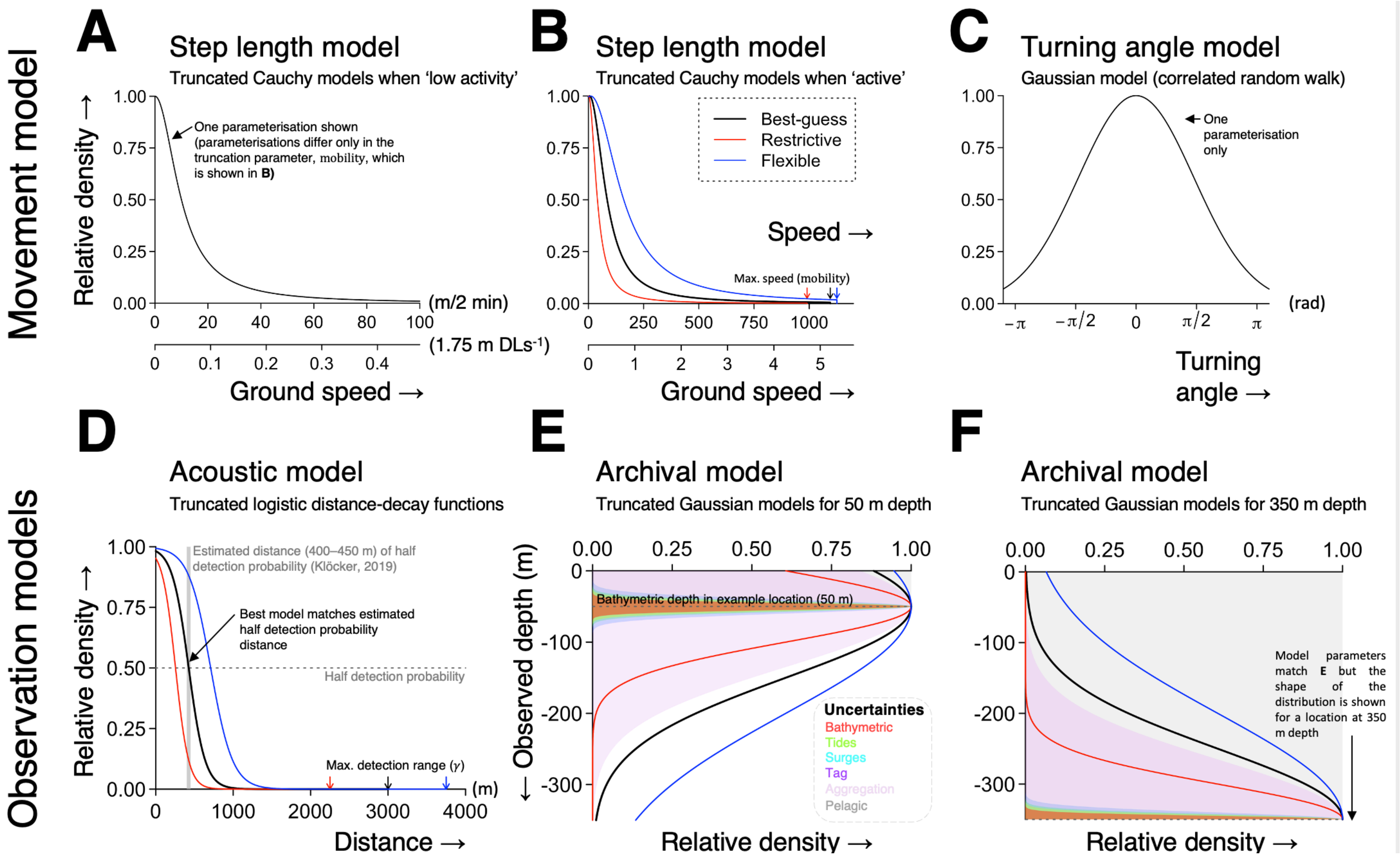

**Figure S1. A state-space model for tracking flapper skate. A–C** illustrate the movement model; that is, the truncated Cauchy distribution of step lengths during (**A**) 'low activity' and (**B**) 'active' behavioural states and (**C**) the Gaussian distribution of turning angles. Step lengths are shown in units of metres per 2 min and disc lengths (DL) per second for a 1.75 m DL individual[53]. **D–F** show the observation models for (**D**) acoustic and (**E–F**) depth observations at two bathymetric depths. The observation models evaluate the probability of the latent locations given (**D**) detection or non-detection at each operational receiver (the acoustic model) and (**E–F**) the depth observation (the depth model)[54]. The acoustic model is based on a truncated logistic, distance-delaying, detection-probability function. The depth model weighs the correspondence between a latent location's bathymetric depth and the observed depth. The required correspondence accounts for uncertainties in the bathymetric depth measurement, tidal range, storm surges, tag accuracy, bathymetry aggregation and potential pelagic movement (these are cumulative and shown by the coloured bands in **E** and **F**). Best-guess models are shown by the thick lines. Restricted and flexible model parameterisations (for the step length and observation models) are shown by the coloured lines and were used to examine algorithm sensitivity. All densities are scaled to a maximum value of one for visualisation. For full details, see Supporting Information §4. For parameter values, see Table S4.

[53] Panels **A–C** represent our correlated-random walk model of the movement process in flapper skate: we assume two behavioural states; a 'low-activity' state when movement is limited and an 'active' state, during which time an individual is more mobile. The Gaussian distribution of turning angles leads to a weakly correlated random walk.
[54] In simple terms, the observation models create a bridge between our model of the underlying movement process and our imperfect observations of that process. In a state-space model, we combine a movement model with the observations to infer the probability distribution of possible locations for an individual.

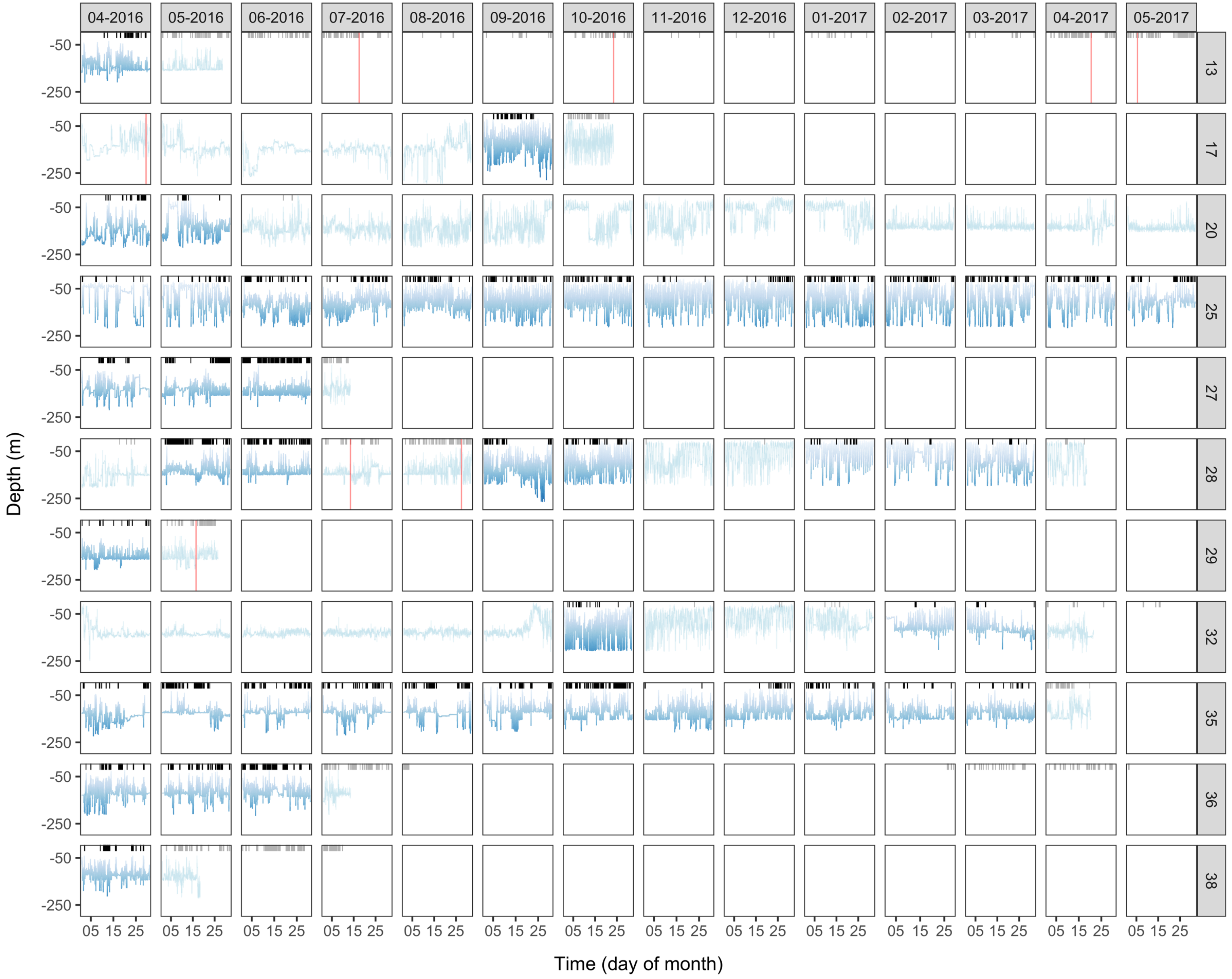
04-2016
05-2016
06-2016
07-2016
08-2016
09-2016
10-2016
11-2016
12-2016
01-2017
02-2017
03-2017
04-2017
05-2017
13
17
20
25
27
28
29
32
35
36
38
-50
-250
Depth (m)
05 15 25
Time (day of month)

**Figure S2. Observed time series.** Panels distinguish individuals (rows) and months (columns) for which acoustic and archival (depth) data were collected from April 2016—May 2017. Points mark detections and lines mark depth observations. The colour scheme for depths is defined with respect to the bathymetry in the study area, following Figure 1. For each individual, we modelled all months (panels) for which the individual was fully at liberty, with sufficient detection data (i.e., detections in at least two 7-day periods) and complete (uninterrupted) depth time series ($N = 48$ panels) (see Supporting Information §7.1). Transparent panels indicate time series that did not meet the criteria for modelling. Recapture events, which interrupted individuals' time at liberty, are marked by red vertical lines. For a summary of modelled individuals, see Table S5.

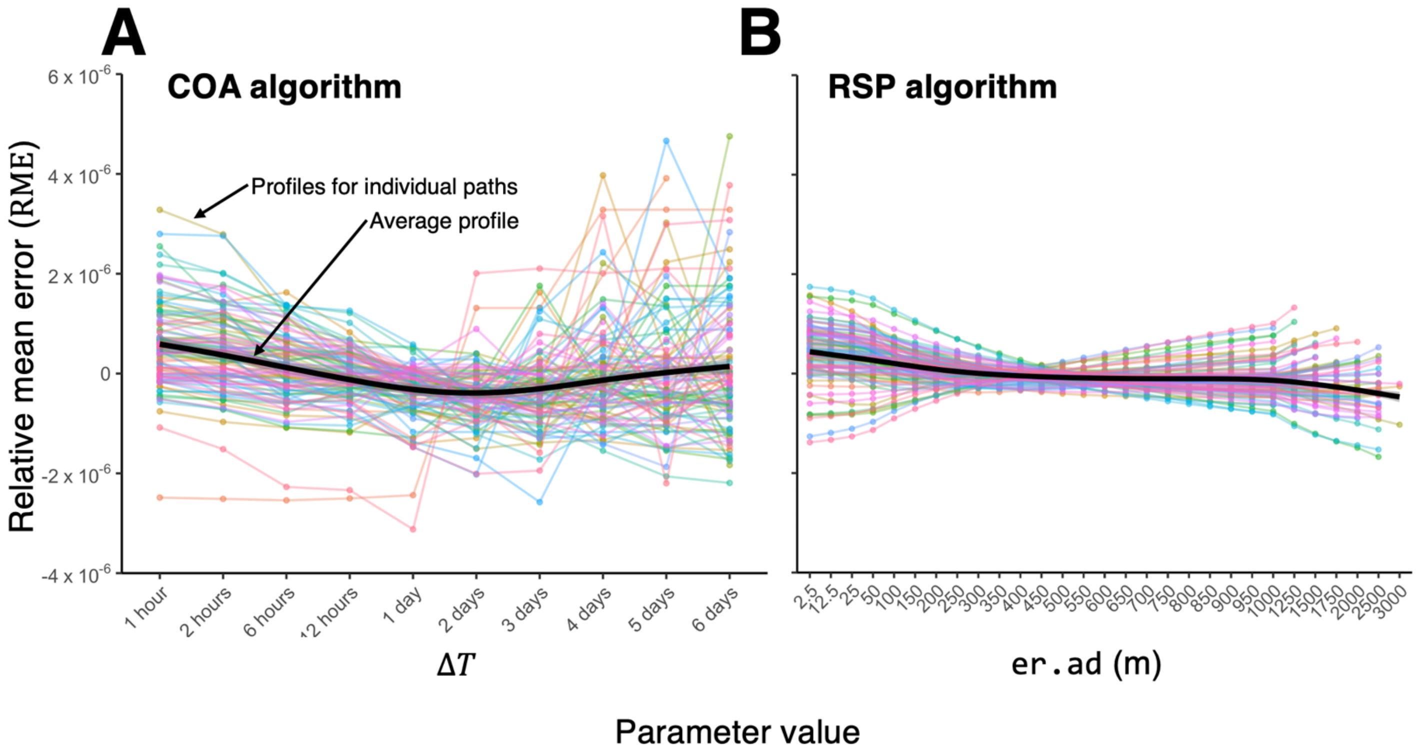


**Figure S3. Simulation analysis: heuristic algorithm performance profiles for (A) COAs and (B) RSPs.** For each simulated path ($i \in \{1, \ldots, 100\}$), the trend in relative mean absolute error (RME) across different values of $\Delta T$ (the time interval over which COAs are calculated) or `er.ad` (the rate at which uncertainty increases along RSPs) is shown. RME is the mean absolute difference between the UD for a simulated path and the corresponding reconstructed UD, centred by the average ME across all parameter values for that path. Note that the x-axes are non-linear. In each panel, the smooth line shows the expected RME as a function of the parameter value. Smooths were derived from generalised additive models of $RME_i \sim s(x_i)$, where $s$ is a thin plate regression spline, fitted by `mgcv` (Wood, 2017). They are shown for visual aid only. We used these profiles to select parameter values for real-world analyses.

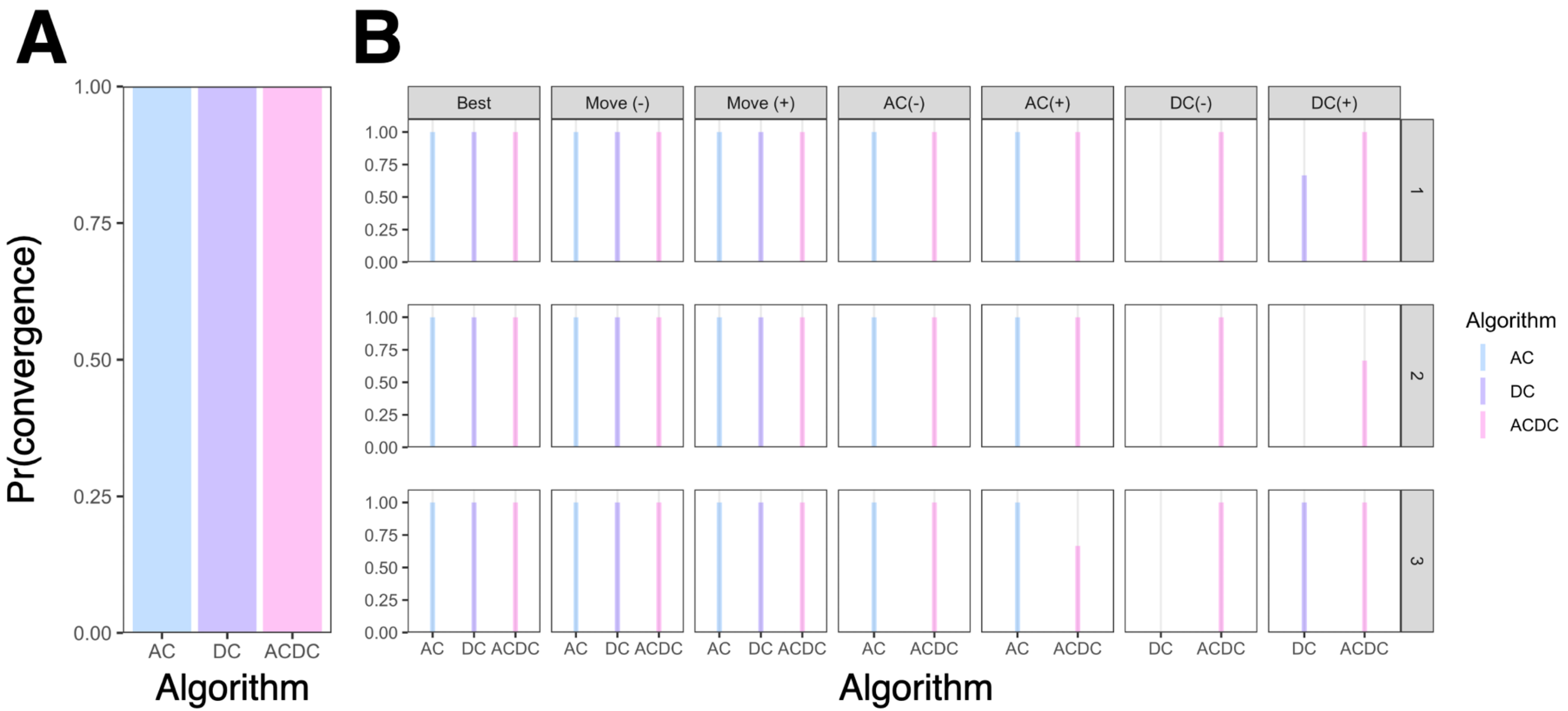


**Figure S4**. **Simulation analysis: particle algorithm convergence properties for (A) performance and (B) sensitivity analyses. A** shows the proportion of algorithm runs that converged in the performance analysis. In this analysis, model inference was performed once for each of the 100 simulated datasets using the data-generating parameter values. **B** shows the proportion of algorithm iterations that converged in the sensitivity analysis. In this analysis, model inference was performed for each of first three simulated paths (rows) three times, using best-guess (data-generating) and mis-specified parameter values (columns). Mis-specified algorithm implementations were characterised by under (-) or over (+) estimation of the movement (Move), acoustic (AC) and depth (DC) observation model parameters, with other components held constant at data-generating values.

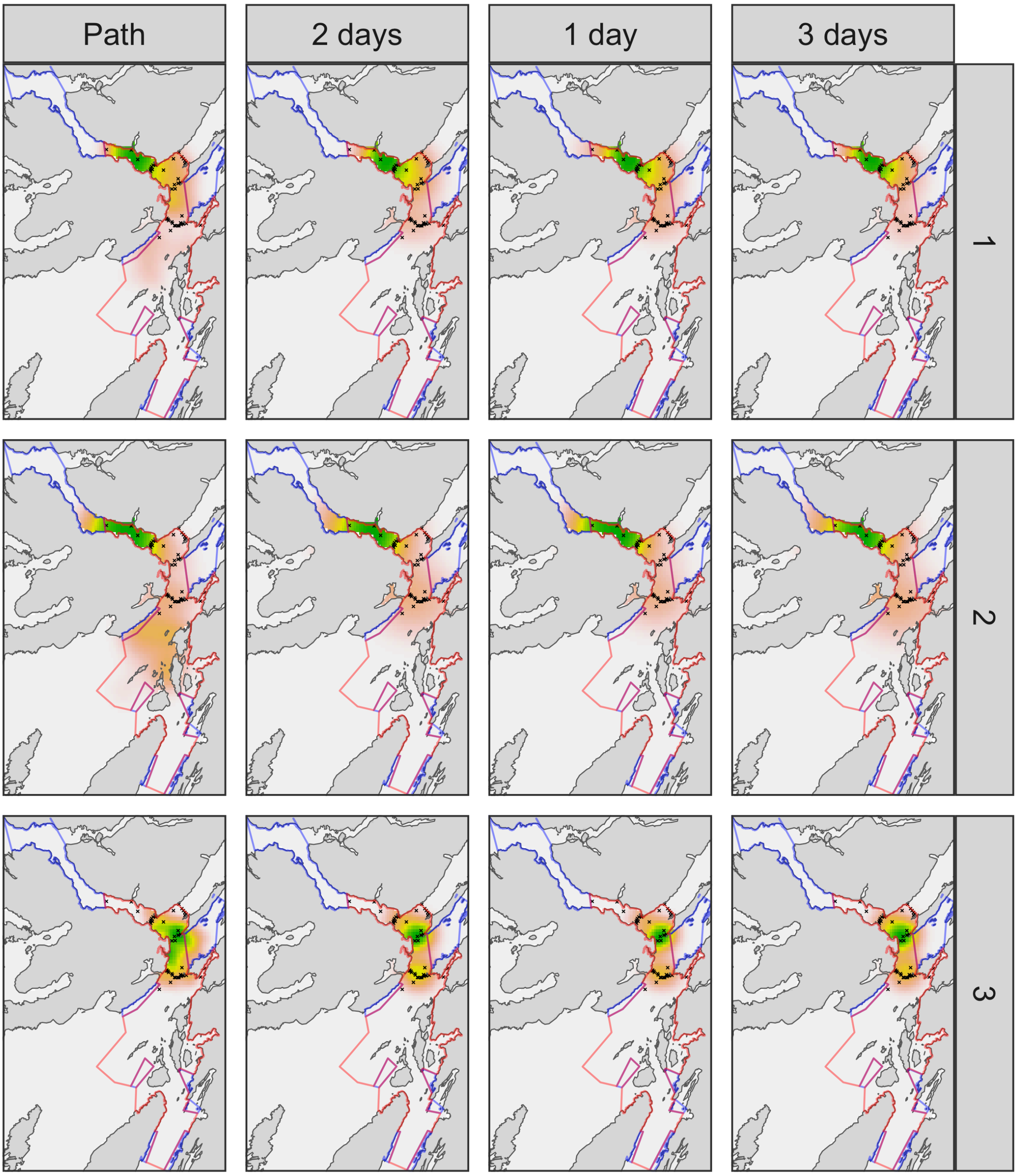


**Figure S5. Simulation analysis: COA algorithm performance and sensitivity.** Rows distinguish simulated paths. For each simulated path, the 'true' UD and the UD from three COA algorithm implementations with different $\Delta T$ values (optimal, restrictive, flexible) are shown. Crosses mark receivers. The colour scale represents probability density and is shown between zero (white) and the maximum density (green) on each panel (following Figures 3–4). For clarity, maps are only shown within the extent of the MPA, as delineated by the blue and red polygons. These enclose protected zones open and closed to fisheries, respectively.

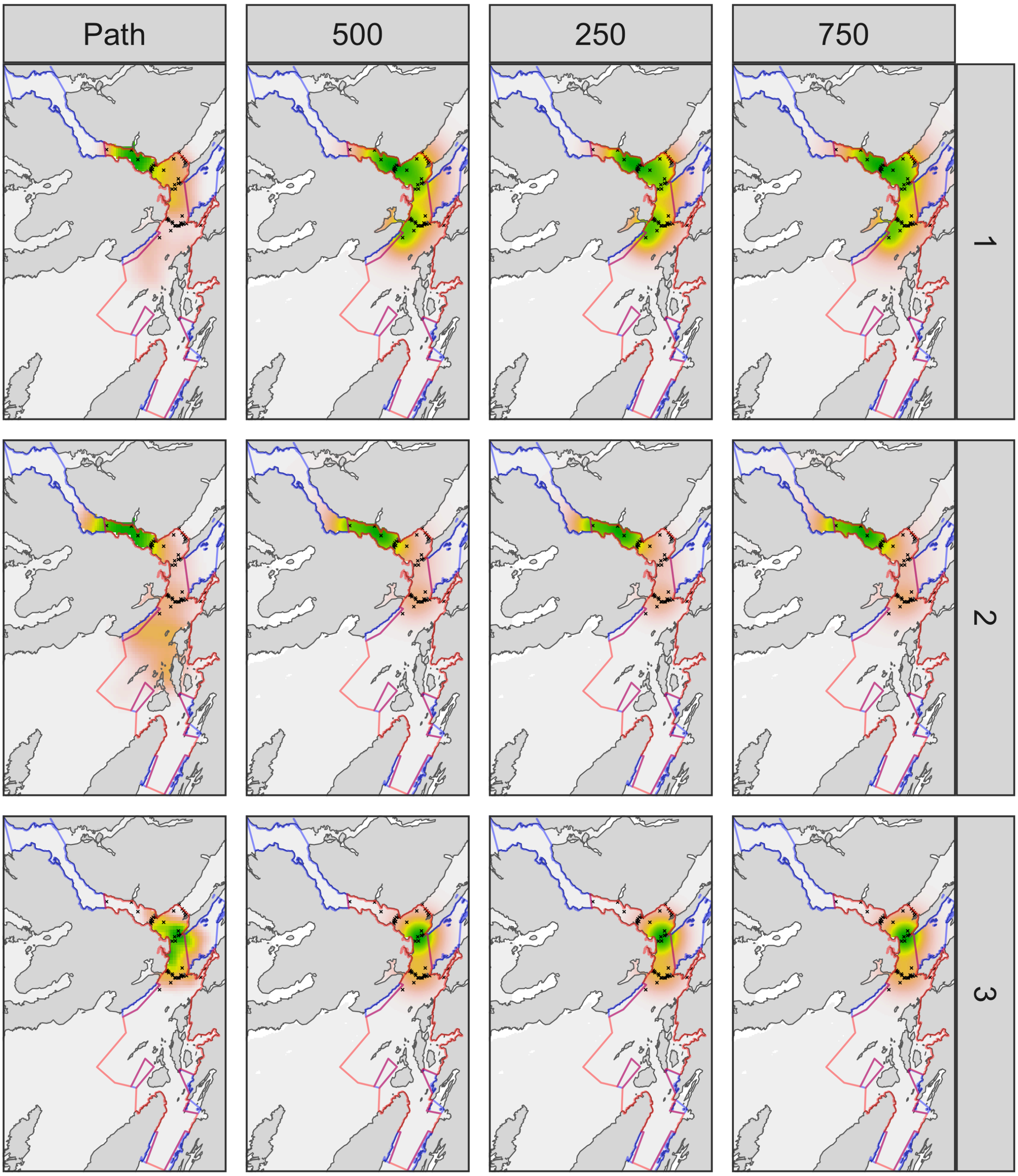


**Figure S6. Simulation analysis: RSP algorithm performance and sensitivity, following Figure S5.**

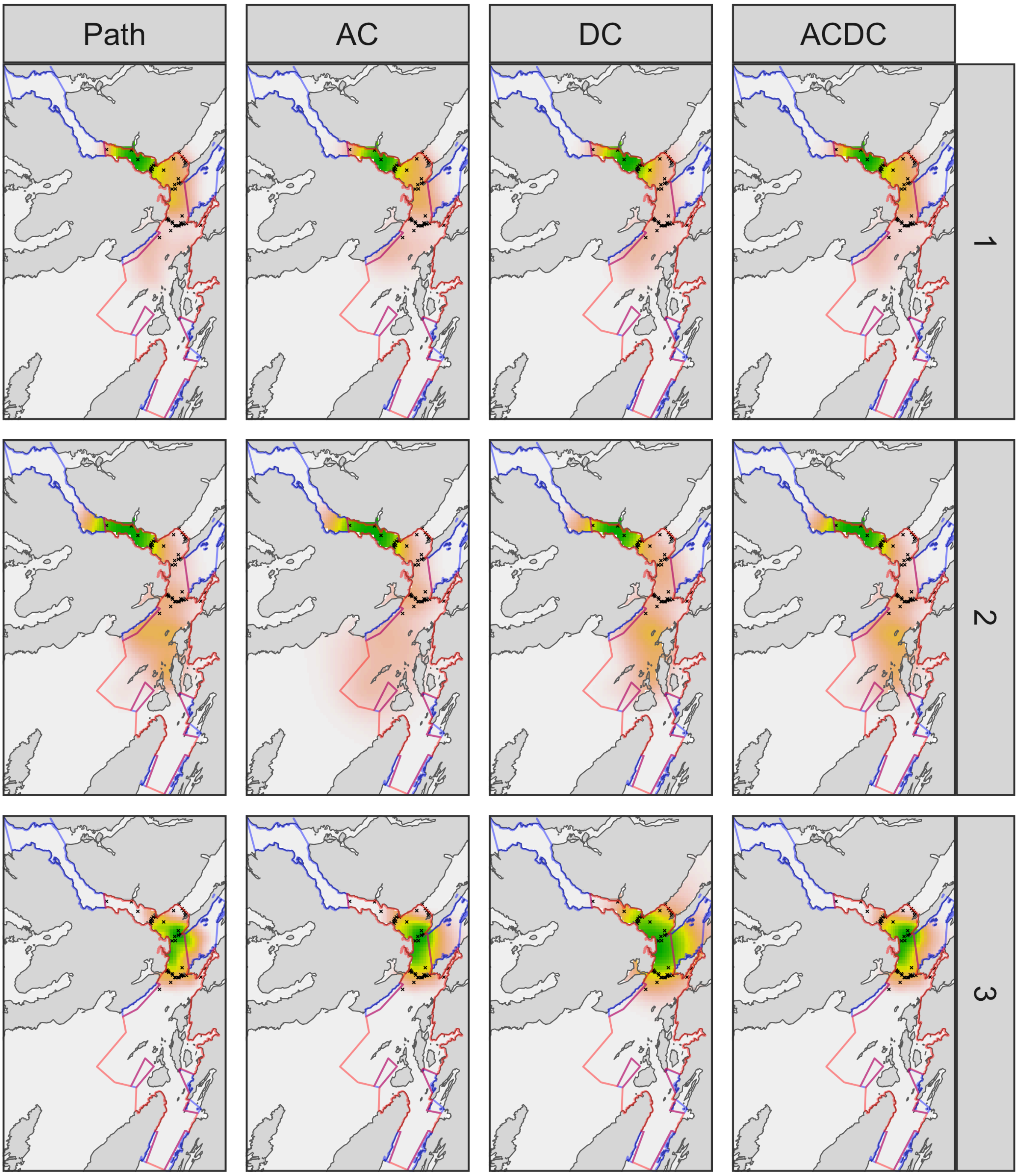


**Figure S7. Simulation analysis: particle algorithm performance, following Figure S5.** Rows distinguish simulated paths. For each simulated path, the 'true' UD and the UD from three particle algorithms (AC, DC and ACDC), correctly parameterised according to the data-generating process, are shown.

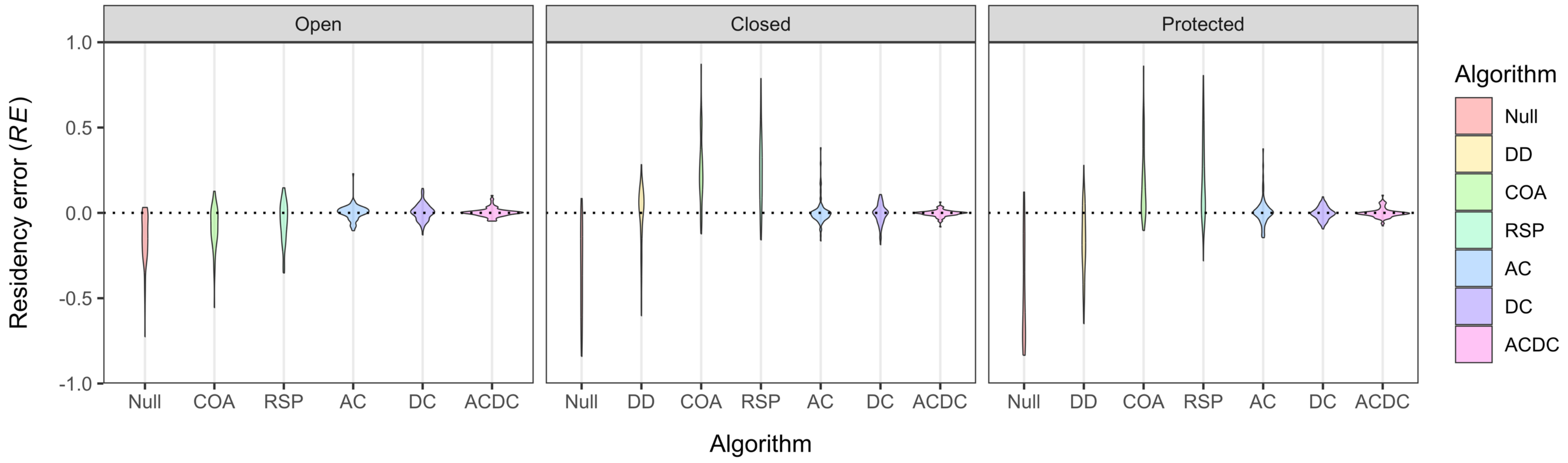


**Figure S8. Simulation analysis: model performance for estimating residency**. Panels show residency error for zones within the MPA that are (a) open or (b) closed to fisheries and (c) the MPA as a whole. Residency error is the difference between estimated residency (the proportion of time spent in each zone) and to the true value. Positive/negative values indicate overestimation/underestimation, respectively. Residency estimates were derived from performance analyses in which patterns of space use were reconstructed for each of the 100 simulated paths using optimal or data-generating parameter values. A null model and detection days (DD) are shown for context. The null model corresponds to the proportion of the study area in each zone. Detection days is the proportion of days with detections at receiver(s) (all of which are in closed areas). For heuristic (COA and RSP) and particle (AC, DC, ACDC) algorithms, residency was estimated (a) as the proportion of a UD's volume or (b) the proportion of particles, in each area, respectively. Each 'violin' shows the distribution of residency between the UD for a simulated path and the UD reconstructed by the corresponding algorithm, across all 100 simulated paths. In performance analyses, model inference was performed once for each of the 100 simulated datasets using heuristic algorithms with optimal parameters and particle algorithms (AC, DC and ACDC) with correctly defined (data-generating) parameters. Violin area is scaled by the number of algorithm runs that converged.

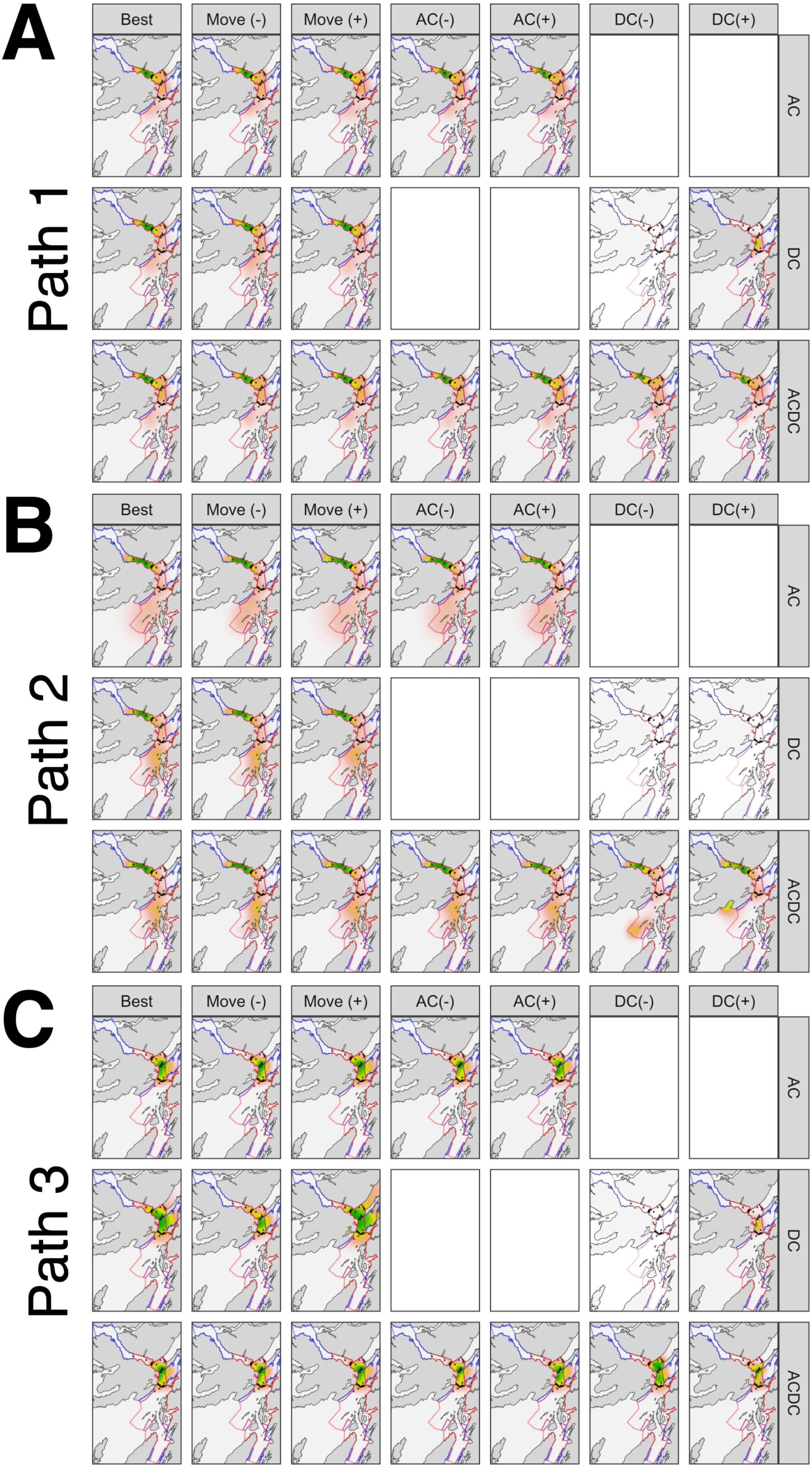
A
Path 1
Best
Move (-)
Move (+)
AC(-)
AC(+)
DC(-)
DC(+)
AC
DC
ACDC
B
Path 2
C
Path 3

**Figure S9. Simulation analysis: particle algorithm sensitivity.** For each simulated path (1 [**A**], 2 [**B**] or 3 [**C**]), for each algorithm (row), UDs are shown for seven algorithm runs with different parameter settings (columns). UDs are shown for the first algorithm iteration (out of three for each parameter setting). Transparent panels indicate convergence failures. Blank panels indicate nonsensical parameter combinations (such as AC algorithm runs with under/overestimated depth observation model parameters).

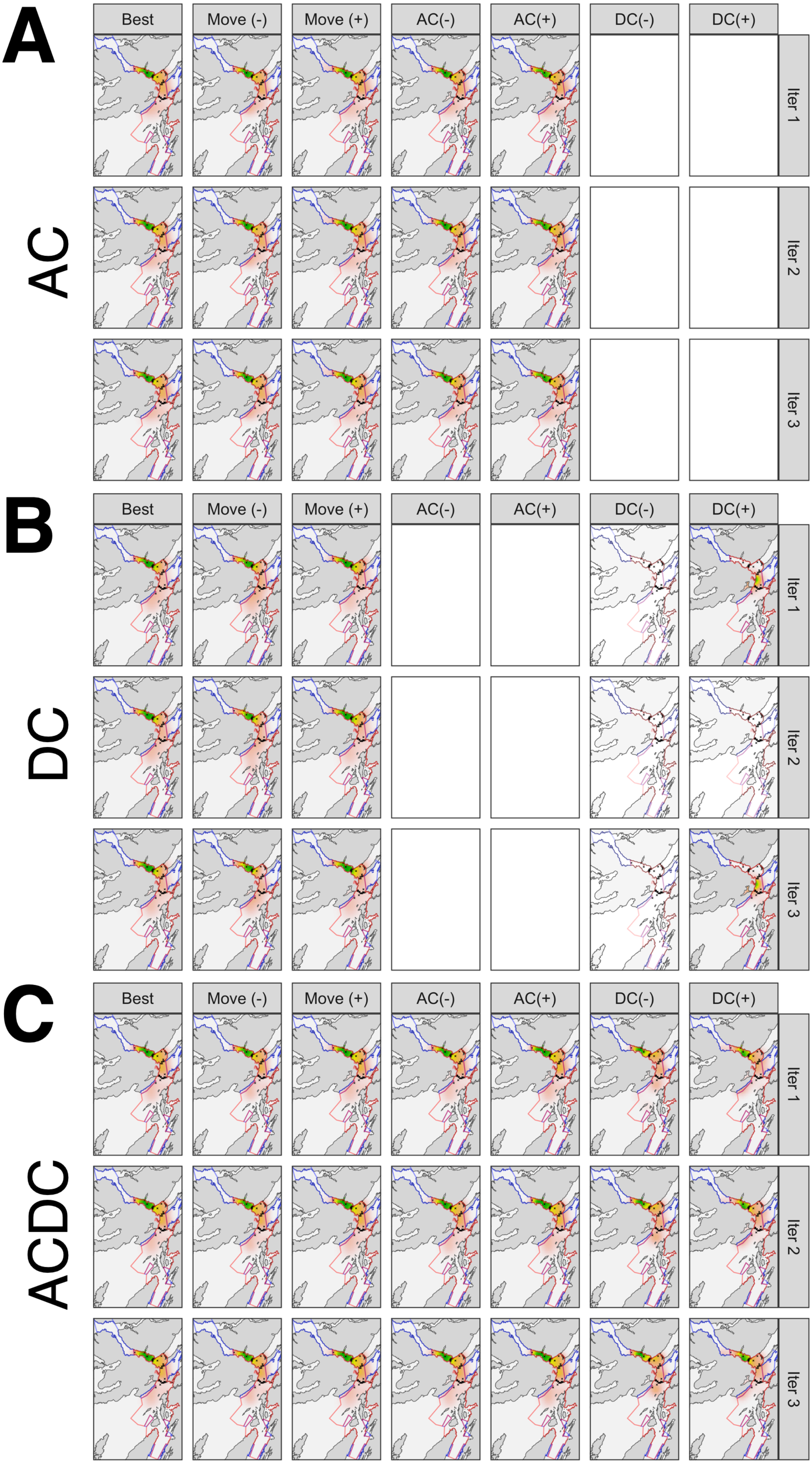
A
AC
Best
Move (-)
Move (+)
AC(-)
AC(+)
DC(-)
DC(+)
Iter 1
Iter 2
Iter 3
B
DC
C
ACDC

**Figure S10. Simulation analysis: particle algorithm repeatability, following Figure S9.** For the first simulated path, for each algorithm (row grouping), for three algorithm iterations (rows), UDs are shown for the seven parameter combinations (columns). Transparent panels indicate convergence failures and blank panels indicate nonsensical parameter combinations.

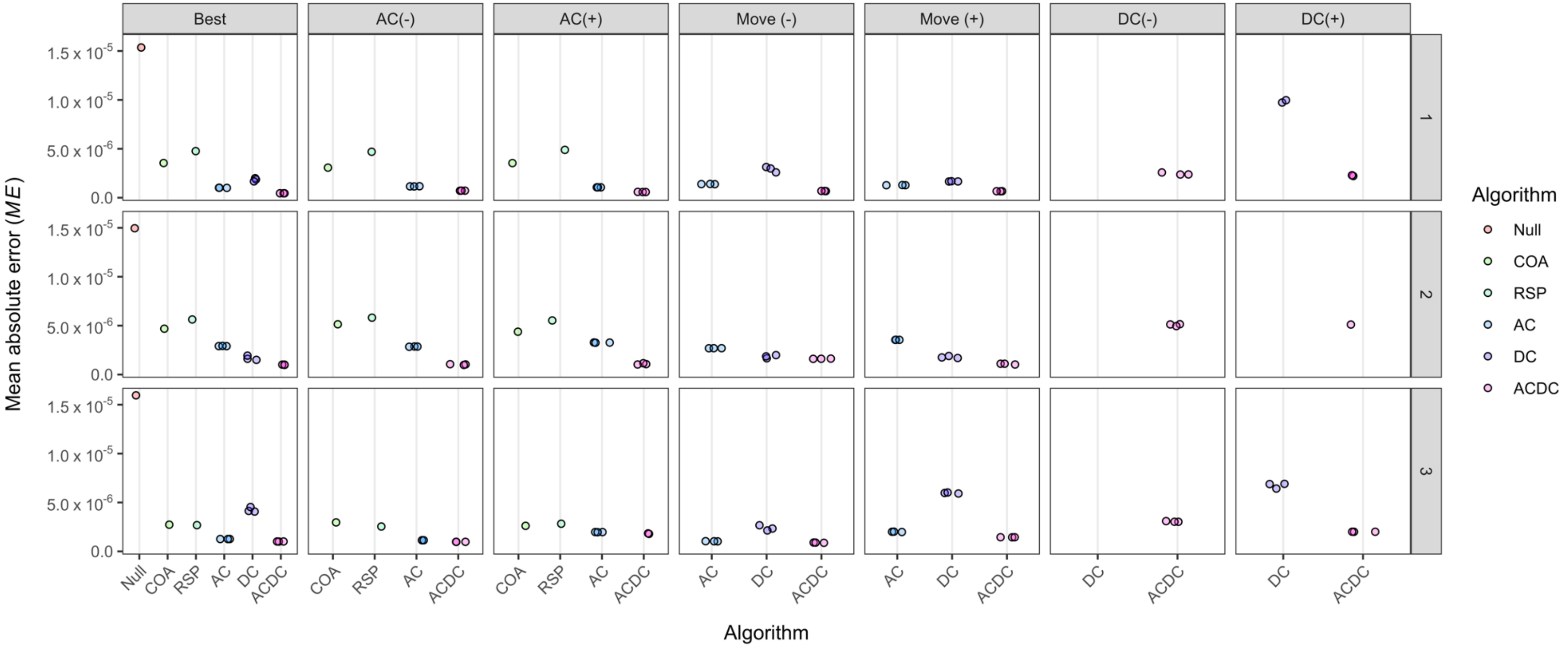


**Figure S11. Simulation analysis: algorithm sensitivity for mapping space use.** Each panel shows the mean absolute error between the UD for a simulated path (rows) and the UD reconstructed by a selected algorithm parameterisation (column), including optimal/best-guess and restrictive/flexible parameterisations (columns). The null model is shown in the 'best' panel for reference. Particle (AC, DC and ACDC) algorithm implementations included a best (data-generating) parameterisation alongside under (-) or over (+) estimation of the movement (Move), acoustic (AC) and depth (DC) observation model parameters, with other components held constant at data-generating values. Particle algorithms were implemented independently three times to examine reproducibility (and three points are shown for each setting, except in the case of convergence failures). The two heuristic (COA and RSP) algorithms were implemented with optimal values (in 'best') and restricted/flexible parameterisations that are nominally included in the AC(-) and AC(+) panel columns, since these algorithms focus exclusively on acoustic detections. These algorithms are deterministic and only a single point is shown in relevant panels.

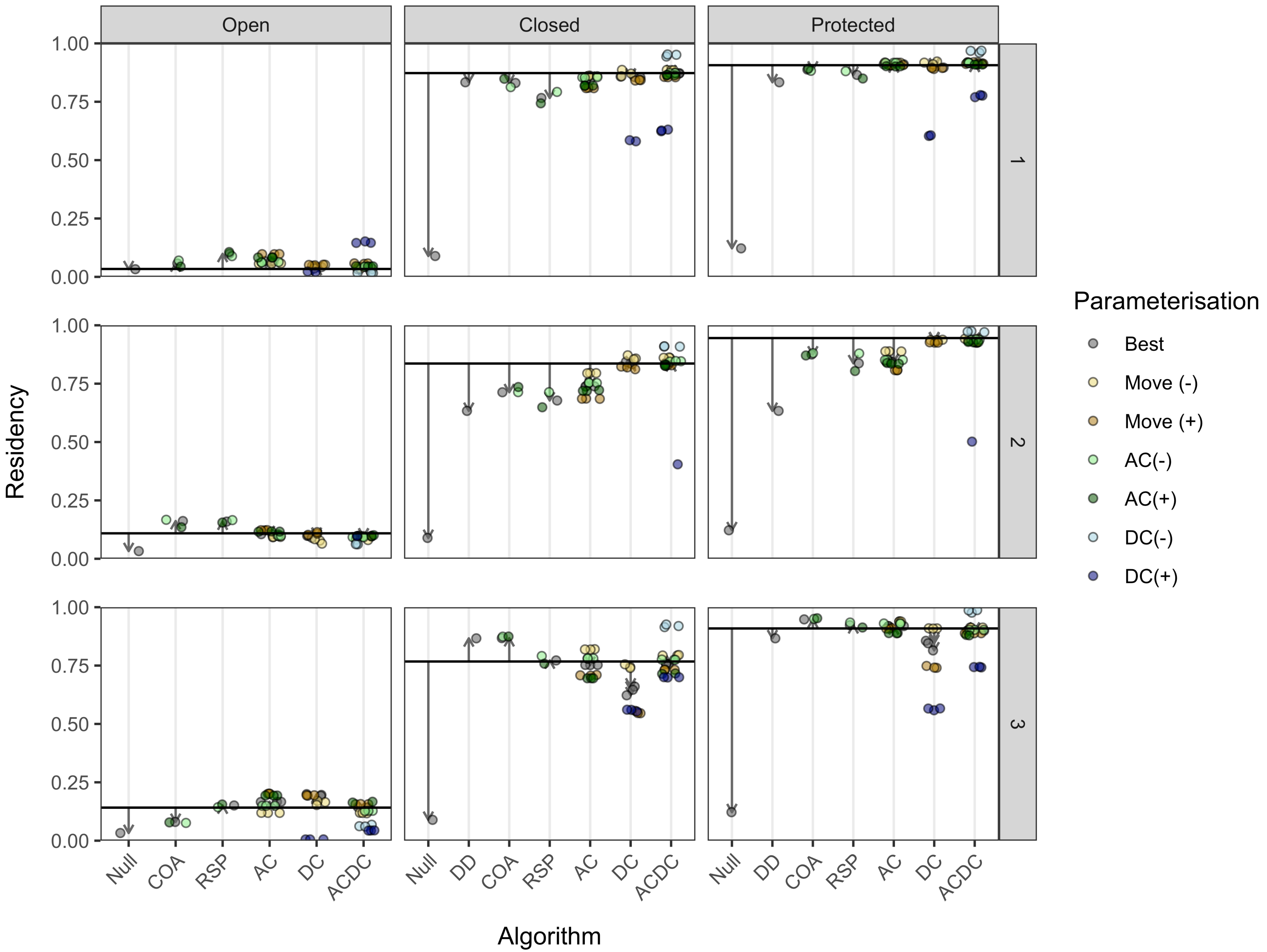
Open
Closed
Protected
1
2
3
Residency
1.00
0.75
0.50
0.25
0.00
Null
DD
COA
RSP
AC
DC
ACDC
Algorithm
Parameterisation
Best
Move (-)
Move (+)
AC(-)
AC(+)
DC(-)
DC(+)

**Figure S12. Simulation analysis: algorithm sensitivity for estimating residency**. Each panel shows residency estimates from different algorithms in open or closed zones, or the MPA at large, for a given path (rows) and selected algorithm implementations (columns). Residency is estimated as the proportion of time spent in each area. The black horizontal line is the proportion of time steps the simulated path spent in each area. The null model corresponds to the proportion of the study area in each area. Detection days is the proportion of days with detections at receiver(s) (all of which are in closed zones). For heuristic (COA and RSP) and particle (AC, DC, ACDC) algorithms, residency was estimated (a) as the proportion of a UD's volume or (b) the proportion of particles, in each area, respectively. Particle (AC, DC and ACDC) algorithm implementations included a best (data-generating) parameterisation alongside under (-) or over (+) estimation of the movement (Move), acoustic (AC) and depth (DC) observation model parameters, with other components held constant at data-generating values. Particle algorithms were implemented independently three times to examine reproducibility (and three points are shown for each setting, except in the case of convergence failures). The two heuristic (COA and RSP) algorithms were implemented with optimal values and restricted/flexible parameterisations that are nominally included in the AC(-) and AC(+) categories, since these algorithms focus exclusively on acoustic detections. These algorithms are deterministic and only a single point is shown for the relevant settings. Vertical arrows point to the best residency estimate in each cluster. The spread of residency estimates around this point indicates the sensitivity, and reproducibility, of residency estimates under the different algorithm parameterisations.

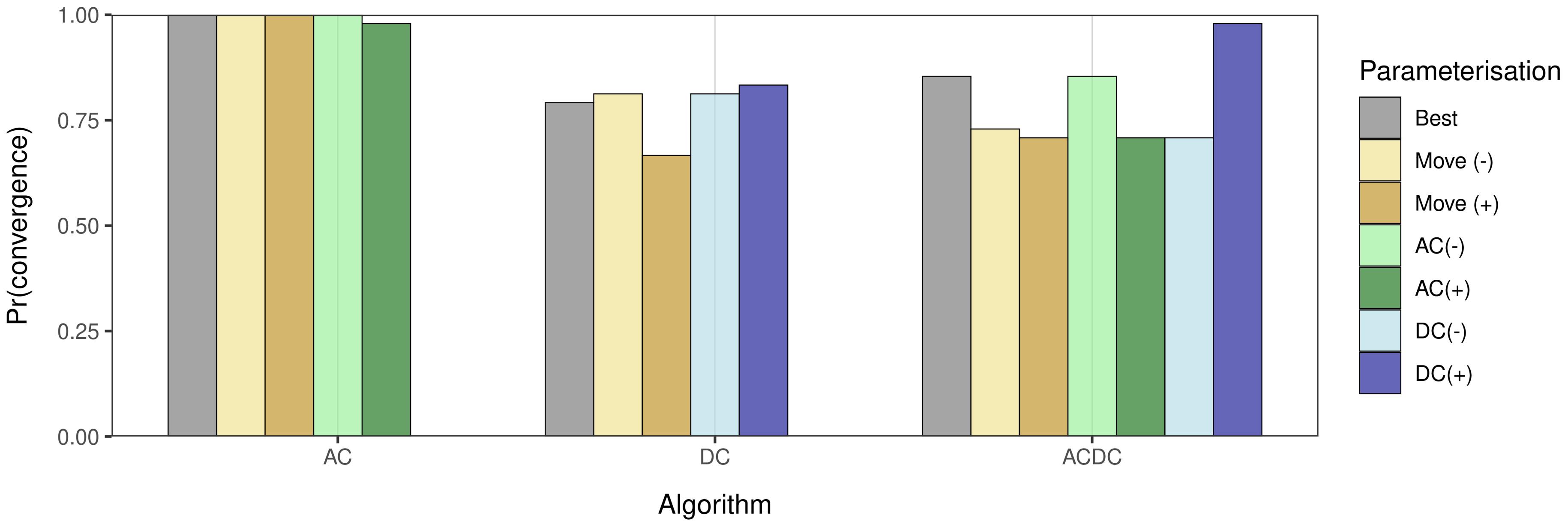


**Figure S13. Real-world analysis: particle algorithm convergence properties.** For each particle algorithm, the proportion of algorithm runs that converged is shown for the five algorithm parameterisations[55].

[55] These are explained in Figures S12–13 with respect to simulation analyses.

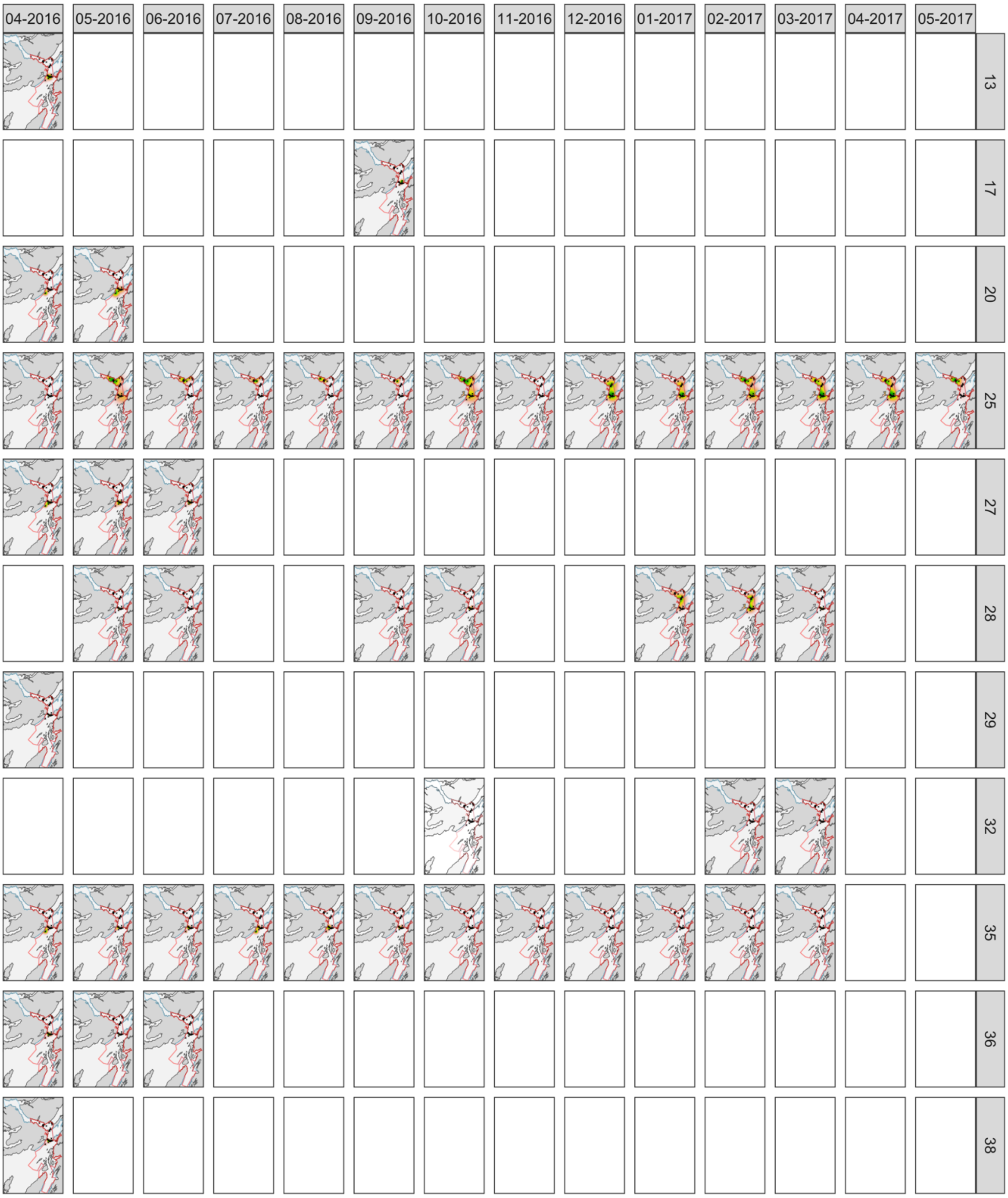


**Figure S14. Real-world analysis: COA algorithm maps from the main analysis.** Each panel is the UD for a given individual (row) in a given month (column). As in earlier figures, crosses mark receivers. The colour scale represents probability density and is shown between zero (white) and the maximum density (green) on each panel. (In some panels, UDs from the COA algorithm are so concentrated they are 'hidden' by receivers.) For clarity, maps are only shown within the extent of the MPA, as delineated by the blue and red polygons (following earlier figures). These enclose protected zones open and closed to fisheries, respectively. Transparent panels indicate convergence failures (if applicable). Blank panels indicate individual/month combinations for which data were not collected or modelled (see Figure S2).

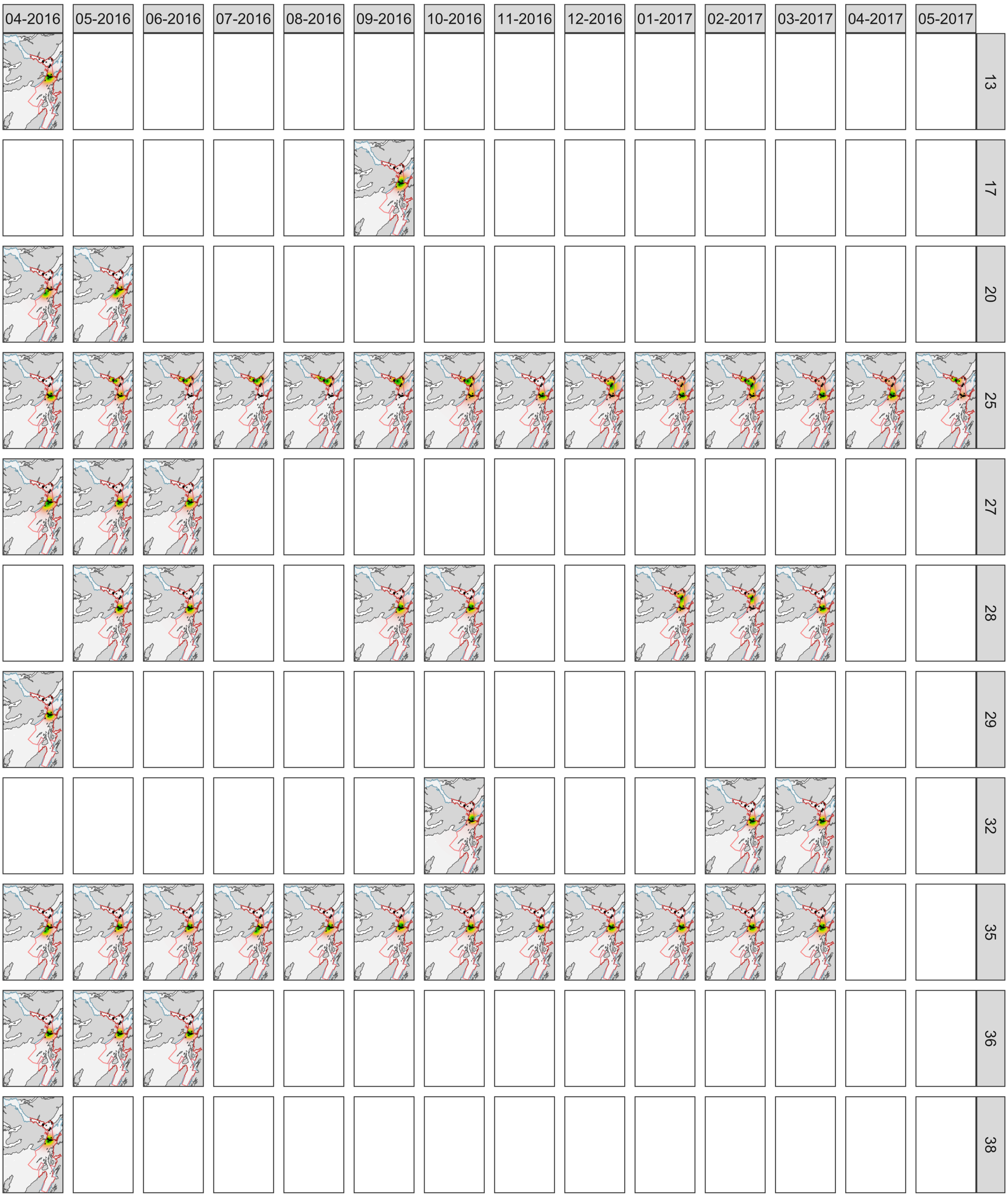


**Figure S15. Real-world analysis: RSP algorithm maps from the main analysis, following Figure S14.**

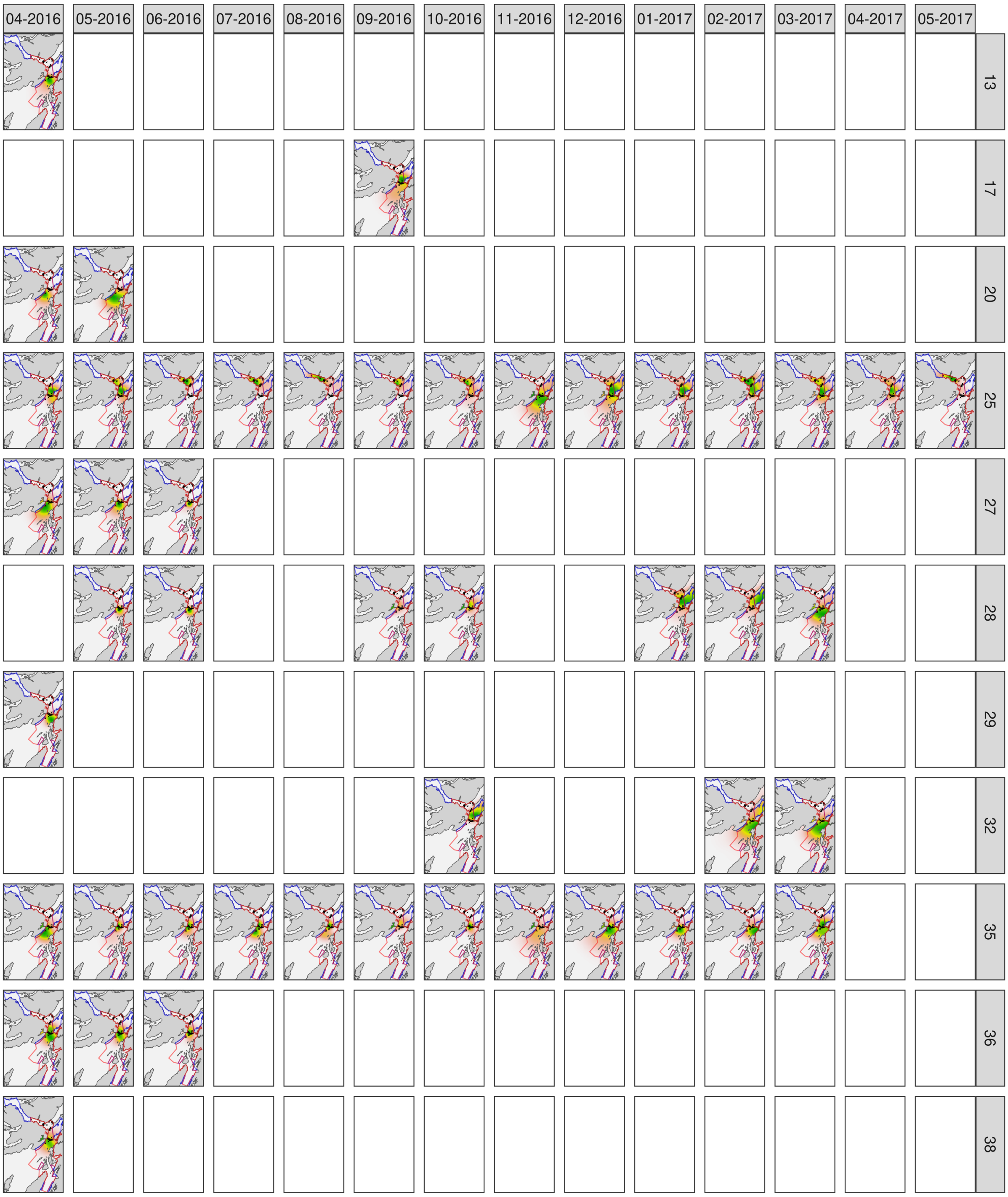


**Figure S16. Real-world analysis: AC algorithm maps from the main analysis, following Figure S14.**

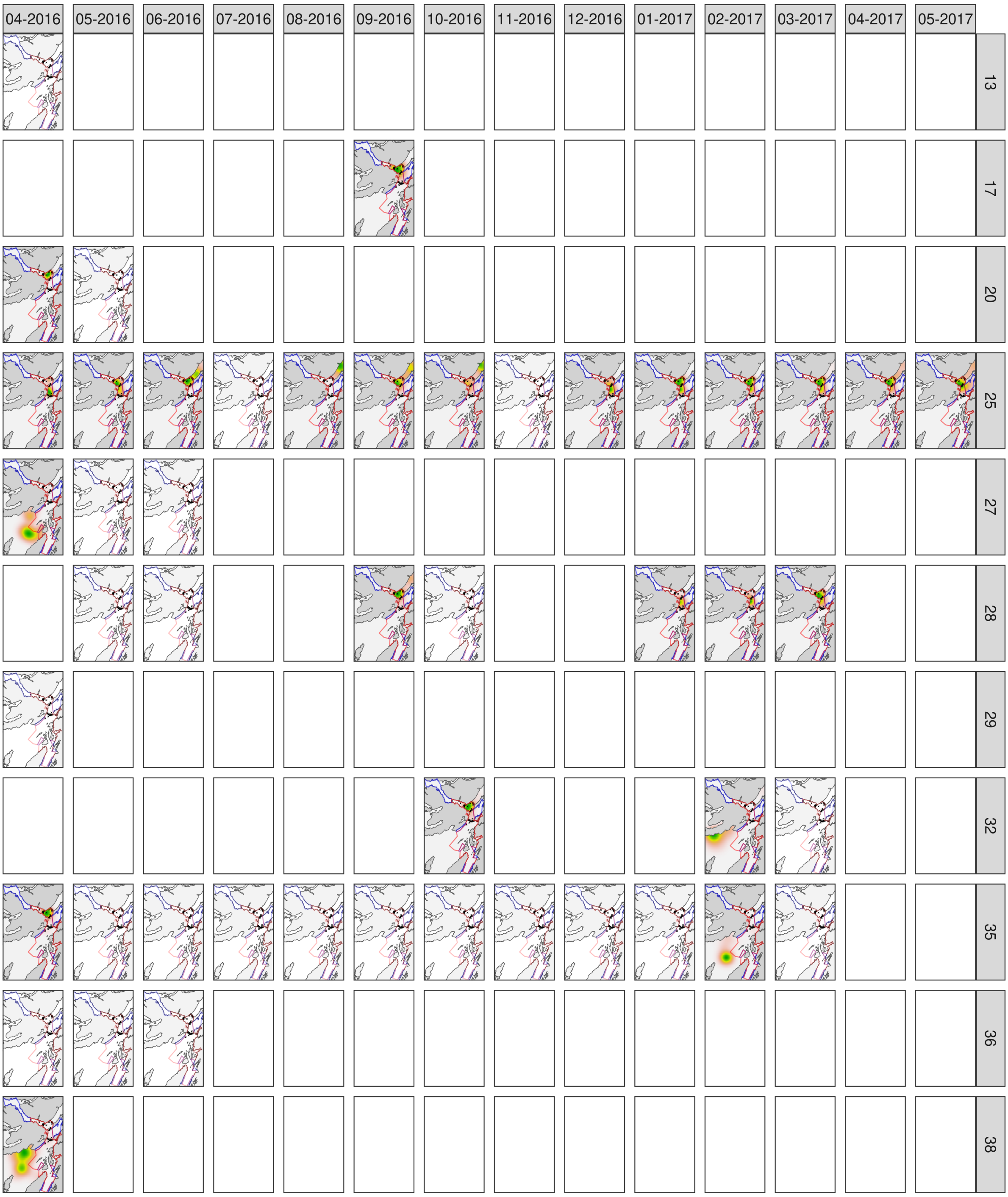


**Figure S17. Real-world analysis: DC algorithm maps from the main analysis, following Figure S14.**

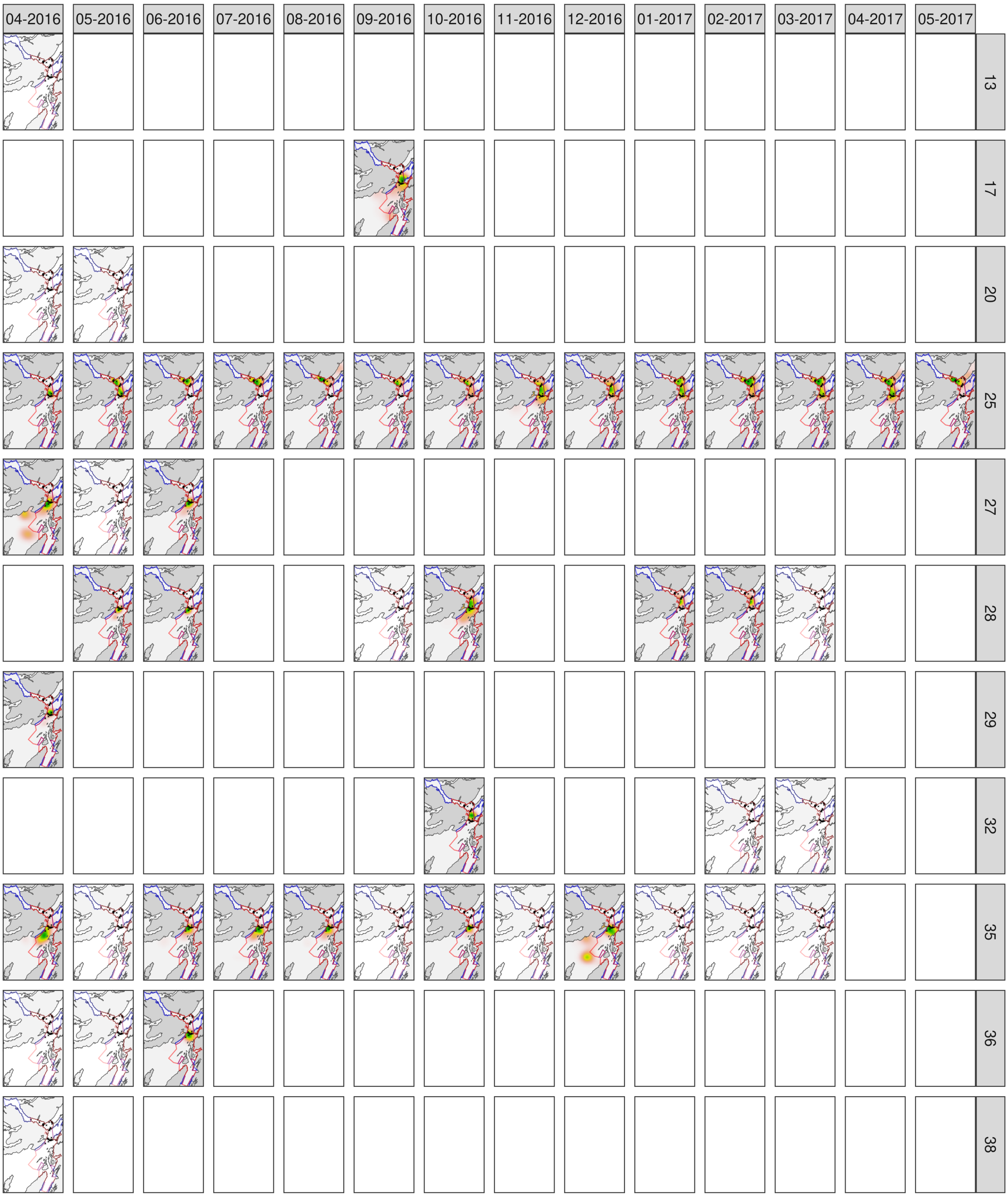


**Figure S18. Real-world analysis: ACDC algorithm maps from the main analysis, following Figure S14.**

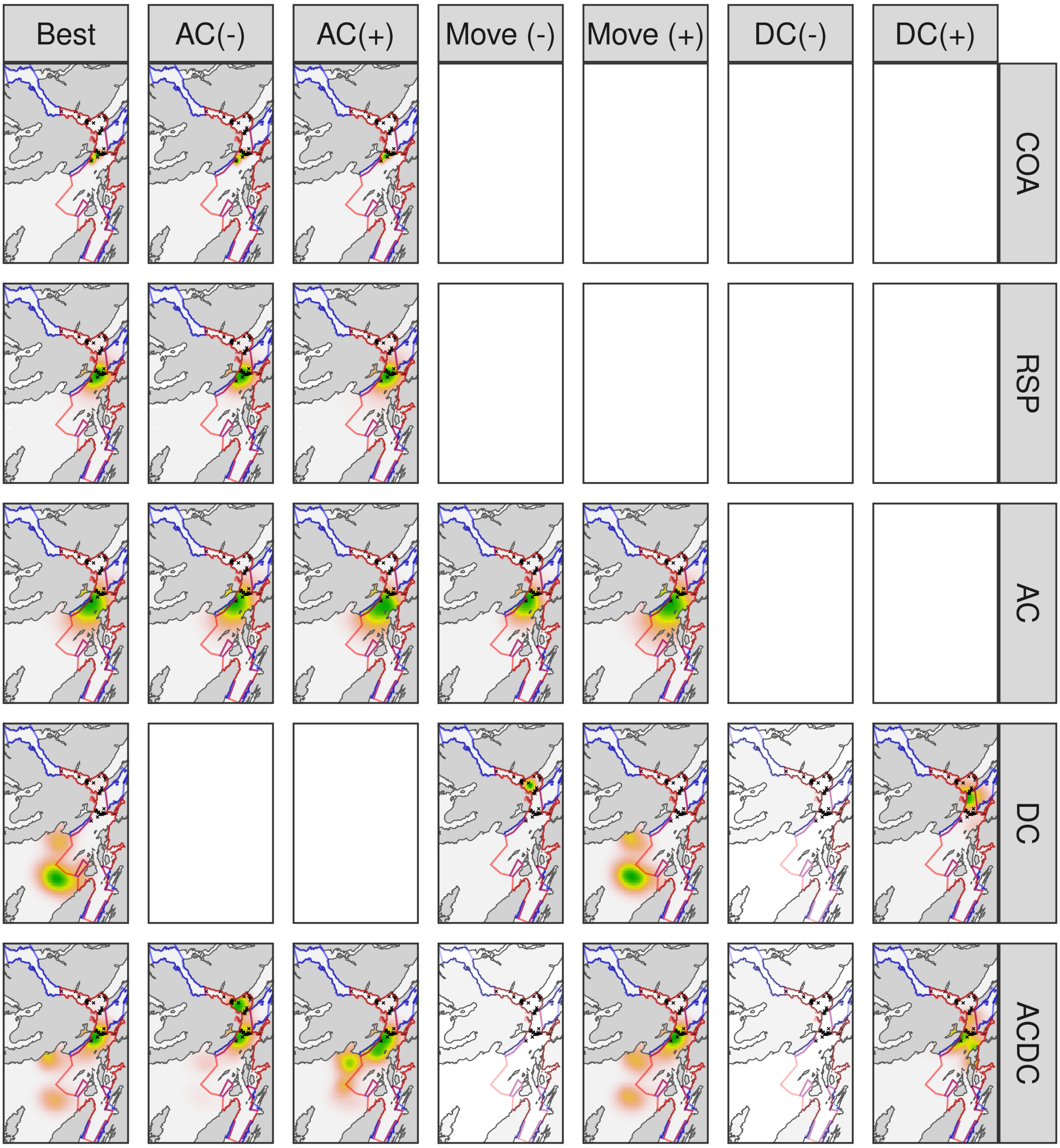


**Figure S19. Real-world analysis: sensitivity in maps of space use.** Each panel shows the UD for an example individual (27) in an example month (April 2016) according to a particular algorithm (row) and parameterisation (column)[56].

[56] Figure properties follow earlier figures. Particle (AC, DC and ACDC) algorithm implementations included a best (data-generating) parameterisation alongside restrictive (-) or flexible (+) settings for the movement (Move), acoustic (AC) and depth (DC) observation models, with other components held constant at data-generating values. The two heuristic (COA and RSP) algorithms were implemented with optimal values and restricted/flexible parameterisations that are nominally included in the AC(-) and AC(+) panel columns, since these algorithms focus exclusively on acoustic detections. Transparent panels indicate convergence failures. Blank panels indicate nonsensical algorithm–parameter combinations.

#### Supporting tables

**Table S1. Bathymetric data sources.** For each region, the bathymetry datasets are listed. For the study system, the base dataset was sourced from Howe et al. (2014). In selected regions, gaps in this dataset were filled with additional datasets sourced from Admiralty's Seabed Mapping Service. The Scotland-wide dataset was obtained from Digimap. Data assembly is described in Supporting Information §1.

| Region | Dataset |
|---|---|
| Study system, Firth of Lorn | Howe et al. (2014) |
| Study system, Coll and Tiree | 2009 HI1257 The Isles of Eigg and Muck Blk4 2m SB |
| | 2009 HI1297 Western App to Small Isles Blk1 2m SB |
| | 2009 HI1297 Western App to Small Isles Blk1 4m SB |
| | 2009 HI1297 Western App to Small Isles Blk1 8m SB |
| | 2009 HI1297 Western App to Small Isles Blk2 2m SB |
| | 2009 HI1297 Western App to Small Isles Blk2 4m SB |
| | 2009 HI1297 Western App to Small Isles Blk2 8m SB |
| | 2009 HI1297 Western App to Small Isles Blk3 2m SB |
| | 2009 HI1297 Western App to Small Isles Blk3 4m SB |
| | 2009 HI1297 Western App to Small Isles Blk3 8m SB |
| | 2009 HI1297 Western App to Small Isles Blk4 2m SB |
| | 2009 HI1297 Western App to Small Isles Blk4 4m SB |
| | 2009 HI1297 Western App to Small Isles Blk4 8m SB |
| | 2010 HI1297 Western App to Small Isles Blk4 Addendum 2m SB |
| | 2010 HI1297 Western App to Small Isles Blk4 Addendum 4m SB |
| | 2012 HI1297 Inner App to Small Isles Blk6 2m SB |
| | 2012 HI1297 Inner App to Small Isles Blk6 4m SB |
| | 2012 HI1297 Inner App to Small Isles Blk6 8m SB |
| | 2012 HI1297 Pt 2 Inner App to Small Isles Blk2 2m SB |
| | 2012 HI1297 Pt 2 Inner App to Small Isles Blk2 4m SB |
| | 2012 HI1297 Pt 2 Inner App to Small Isles Blk2 8m SB |
| | 2012 HI1297 Pt 2 Inner App to Small Isles Blk4 2m SB |
| | 2012 HI1297 Pt 2 Inner App to Small Isles Blk4 4m SB |
| | 2012 HI1298 Passage of Tiree Blk B 2m SB |
| Study system, Islay | 2010 HI1329 Sound of Islay Blk1 2m SB |
| | 2010 HI1329 Sound of Islay Blk2 2m SB |
| | 2012 2014-249146 South West Islay 2m SB |
| | 2012 HI1362 Outer Approaches to Firth of Lorne Blk E 2m SB |
| | 2012 HI1362 Outer Approaches to Firth of Lorne Blk E 4m SB |
| | 2013 HI1370 Approaches to Sound of Jura 2m SB |
| | 2013 HI1370 Approaches to Sound of Jura 4m SB |
| | 2013 HI1371 Sound of Jura 2m SB |
| | 2013 HI1371 Sound of Jura 4m SB |
| Study system, Loch Creran | 2017 2021-230548 Loch Creran 2m SDTP |
| Study system, Loch Etive | 2014 HI1441 Loch Etive 2m SDTP |
| | 2014 HI1441 Loch Etive 4m SDTP |
| | 2014 HI1441 Loch Etive 8m SDTP |
| | 2012 HI1373 Loch Linnhe Blk A 2m SB |

Supporting information, figures and tables

| Region | Dataset |
|---|---|
| Study system, Loch Linnhe | 2012 HI1373 Loch Linnhe Blk A 4m SB |
| | 2012 HI1373 Loch Linnhe Blk A 8m SB |
| | 2012 HI1373 Loch Linnhe Blk A 12m SB |
| | 2012 HI1373 Loch Linnhe Blk B 2m SB |
| | 2012 HI1373 Loch Linnhe Blk B 4m SB |
| | 2012 HI1373 Loch Linnhe Blk B 8m SB |
| | 2012 HI1373 Loch Linnhe Blk C 2m SB |
| | 2012 HI1373 Loch Linnhe Blk C 4m SB |
| | 2012 HI1373 Loch Linnhe Blk C 8m SB |
| | 2012 HI1373 Loch Linnhe Blk C 12m SB |
| Study system, Lochgilphead | 2015 2015-209463 Claonaig Slipway |
| | 2015 2015-209463 Lochranza |
| | 2016 2016-298157 Port Crannaich 2m |
| | 2016 2017-310993 Loch Fyne Otter Spit |
| | 2016 2020-191663 Loch Fyne Wreck and Beaufort Rock 2m SDTP |
| | 2020 2021-262711 Carradale Harbour |
| | 2021 HI1680 Kilbrannan Sound 0-40m 2m SDTP |
| | 2021 HI1680 Kilbrannan Sound 36-152m 4m SDTP |
| | 2022 2022-067840 Claonaig Slipway |
| | 2022 2022-067840 Lochranza |
| | 2023 2023-150089 Claonaig Slipway |
| | 2023 2023-129344 Carradale Harbour |
| Scotland | marine-dem-6-sec_3965365 |

**Table S2. Summary of the workflow of simulation and real-world analyses.**

**Workflow summary**

**Simulation experiments**

***Examine algorithm behaviour (relative performance, sensitivity and reproducibility) in our study system and inform real-world analyses***

**A1. Performance evaluation**

***Compare the performance of heuristic and particle algorithms for reconstructing patterns of space use and estimating residency, using 100 simulated paths/observational datasets***

**A1.1. Algorithm parameterisation**

**A1.1.1. *Heuristic algorithms.*** Select optimal parameters for the COA ($\Delta T$) and RSP (`er.ad`) algorithms from profiles of the mean absolute error (ME), between UDs for the simulated paths and UDs reconstructed with different parameter values (Figure S3).

**A1.1.2. *Particle algorithms.*** Use data-generating (i.e., best-guess) parameters (chosen for flapper skate, based on biological expertise).

**A1.2. Algorithm implementation.** Implement the algorithms with optimal or best-guess parameters and validate convergence (Figure S4).

**A1.3. Algorithm performance**

**A1.3.1. *Maps of space use: visual evaluation.*** Visualise maps of space use for the first three paths (Figures S5–7).

**A1.3.2. *Maps of space use: quantitative evaluation.*** Compute the distribution of ME (from the UD for each simulated path and the reconstructed UD), across all paths by algorithm (Figure 2).

**A1.3.3. *Residency*.** Compute the distribution of residency error (RE) as the difference between estimated and true residency, across all paths by algorithm, in open/closed zones and the entire MPA (Figures 2 and S8).

**A2. Sensitivity and reproducibility evaluation**

***Objective: compare algorithm sensitivity to parameter choices for maps of space use and residency estimates, using the first three simulated paths/observational datasets***

**A2.1. Algorithm implementation.** Implement the algorithms with restrictive and flexible settings. For COAs and RSPs, restrict/inflate $\Delta T \in \{2 \text{ days}, 1 \text{ day}, 3 \text{ days}\}$ and $er.ad \in \{500, 250, 750\}$. For particle algorithms, restrict/inflate the movement model, acoustic observation model and/or depth observation model parameters, while holding other parameters constant. Repeat particle algorithms independently three times to examine reproducibility.

**A2.2. Algorithm sensitivity and reproducibility**

**A2.2.1. *Maps of space use: visual evaluation.*** Compare maps of space use using optimal/best-guess, restrictive and flexible parameters, and across multiple iterations of the particle algorithms (Figures S5–6 and S9–10).

**A2.2.2. *Maps of space use: quantitative evaluation.*** Compute ME for each algorithm parameterisation/iteration (Figure S11).

| Workflow summary |
|---|
| **A2.2.3.** ***Residency.*** Compute RE for each algorithm parameterisation/iteration (Figure S12). |
| **Real-world analyses**<br>***Reconstruct patterns of space use and residency of flapper skate in the Loch Sunart to the Sound of Jura MPA***<br><br>**A3. Main analyses**<br>***Reconstruct patterns of space use and estimate residency via heuristic and particle algorithms for each individual and month with sufficient observations, using optimal/best-guess algorithm parameterisations***<br><br>**A3.1. Algorithm implementation.** Implement the algorithms with optimal/best-guess parameters (as in simulations) and validate convergence (Figure S13).<br><br>**A3.2. Analyses**<br>**A3.2.1.** ***Maps of space use: visual evaluation.*** Reconstruct and visualise patterns of space use for each time series using best-guess algorithm parameterisations (Figures 3 and S14–18).<br>**A3.2.2.** ***Maps of space use: summary.*** Reconstruct an overall map of space use and visually assess the overlap with presence records from angling (Figure 4).<br>**A3.2.3.** ***Residency***. Estimate residency within open/closed zones and the entire MPA (Figure 5).<br><br>**A4. Sensitivity analyses**<br>***Examine sensitivity to parameter choices for maps of space use and residency estimates***<br><br>**A4.1. Algorithm implementation.** Implement the algorithms with restrictive and flexible parameters (as in simulations) and validate convergence (Figure S13).<br><br>**A4.2. Analyses**<br>**A4.2.1.** ***Maps of space use.*** Reconstruct patterns of space use and visually compare sensitivity to algorithm parameterisation (Figure S19).<br>**A4.2.2.** ***Residency***. Estimate residency in open/closed zones and the entire MPA, and visually compare sensitivity to algorithm parameterisation (Figure 5). |

**Table S3. Summary of notation.** The essential notation in the Main Text and the Supporting Information is listed, ordered alphabetically. Example references to Supporting Information equations are provided. Notation broadly follows Lavender et al. (2024a) where further details are provided.

| Notation | Description |
|---|---|
| $A$ | A set/subset of locations in the study area compatible with the initial observation(s). See eqn 3. |
| $\alpha$ | A linear coefficient in a logistic, distance-decaying, detection-probability model. See eqn 12. |
| behaviour | The behavioural state of the individual: behaviour $=0$ (low activity) or behaviour $=1$ (active). See eqn 9. |
| $b(\boldsymbol{s})$ | The bathymetric depth (m) in the location defined by $\boldsymbol{s}$. See eqn 13. |
| $\beta$ | A linear coefficient in a logistic, distance-decaying, detection-probability model. See eqn 12. |
| $d$ | The movement step length (m). See eqn 4. |
| $\delta_{I,I_{i,t}}$ | The Kronecker delta. This evaluates to one if the grid cell of coordinate $i$ at time $t$ ($I_{i,t}$) equals grid cell $I$ or 0 otherwise. See eqn 23. |
| $f(\cdot)$ | A probability density function. See eqn 1. |
| $f(\boldsymbol{s}_{1:T})$ | The prior probability density (i.e., the movement process), which is modelled as a discrete-time Markovian process. See eqn 1. |
| $f(\boldsymbol{s}_{t=1})$ | The probability density of the individual's initial state. See eqn 2. |
| $f(\boldsymbol{s}_t \mid \boldsymbol{s}_{t-1})$ | The probability density of movement from the individual's state at time $t-1$ to the location at time $t$; i.e., the movement model. See eqn 2. |
| $f(\boldsymbol{s}_t \mid \boldsymbol{y}_{1:T})$ | The full marginal distribution of the individual's state at a given time ($\boldsymbol{s}_t$) given all of the data ($\boldsymbol{y}_{1:T}$). This is approximated by particle smoothing. |
| $f(\boldsymbol{s}_{1:T} \mid \boldsymbol{y}_{1:T})$ | The joint probability distribution of the individual's states ($\boldsymbol{s}_{1:T}$) and all data ($\boldsymbol{y}_{1:T}$). See eqn 1. |
| $f(\boldsymbol{y}_t \mid \boldsymbol{s}_t)$ | The probability density of the observations at a given time ($\boldsymbol{y}_t$) given the individual's state at that time ($\boldsymbol{s}_t$); i.e., the likelihood. |
| $f(\boldsymbol{y}_{1:T} \mid \boldsymbol{s}_{1:T})$ | The joint probability density of all observations ($\boldsymbol{y}_{1:T}$) and all states ($\boldsymbol{s}_{1:T}$); i.e., the joint likelihood. See eqn 1. |
| $f_d(d)$ | The probability density of step lengths ($d$). See eqn 5. |
| $f_{\Delta\phi}(\Delta\phi)$ | The probability density of turning angles ($\Delta\phi$). See eqn 5. |
| $\phi$ | A heading (in radians). See eqn 4. |
| $\gamma$ | The maximum detection range (m); that is, the distance from a receiver beyond which the detection of an acoustic transmission is assumed impossible. See eqn 12. |
| $h(\boldsymbol{s}_t, \boldsymbol{r}_k)$ | The Euclidean distance between the transmitter, at the location defined by $\boldsymbol{s}_t$, and receiver $k$ (at $\boldsymbol{r}_k = (s_{x;k}, s_{y;k})$). |
| $I$ | An index over grid cells. See eqn 23. |
| $J$ | The Jacobian. See eqn 7. |
| $k_1$, $k_2$ | The location parameters of the (truncated) Cauchy distributions, used to model movement step lengths during 'low activity' ($k_1$) and 'active' ($k_2$) behavioural states. See eqn 9. We also separately use $k$ |

| Notation | Description |
|---|---|
| | as an index over operational receivers (see eqn 11) and $k_i$ to represent the number of time steps over which a behaviour is maintained (see eqn 26). |
| $\text{lim}_{\text{shallow}}, \text{lim}_{\text{deep}}$ | Shallow and deep truncation limits in the depth observation model. See eqn 13. |
| mobility | The maximum moveable distance (m) between two consecutive time steps. (This is the upper truncation parameter of the truncated Cauchy distribution used to model movement step lengths.) See eqn 9. |
| $p_{k,t}(\boldsymbol{s}_t)$ | The probability of a detection at receiver $k$ at time $t$, given an acoustic transmission from the location defined by $\boldsymbol{s}_t$. See eqn 11. |
| $P_I$ | Probability-of-use for grid cell $I$; that is, the probability that an individual is located in that cell at a randomly chosen time. See eqn 23. |
| $\boldsymbol{r}_k = (s_{x;k}, s_{y;k})$ | Receiver $k$'s two-dimensional location. The expression $\boldsymbol{r}_k = (s_{x;k}, s_{y;k})$ denotes that $\boldsymbol{r}_k$ is a two-dimensional vector with x ($s_{x;k}$) and y ($s_{y;k}$) coordinates. See eqn 12. |
| $\boldsymbol{s}_t$ | The individual's state $\boldsymbol{s}_t = (s_{x,t},\ s_{y,t},\ \phi_t,\ \text{behaviour}_t)$. This comprises the individual's two-dimensional location $(s_{x,t},\ s_{y,t})$, the heading $(\phi_t)$ and the behaviour. See eqn 1. |
| $\sigma_{\text{depth}}$ | The standard deviation of the (truncated) Gaussian distribution of the depth observation model. See eqn 13. |
| $\sigma_{\Delta\phi}$ | The standard deviation of the Gaussian model of turning angle. See eqn 10. |
| $t$ | The time step index $(1, 2, \dots, T)$. See eqn 1. |
| $T$ | The final time step. See eqn 1. |
| $\theta_1, \theta_2$ | The scale parameters of the (truncated) Cauchy distribution used to model movement step lengths during 'low activity' $(\theta_1)$ and 'active' $(\theta_2)$ behavioural states. See eqn 9. |
| $\mu_{\text{depth}}$ | The mean of the (truncated) Gaussian distribution of the depth observation model. See eqn 13. |
| $w_{i,t}$ | The normalised weight of the $i^{\text{th}}$ pair of coordinates at time $t$. See eqn 23. |
| $\boldsymbol{y}$ | A dataset of observations. We consider a combined dataset comprising acoustic and archival (depth) observations; i.e., $\boldsymbol{y} = \{\boldsymbol{y}^{(A)}, \boldsymbol{y}^{(D)}\}$. See eqn 1. |
| $\boldsymbol{y}^{(A)}$ | An $M \times T$ matrix of acoustic observations, which comprise detections (1) and non-detections (0) at each receiver $k$; i.e., $y_{k,t}^{(A)} \in \{0, 1\}$. A column-vector of acoustic observations at time $t$ is denoted $\boldsymbol{y}_t^{(A)}$. See eqn 11. |
| $\boldsymbol{y}^{(D)}$ | A row vector of depth (m) observations. A single depth observation at time $t$ is denoted $y_t^{(D)}$. See eqn 13. |
| $\frac{1}{z(\boldsymbol{s}_{t-1})}$ | A normalisation constant required to evaluate the probability density of a movement from location $\boldsymbol{s}_{t-1} \rightarrow \boldsymbol{s}_t$. See eqn 8. |

**Table S4. Parameter values for algorithm implementations.** For each algorithm, component sub-model, parameter and parameter setting (1, 2, 3), parameter values are shown. Parameter settings 1, 2 and 3, correspond towards the optimal/best-guess, restricted and flexible parameterisations. In analyses A1 and A3, optimal/best-guess algorithm parameterisations were used to evaluate algorithm performance and for the main real-world analyses (Table S2). In analyses A2 and A4, the different parameter settings were used to analyse algorithm sensitivity (Table S2). The COA algorithm depends only on time interval over which COAs are calculated ($\Delta T$). The RSP algorithm depends on a suite of tuning parameters. We trialled three settings for the `er.ad` parameter, which tunes the rate at which uncertainty increases with distance from receivers; the detection range was set to $\gamma = 3000$ m (as in best-guess particle algorithms); and other parameters were set to default values. The particle algorithms (AC, DC and ACDC) depend on movement (Move) and observation (acoustic [AC] and/or depth [DC]) model parameters: $k_1, k_2, \theta_1, \theta_2$ and mobility are the parameters of a truncated, two-state Cauchy model of step length; $\sigma_{\Delta\phi}$ is the standard deviation of a Gaussian model of turning angle; $\alpha$, $\beta$, $\gamma$ are the parameters of a truncated logistic distance-decaying acoustic detection-probability model; and $\mu_{\text{depth}}$, $\sigma_{\text{depth}}$, $\text{lim}_{\text{shallow}}$ and $\text{lim}_{\text{deep}}$ are the parameters of a truncated Gaussian depth observation model. For details, see Supporting Information §4. For a visualisation of particle algorithm sub-models, see Figure S1.

| **Algorithm** | **Sub-model** | **Parameter** | **Setting** | **Value** |
|---|---|---|---|---|
| COA | - | $\Delta T$ | 1 | 2 days |
| | | | 2 | 1 day |
| | | | 3 | 3 days |
| RSP | - | $\text{er.ad}$ | 1 | 500.0 |
| | | | 2 | 250.0 |
| | | | 3 | 750.0 |
| AC | Movement (Move) | $\begin{cases}(k_1, \theta_1, \text{mobility}, \sigma_{\Delta\phi})\\(k_2, \theta_2, \text{mobility}, \sigma_{\Delta\phi})\end{cases}$ | 1 | $\begin{cases}(0.0, 5.0, 1095.0, 1.5)\\(5.0, 100.0, 1095.0, 1.5)\end{cases}$ |
| | | | 2 | $\begin{cases}(0.0, 5.0, 985.5, 1.5)\\(2.5, 50.0, 985.5, 1.5)\end{cases}$ |
| | | | 3 | $\begin{cases}(0.0, 5.0, 1204.5, 1.5)\\(7.5, 150.0, 1204.5, 1.5)\end{cases}$ |
| | Acoustic (AC) | $(\alpha, \beta, \gamma)$ | 1 | $(4.0, -0.0094, 3000.0)$ |
| | | | 2 | $(3.0, -0.01175, 2250.0)$ |
| | | | 3 | $(5.0, -0.00705, 3750.0)$ |
| DC | Movement (Move) | | 1 | $\begin{cases}(0.0, 5.0, 1095.0, 1.5)\\(5.0, 100.0, 1095.0, 1.5)\end{cases}$ |

| Algorithm | Sub-model | Parameter | Setting | Value |
|---|---|---|---|---|
| | | $\begin{cases}(k_1, \theta_1, \text{mobility}, \sigma_{\Delta\phi}) \\ (k_2, \theta_2, \text{mobility}, \sigma_{\Delta\phi})\end{cases}$ | 2 | $\begin{cases}(0.0, 5.0, 985.5, 1.5) \\ (2.5, 50.0, 985.5, 1.5)\end{cases}$ |
| | | | 3 | $\begin{cases}(0.0, 5.0, 1204.5, 1.5) \\ (7.5, 150.0, 1204.5, 1.5)\end{cases}$ |
| | Depth (DC) | $(\mu_{\text{depth}}, \sigma_{\text{depth}}, \text{lim}_{\text{shallow}}, \text{lim}_{\text{deep}})$ | 1 | $(b(\boldsymbol{s}), 100.0, 0.0, 350.0)$ |
| | | | 2 | $(b(\boldsymbol{s}), 50.0, 0.0, 350.0)$ |
| | | | 3 | $(b(\boldsymbol{s}), 150.0, 0.0, 350.0)$ |
| ACDC | Movement (Move) | $\begin{cases}(k_1, \theta_1, \text{mobility}, \sigma_{\Delta\phi}) \\ (k_2, \theta_2, \text{mobility}, \sigma_{\Delta\phi})\end{cases}$ | 1 | $\begin{cases}(0.0, 5.0, 1095.0, 1.5) \\ (5.0, 100.0, 1095.0, 1.5)\end{cases}$ |
| | | | 2 | $\begin{cases}(0.0, 5.0, 985.5, 1.5) \\ (2.5, 50.0, 985.5, 1.5)\end{cases}$ |
| | | | 3 | $\begin{cases}(0.0, 5.0, 1204.5, 1.5) \\ (7.5, 150.0, 1204.5, 1.5)\end{cases}$ |
| | Acoustic (AC) | $(\alpha, \beta, \gamma)$ | 1 | $(4.0, -0.0094, 3000.0)$ |
| | | | 2 | $(3.0, -0.01175, 2250.0)$ |
| | | | 3 | $(5.0, -0.00705, 3750.0)$ |
| | Depth (DC) | $(\mu_{\text{depth}}, \sigma_{\text{depth}}, \text{lim}_{\text{shallow}}, \text{lim}_{\text{deep}})$ | 1 | $(b(\boldsymbol{s}), 100.0, 0.0, 350.0)$ |
| | | | 2 | $(b(\boldsymbol{s}), 50.0, 0.0, 350.0)$ |
| | | | 3 | $(b(\boldsymbol{s}), 150.0, 0.0, 350.0)$ |

**Table S5. Summary of modelled flapper skate.** For each individual, the acoustic and archival (depth) tag identifiers are provided for reference to previous publications (e.g., Lavender et al., 2021a, 2021b). Individual sex, maturation status and residency labels assigned by Lavender et al. (2021a) are also provided. Residency labels comprise the residency category (non [N], short-term [S] or long-term [L] resident) and the total number of days with detections. (For details, see Lavender et al. (2021a).) In the present study, only a subset of ($N$) months with sufficient data (listed) were analysed for most individuals.

| Individual | Acoustic | Archival | Sex | Maturity | Resident | Months |
|---|---|---|---|---|---|---|
| 13 | 249 | 1558 | F | Immature | L185 | (1) 04-2016 |
| 17 | 244 | 1536 | M | Mature | N38 | (1) 09-2016 |
| 20 | 242 | 1538 | F | Immature | S24 | (2) 04-2016, 05-2016 |
| 25 | 555 | 1547 | F | Mature | L256 | (14) 04-2016, 05-2016, 06-2016, 07-2016, 08-2016, 09-2016, 10-2016, 11-2016, 12-2016, 01-2017, 02-2017, 03-2017, 04-2017, 05-2016 |
| 27 | 563 | 1520 | M | Mature | S64 | (3) 04-2016, 05-2016, 06-2016 |
| 28 | 560 | 1522 | F | Immature | S150 | (7) 05-2016, 06-2016, 09-2016, 10-2016, 01-2016, 02-2016, 03-2016 |
| 29 | 542 | 1523 | M | Mature | N34 | (1) 04-2016 |
| 32 | 549 | 1548 | M | Mature | S38 | (3) 10-2016, 02-2017, 03-2017 |
| 35 | 540 | 1509 | F | Immature | L194 | (12) 04-2016, 05-2016, 06-2016, 07-2016, 08-2016, 09-2016, 10-2016, 11-2016, 12-2016, 01-2016, 02-2017, 03-2017 |
| 36 | 543 | 1512 | F | Mature | S130 | (3) 04-2016, 05-2016, 06-2016 |
| 38 | 547 | 1507 | F | Mature | S67 | (1) 04-2016 |
